# 31,600-year-old human virus genomes support a Pleistocene origin for common childhood infections

**DOI:** 10.1101/2021.06.28.450199

**Authors:** Sofie Holtsmark Nielsen, Lucy van Dorp, Charlotte J. Houldcroft, Anders G. Pedersen, Morten E. Allentoft, Lasse Vinner, Ashot Margaryan, Elena Pavlova, Vyacheslav Chasnyk, Pavel Nikolskiy, Vladimir Pitulko, Ville N. Pimenoff, François Balloux, Martin Sikora

## Abstract

The origins of viral pathogens and the age of their association with humans remains largely elusive. To date, there is no direct evidence about the diversity of viral infections in early modern humans pre-dating the Holocene. We recovered two near-complete genomes (5.2X and 0.7X) of human adenovirus C (HAdV-C), as well as low-coverage genomes from four distinct species of human herpesvirus obtained from two 31,630-year-old milk teeth excavated at Yana, in northeastern Siberia. Phylogenetic analysis of the two HAdV-C genomes suggests an evolutionary origin around 700,000 years ago consistent with a common evolutionary history with hominin hosts. Our findings push back the earliest direct molecular evidence for human viral infections by ∼25,000 years, and demonstrate that viral species causing common childhood viral infections today have been in circulation in humans at least since the Pleistocene.

## Introduction

Hundreds of viruses infect humans today, causing significant morbidity and mortality worldwide. It is of fundamental interest to understand when, where, and how pathogens affecting humans first emerged. Such knowledge can be leveraged for improved surveillance, control and treatment. Evolutionary genomic studies of viral pathogens can yield important insights into these questions, but have so far mainly been conducted using extant or historical samples collected over the last 50-100 years(*1*–*10*). This shallow time-depth of samples along the respective virus phylogenies poses considerable challenges to the inference of the origins and diversification of viral pathogens. Differences in the choices of molecular clock models, unknown rates of substitution and recombination, and unreliable or unavailable external calibration points can all lead to widely differing estimates for the rates of viral evolution and the timing of evolutionary events(*11, 12*). The incorporation of heterochronous samples can address many of these shortcomings but have until recently been restricted to fast-evolving viruses where the time-depth of the analysed samples is sufficiently spread along the phylogeny of interest. Furthermore, analyses of extant virus genomes can only provide estimates for the time to the most recent common ancestor (tMRCA) of the sampled lineages, which can be substantially more recent than the age of the association of a virus with its host whenever some lineages of the pathogen went extinct (*13*).

Recent progress in the recovery and analysis of ancient DNA (aDNA) of pathogens from archaeological remains has transformed the study of infectious disease evolution. Ancient pathogen DNA has been successfully isolated for a number of major human bacterial and viral pathogens, such as *Yersinia pestis*, the causative agent of the plague; *Mycobacterium tuberculosis* and *M. leprae*, causing tuberculosis and leprosy; Hepatitis B virus; and Variola virus, causing smallpox(*14*–*27*). Direct molecular evidence for the spatiotemporal distributions of these pathogens has often pointed to an earlier and more widespread presence of the respective diseases than previously thought. Furthermore, phylogenetic analyses of ancient pathogen genomes have revealed extensive pathogen genetic diversity in the past, involving divergent ancient lineages without any known extant descendants(*14, 19, 24, 25, 27*–*29*). Despite these considerable advances, our current knowledge of ancient pathogen genomics is restricted to the Holocene (the last ∼11,500 years of human evolution), with the oldest human viral genome to date recovered from a ∼7,000-year-old Neolithic individual with Hepatitis B virus infection(*24*). The diversity and genetic makeup of viruses circulating among early modern humans during the Pleistocene thus remain virtually unknown.

Large, double-stranded DNA (dsDNA) viruses such as herpesviruses and adenoviruses are among the major causative agents of common childhood infections. There are over 100 known herpesviruses, nine of which infect humans. These are divided into three subfamilies of herpesviruses, alpha, beta and gamma, and are among the most widespread pathogens known globally. For example, the seroprevalence of human cytomegalovirus (*Human betaherpesvirus 5*) ranges from 45% to 100% depending on the human population(*30*). Primary infection is usually mild or asymptomatic, which is often ascribed to coadaptation of herpesviruses with their hosts(*31*). The virus is excreted in bodily fluids following infection and spreads both vertically and horizontally to new hosts by close contact(*32*).

There are currently seven described species of human adenoviruses (HAdV), A-G, and six genotypes of HAdV-C (*1, 2, 5, 6, 57* and *89*). HAdV-C is a ∼36 kilobase (kb) long linear dsDNA virus, classified by its capsid genes, the penton (L2), hexon (L3) and fiber (L5), which contain the most important neutralizing epitopes (*33*). HAdV usually causes mild self-limiting respiratory infections in children but can be severe in immunosuppressed individuals with the central nervous system (CNS) suggested as a common site of persistent adenovirus infection(*33*). HAdV-C genotypes 1 and 2 are associated with over 50% of adenovirus infections in immunocompromised people and HAdV accounts for 15% of upper respiratory tract infections and 5% of lower respiratory infections in adult and paediatric immunocompetent patients(*34*).

Adenovirus was first isolated in 1953 and in 1962 genotype 12 was the first human virus shown to be oncogenic in animals, but not linked to malignant disease in humans. The virus is used as a model system and is intensively studied. It was the organism in which mRNA splicing was discovered and it is widely used as a vector for gene therapy, experimental vaccines, and recently also approved for different COVID-19 vaccines. However, many aspects of the evolution of HAdV-C remain unclear. For example, some viral genotypes seem to recombine readily within the same species, whereas others appear to be clonal (*5, 34*–*40*). Substitution and recombination rates are also largely unknown. Data on incidence rates is also limited though existing immunity is a barrier to the use of HAdV-5 as an adenoviral vector system(*41*) as HAdV-5 has the highest seroprevalence worldwide. Deciphering the extant diversity and distribution in time of adenoviruses infecting humans and revealing their comprehensive long-term evolution may aid in understanding the determinants of its clinical manifestations and possible limitations in its use in gene therapy and as a recombinant vaccine vector.

The aforementioned dsDNA viruses causing childhood infections are generally characterized by low virulence, which for Herpesviruses leads to latent infections after the resolution of primary infection. As such, they are commonly thought to have evolved in a long-term association and co-divergence with their respective host species, sometimes over millions of years (*42, 43*). However, whether these infections were indeed present in early modern humans, and if so, how they relate to extant viral genetic diversity is unknown. Here, we report ancient viral DNA sequences isolated from two milk teeth excavated at the ∼31,600 year-old archaeological site of Yana RHS(*44*), one of the most remote and extreme environments in Northern Siberia. We recovered partial genomes from four species of human herpesviruses (HHV), as well as two near-complete genomes from different genotypes of HAdV-C. These data represent, to our knowledge, the oldest ancient pathogens isolated from early modern humans, and provide direct evidence for a long-term association of common childhood infections with human hosts since the Pleistocene.

## Results

### Ancient pathogen DNA screening and authentication

The archaeological site of Yana RHS, located at high latitude (∼70° N) in far northeastern Siberia, represents the earliest direct evidence for human occupation in the high Arctic, dated to ∼31,600 ybp(*44*). The only human remains recovered at the site are three fragmented milk teeth, two of which yielded DNA that was sequenced to high genomic coverage to investigate the demographic history of the early humans occupying the site(*45*). The two teeth were found to originate from two different individuals and estimated to have been shed at an age of around 10-12 years. To search for evidence of childhood pathogen infections in the two individuals, we used the metagenomic classifier Kraken(*46*) to perform a metagenomic screening of the ∼12 billion shotgun sequencing reads generated previously. We detected putative ancient pathogen DNA sequences in both individuals, with between 677 and 6,311 reads assigned to two families of dsDNA viruses infecting humans, *Adenoviridae* and *Herpesviridae* (Supplementary Table 1-2). At higher taxonomic resolution, the HAdV-C species within *Adenoviridae* had the most reads assigned in both individuals (Yana1: 6,211 reads; Yana2: 677 reads). For *Herpesviridae*, we found DNA derived from multiple species of HHV present in both individuals. Based on these initial results and availability of sample material, we performed additional targeted enrichment of viral DNA followed by sequencing for individual Yana2, as previously described(*25*).

To authenticate the putative ancient virus DNA, the reads from each sample were first mapped to different sets of reference genomes, using bowtie2 (*47*). This was followed by analyses of aDNA damage patterns (Fig. 1; Fig. S1-6), evenness of genomic coverage (Fig. 1; Fig. S7) and distributions of the read edit distance from the reference genomes (Fig. S8, 9). Mapping to reference genomes of the nine human-infecting species of *Herpesviridae* demonstrated the presence of authentic ancient viral DNA from four different HHV species in the two individuals, with average genomic coverage ranging from 0.02X to 0.48X and no cross-mapping between species (Fig. S7, Supplementary Tables 3-4). Three species, HHV5 (Cytomegalovirus), HHV6B (Roseolovirus) and HHV7 (Roseolovirus) were present in both individuals, whereas HHV1 (Simplexvirus) could only be conclusively detected in Yana1. Additionally, our data indicated the presence of HHV1 and possibly HHV4 in Yana2, however, due to the low numbers of mapped reads after quality filtering (HHV1 19 reads; HHV4 26 reads), the DNA damage patterns were too noisy to confirm their authenticity.

**Fig. 1.**
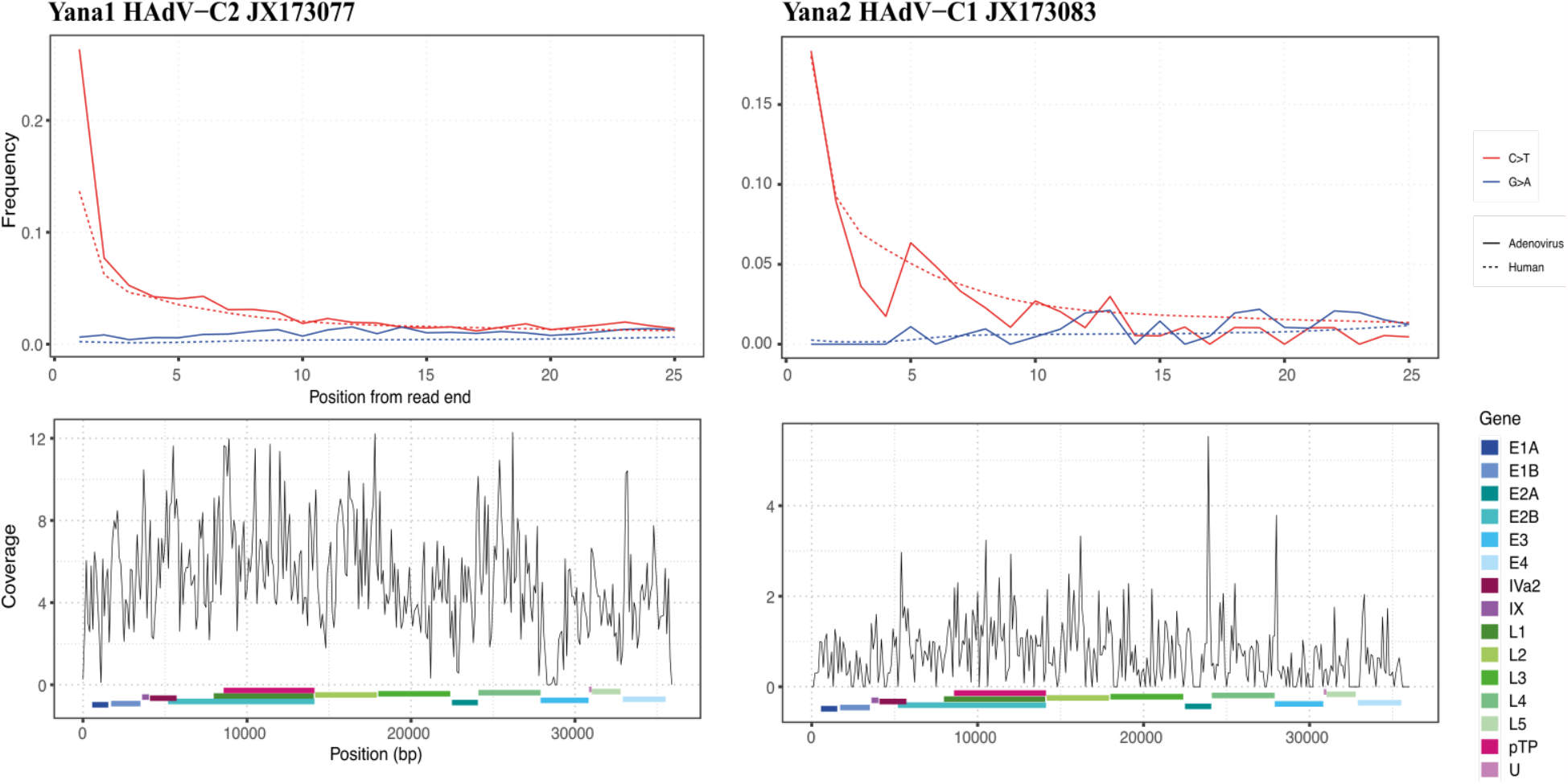
Ancient DNA authentication. Ancient DNA damage patterns and genomic coverage for HAdV-C in the two Yana individuals. (top) Rates of cytosine to thymine (C>T) and guanine to adenine (G>A) transitions, as a function of distance from the 5’ end for reads mapping to HAdV-C (full line) and human genomes (dotted line). (bottom) Average genomic coverage (200 bp windows) across the chosen HAdV-C reference genome for each individual. The start and end positions of HAdV-C gene regions are indicated by coloured boxes.

For the analysis of *Adenoviridae*, we used 44 HAdV-C reference sequences uploaded to NCBI (November 2018), as well as one reference genome for each of the remaining species isolated from simian, bat, fowl, porcine and bovine hosts. Results confirmed the presence of HAdV-C in both individuals, with aligned reads showing low edit distances (Fig. S9), as well as clear aDNA damage patterns at levels exceeding those observed for the ancient human DNA(*45*) (Fig. 1). We obtained an average genomic coverage of 5.2X (after removal of duplicates and reads with mapping quality below 30) against KF268310 (HAdV-C2), the reference genome with the best coverage for Yana1. However, because this reference genome lacks 161 bp at the beginning of the genome, which is present in many other HAdV-C genomes and Yana1, the reference sequence JX173077 with an average genomic coverage of 5.15X was selected for further analyses. For Yana2, average genomic coverage of 0.74X was obtained against reference genome JX173083 (HAdV-C1). Genomic coverage was generally highest for HAdV-C2 reference genomes in Yana1, and HAdV-C1 references in Yana2, indicating infections with different HAdV genotypes in the two individuals (Fig. S10). Conversely, only a small number of reads could be aligned to any of the other HAdV or non-human adenovirus reference genomes (Fig. S10). Coverage was even across the genome for both individuals, with 90.1% and 44.1% of the genome covered for Yana1 and Yana2, respectively (Fig. 1). A noticeable drop in coverage was observed for Yana1 in a hypervariable region of the E3 transcription unit, which encodes for proteins mediating immune evasion of the virus, which together with the fiber and hexon genes, drives genotype diversification(*35, 38, 39*).

### Ancient genome reconstruction, variants and recombination

To investigate how the ancient adenovirus sequences relate to modern virus isolates, we constructed consensus virus genomes from the Yana individuals. Limited coverage combined with very short DNA fragment lengths renders a direct virus genome assembly challenging. Thus, we employed an iterative mapping-based approach to maximize accuracy and completeness for the obtained consensus. First, individual genomes were constructed with mapping and variant calling to each reference genome used in the authentication. Then a final consensus was built from a multiple sequence alignment of these consensuses for each individuas and finally a consensus genome reconstructed for both.

We performed multiple sequence alignment of the two ancient HAdV-C genomes together with a set of 85 modern HAdV-C reference sequences (NCBI, April 2019), from which a total of 3,634 SNPs were called across 23,013 bp aligned sequence after excluding positions with gaps or missing data in the modern isolates. Clustering of virus sequences based on SNP genotypes showed a clear separation of the four major HAdV-C genotypes (Fig. 2, Fig. S11). Genotype clustering is largely driven by high genetic diversity and differentiation in the major capsid proteins hexon (transcript unit L3, aligned positions 18,118 - 22,543 bp) and fiber (L5, 31,605 - 33,367 bp), as well as the early gene region E3 (28,086 - 31,392 bp), as previously reported(*34, 35, 38, 39*) (Fig. 2, Fig. S12).

**Fig. 2.**
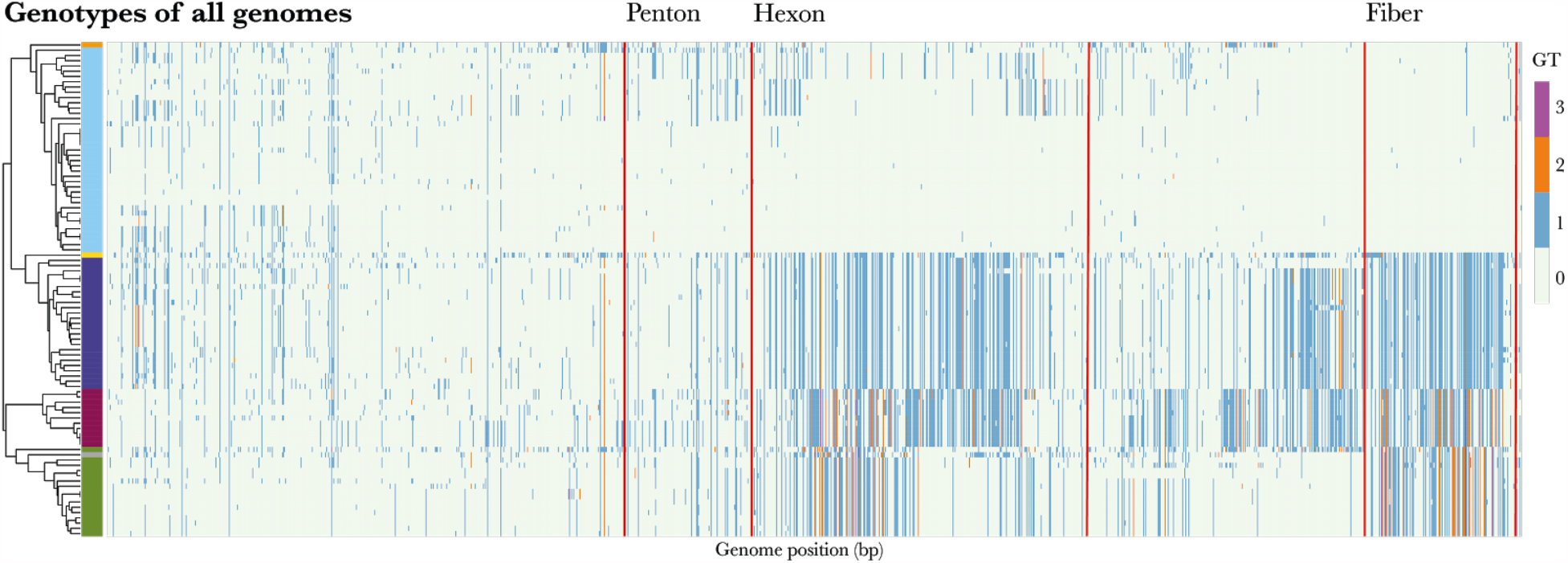
Genomic variants in the HAdV-C multiple sequence alignment. Genotypes of SNPs compared to a pseudo reference HAdV-C2 (JX173077) are indicated using different colours. The dendrogram shows the result of hierarchical clustering of modern and ancient genomes based on observed genotypes (GT), with major types indicated by coloured bars. Only variants without missing data in all samples were included. Genomic regions corresponding to variants in major capsid genes are indicated with read lines.

The two ancient genomes clustered with two different genotypes, HAdV-C2 for Yana1 and HAdV-C1 for Yana2, consistent with the higher genomic coverage obtained when mapping against these respective reference genomes (Fig. S10). Genome-wide genetic similarity as measured by Average Nucleotide Identity (ANI) also supported these results. For Yana1, ANI was highest with HAdV-C2 genotypes (98.1-98.4%), whereas for Yana2 it was highest with HAdV-C1 genotypes (97.1 – 97.9%), both exceeding their similarity to each other (96.3%) (Fig. S13-15). Investigating ANI locally along the alignment, we found the highest genetic similarity between Yana1 and Yana2 and their respective modern genotypes C2 and C1 in the high diversity regions of the major capsid genes hexon (L3) and fiber (L5) of (Fig. 3, Fig. S16, 17). The presence of two distinct virus genotypes in the individuals from Yana at ∼31,600 ybp therefore suggests a substantially older age for their diversification.

**Fig. 3.**
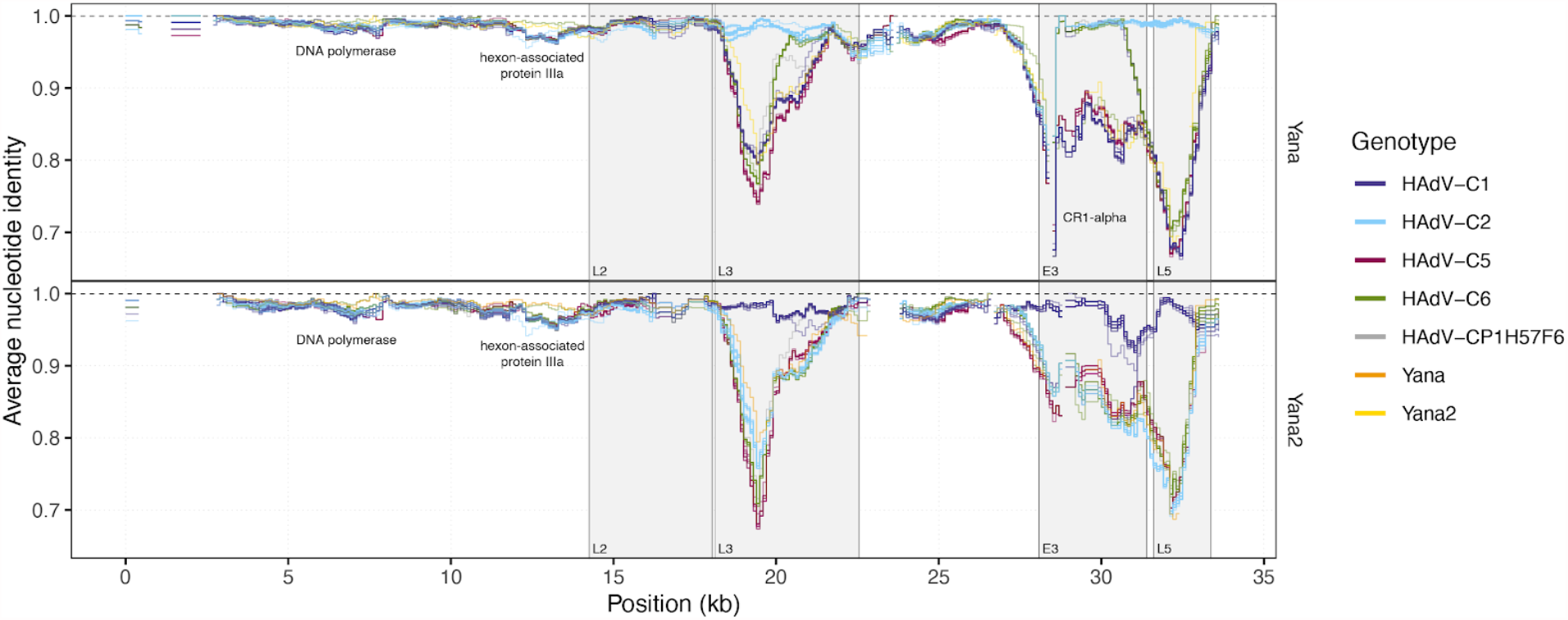
Pairwise genetic similarity of ancient Adenoviruses. Coloured lines indicate average nucleotide identity (ANI) in 1kb sliding windows (100 bp step size) between the ancient genomes and modern isolates. Positions of major capsid genes L2, L3, L5, as well as early gene region E3 are indicated by grey boxes.

Another region yielding an interesting observation is the CR1-alpha gene in the E3 transcript unit (28,862-29,060 bp), where Yana1 shows a drop in coverage flanked by regions of ANI < 80% with any other virus genome, suggesting the presence of a gene variant that is too differentiated to be recovered by mapping to present-day variation (Fig. S16). Finally, in transcript units E2B (DNA polymerase, 5,244-8,823 bp) and L1 (hexon-associated protein IIIa, 12,404-14,164 bp) both Yana genomes showed higher differentiation to all modern isolates than to each other, with the exception of strain UFL_Adv6/2005/6 (accession KF268129.1), a genotype 6 virus from Colorado, the USA in 2005 (Fig. 3, Fig. S16, 17). Hence, parts of this North American modern adenovirus isolate seem to be more closely related to the viral genomes in circulation ∼30,000 ybp than to any sampled present-day genomes currently available. We also observed a higher rate of unique variants for the two Yana individuals than in most of the modern isolates. Unique variants were clustered in regions of high overall diversity, indicating the presence of diverse haplotypes specific to those individuals (Fig. S18, 19).

### Recombination and chromosome painting

Recombination events between viral genomes can be a major driver of virus evolution and are frequently observed in human adenoviruses(*34*–*39, 48*). Consistent with this notion, we found that the variation in ANI between the ancient and modern virus genomes along the genome was not compatible with strictly clonal evolution. In many genomic regions outside the genotype-differentiating L3 (hexon) and L5 (fiber) transcript regions, the ancient genomes had a high ANI with isolates belonging to other genotypes. This may indicate intertype recombination in high diversity regions such as e.g. HAdV-C2 derived E3 genes in HAdV-C1 isolates EGY/E13/2001/1 and 43C1, and particularly in the regions upstream of the L3 transcript unit (the first ∼18kb of the virus genome alignment) (Fig. S20).

To quantify the rate of recombination, we examined pairwise linkage disequilibrium (LD) in the modern sequences using the classical estimators D’ and *r*^*2*^. We observed high levels of LD both within and between the genotype-determining regions L3, E3 and L5, but low linkage elsewhere along the genome consistent with a recombining genome (Fig. 4). The strong linkage between the major capsid genes L3 (hexon gene) and L5 (fiber gene) was also reflected in the absence of intra-genotype recombination observed for the two ancient genomes. Average LD, measured by *r*^*2*^ decayed rapidly with the distance between SNPs, showing pairwise *r*^*2*^ values below 0.1 at distances above ∼4.5kb (Fig. 4). Fitting a decay curve to the observed LD decay (equation in appendix 2 of Hill and Weir(*49*)), we estimated the population-scaled recombination rate for the modern isolates at ρ=1.9×10^−3^ / bp / generation. When excluding the linked regions L3, E3 and L5, the rate of recombination was estimated more than an order of magnitude higher at ρ=4.2×10^−2^ / bp / generation (Fig. S21).

**Fig. 4.**
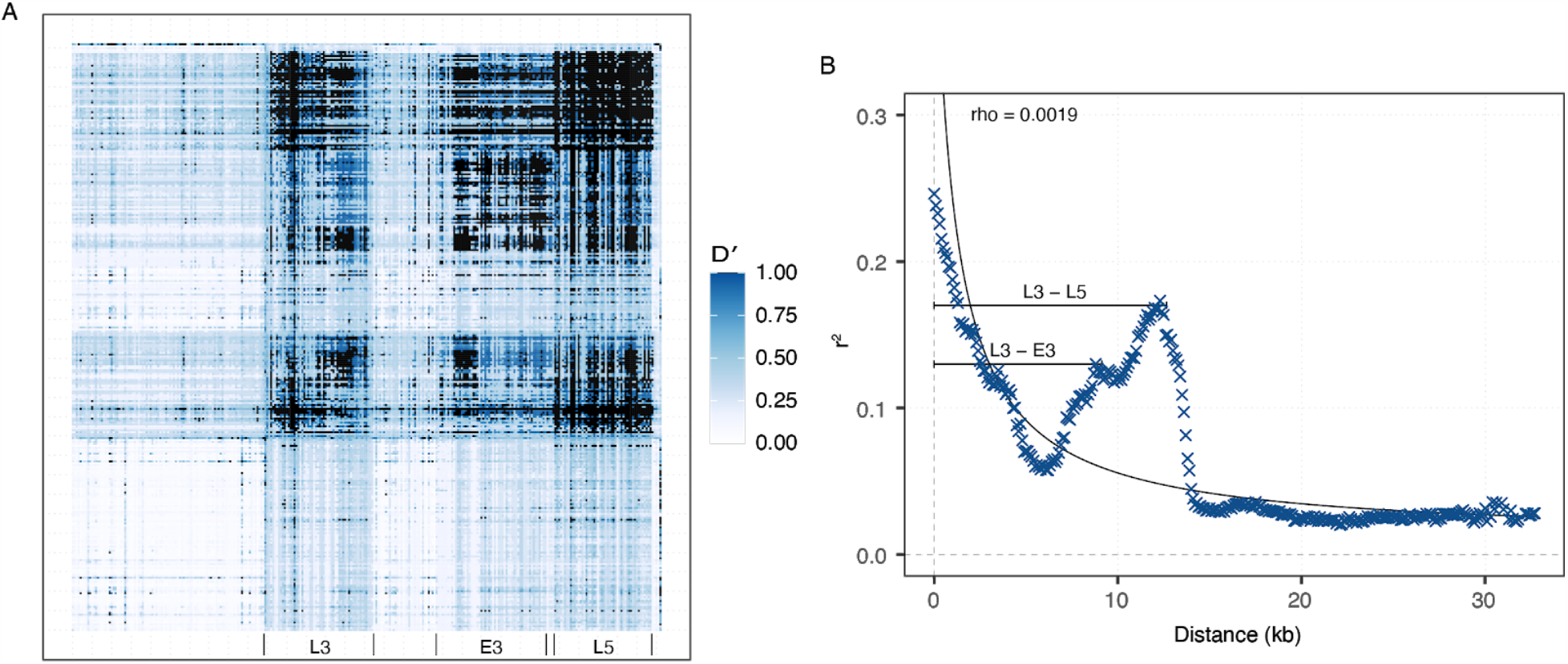
Linkage disequilibrium in HAdV-C. (A) Heatmap of pairwise LD using median D’ aggregated in bins of 10 consecutive SNPs. Bins corresponding to SNPs in the L3, E3 and L5 transcript units are indicated. (B) LD decay curve showing average pairwise r2 as a function of the distance between SNPs (bins of 100 bp). The black curve indicates the theoretically expected decay from the estimated population recombination rate. The two peaks in r2 around 10kb correspond to the distance between gene pairs L3-L5 and L3-E3 (indicated with the black line).

We used Chromopainter(*50*) to analyse the relationship between the modern and ancient HAdV genomes while accounting for recombination. With this approach, an “ancestry painting” for the genome of each “recipient” strain is obtained by modelling them as a mosaic of chunks of DNA copied from a panel of “donor” genomes. Painting each recipient genome using all other genomes as possible donors, we found that the majority of recipient genomes copied predominantly from donors of the same genotype (Fig. 5; Fig. S22-26). However, many recipients also copied substantial parts of their genome from donors of other genotypes, as expected in the presence of frequent recombination between types. The two ancient genomes could be modelled as mixtures with varying proportions of chunks donated from all other genotypes, without clear dominance of any single type. Examples of copying from the two ancient donor genomes were only rarely observed in the modern recipients, consistent with their distant evolutionary relationship. A noticeable exception was modern strain UFL_Adv6/2005/6 (HAdV-C6, KF268129), which could be modelled as deriving >50% of its donated chunks from Yana1 (HAdV-C2). Investigating copying probabilities locally along the genome revealed that copying from Yana1 was predominantly found in the first half of the virus genome upstream of L3, which also showed high ANI with Yana1 (Fig. S16, 23). These results suggest that UFL_Adv6/2005/6 represents an ancient lineage, which likely recombined with HAdV-C6 related L3 and L5 genes(*35*).

**Fig. 5.**
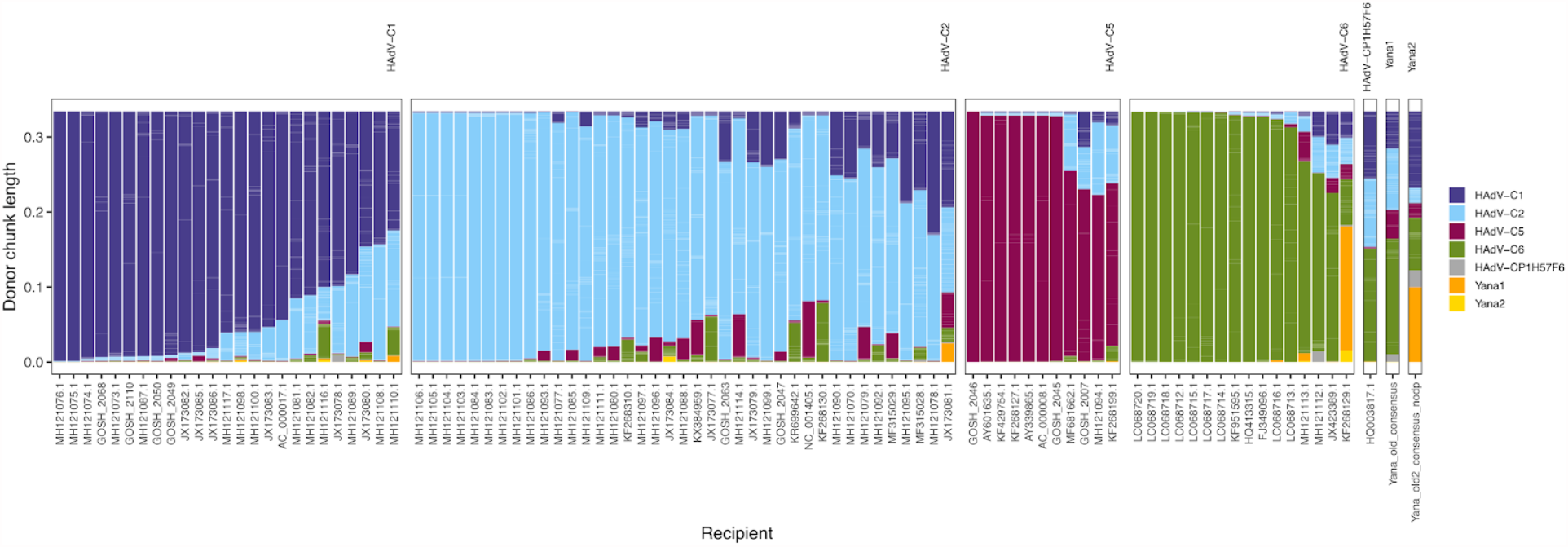
Chromosome painting of viral genomes. Bar plot showing total length of chunks donated to recipient genomes from all other donor genomes, coloured by the genotype of the donor.

### Phylogenetic reconstruction and dating

We reconstructed a maximum likelihood phylogenetic tree from the whole-genome alignments using RAxML(*51*). The resulting tree separated the viral genomes into four clades, corresponding to the predominant genotypes of HAdV-C 1, 2, 5 and 6 (bootstrap support > 95%; Fig. S27). The higher coverage Yana1 virus genome formed a clade with the modern strain EGY/E52/2001 from Egypt (accession JX173081.1), diverging basally from all other modern HAdV-C2 types. The strain EGY/E52/2001 has previously been shown to differ notably from other HAdV-C2 genomes(*35*), and its phylogenetic relationship with Yana1 appears to be driven by harbouring closely related variants of the L3 gene (Fig. S16, 28, 29). The lower coverage Yana2 virus genome fell within the diversity of other HAdV-C1 genomes. With the two ancient Yana genomes falling distinctively in different lineages, our genome-wide phylogeny also supports the diversification of the four major genotypes predating 31,600 years.

Phylogenetic trees built separately for all annotated genes and their coding DNA sequences (CDS) revealed considerable differences in tree topology between genes and even between coding sequences within a gene, consistent with recombination between virus types (Fig. S28, 29). Of the early genes, only E3 (host modulating and death proteins) and E2A (ssDNA binding protein) clustered according to major genotypes. For E3, three major clades corresponding to genotypes 2 and 6, genotype 1, and genotype 5 were observed, and the genes from the ancient strains fell in positions basal to clades of modern sequences (Fig. S28, 29). Only a weak phylogenetic signal was observed for the remainder of the early genes, E1A (transcription activation and induces the host cell to enter the S phase), E1B (blocking apoptosis), E2B (DNA polymerase) and E4 (mediating transcription, RNA splicing and translational regulation, mRNA nuclear transport and modulating DNA replication and apoptosis), as well as intermediate genes IX (hexon associated protein) and Iva2 (maturation protein, packaging protein). Gene sequences from the two ancient strains were mostly found clustering together within the same clade, sometimes exclusively so except strain UFL_Adv6/2005/6 discussed above. The late genes code for the structural components of the virus, as well as the encapsidation and maturation of the virions in the nucleus. Among them, both L1 and L2 (penton), which defines part of the virus classification, did not clearly separate by genotype but clustered the ancient sequences in a clade together with UFL_Adv6/2005/6. The genes for the major capsid proteins L3 (hexon) and L5 (fiber) on the other hand led to genotypes clustering into deeply diverged clades. L3 and L5 sequences from both ancient genomes fell either basal to the entire clade or to a clade containing the majority of sequences for a genotype, consistent with their ancient age.

To investigate the evolutionary timing of HAdV-C diversification, we performed molecular dating of the HAdV-C phylogeny, incorporating the two ancient genomes as tip calibration points. We chose not to include an outgroup as all possible outgroups are very distant from HAdV-C and the genome-wide tree topology is well supported. The tree was instead rooted with TempEst(*52*) so as to maximise the temporal signal. Measurable temporal evolution over the phylogeny was inspected using Phylo STemS(*53*). Initially, testing an unfiltered alignment, no statistically significant temporal regression was observed at the base of any of the four major clades likely due to extensive recombination in the history of HAdV-C (Fig. S30). To exclude sites putatively deriving from recombination, we identified all homoplasies in the alignment which were subsequently excluded. The homoplasy-pruned alignment still lacked measurable temporal evolution at any basal clade, likely due to some samples placed on tips with long terminal branches (Fig. S31). A possible explanation for the presence of unusually long terminal branches is that viral isolates, such as adenoviruses, are generally serially passaged prior to whole-genome sequencing, which typically leads to the accumulation of artefactual mutations. To circumvent this problem, we identified all singleton variants in the alignment which were further pruned. The resulting maximum likelihood phylogeny, following these strict filtering criteria to account for recombination and artefactual mutations, resulted in a statistically significant temporal regression (adjusted *R*^*2*^=0.44, p-value<0.0002) at the clade of genotype 1 including Yana2 (Fig. S32, 33).

We initially explored the time to the Most Recent Common Ancestor (tMRCA) using Bayesian tip-calibration implemented in BactDating(*54*). We obtained a weak but significant regression (*R*^*2*^=0.04, p-value=0.034) following 10,000 permutations of the sampling date (Fig. S35) allowing estimation of the tMRCA, using the bactdate() function, to 726,828 BP (2,585,687-152,225; 95% credibility interval) (Supplementary Table 5). While we also obtained a significant temporal regression when omitting the ancient samples (*R*^*2*^=0.05, p-value=0.031) (Fig. SS36), the estimated age of the root was strikingly more recent than the age of our Yana samples and with a marginally worse model likelihood (tMRCA 7297 BP [-11,801 - 2,388 CI]) (Supplementary Table 5). To refine these estimates, we next conducted formal Bayesian estimation of evolutionary rates and divergence times using BEAST2(*55, 56*). As no substitution model had overriding support (maximum 42.42% posterior support, Fig. S39), a model averaging approach was applied. Following convergence and evaluation of a range of demographic and clock models, the best-supported model was a coalescent Bayesian skyline model with a strict prior on the clock rate (Supplementary Table 6-7, Fig. S40-43).

Results using both approaches and across all demographic models tested consistently supported a very deep divergence of the HAdV-C phylogeny, with an estimated mean substitution rate of 2.52×10^−10^ substitutions / site / year. The time-calibrated phylogeny clustered the predominant genotypes (1, 2, 5, 6) into four major clades, with a common ancestor estimated to 701,850 ya (95% HPD 487-963 ya) (Fig. 6). The divergence dates between all HAdV-C major genotypes were also old, with the most recent divergence estimated between genotype 2 and 6 to around 351 to 698 kya. Interestingly, the evolution of within-genotype diversification was estimated to have occurred only within the last ∼70,000 years for all four major genotypes. Interestingly, the main Neanderthal admixture event into the early out-of-Africa population has been dated to the same time period (*57, 58*).

**Fig. 6.**
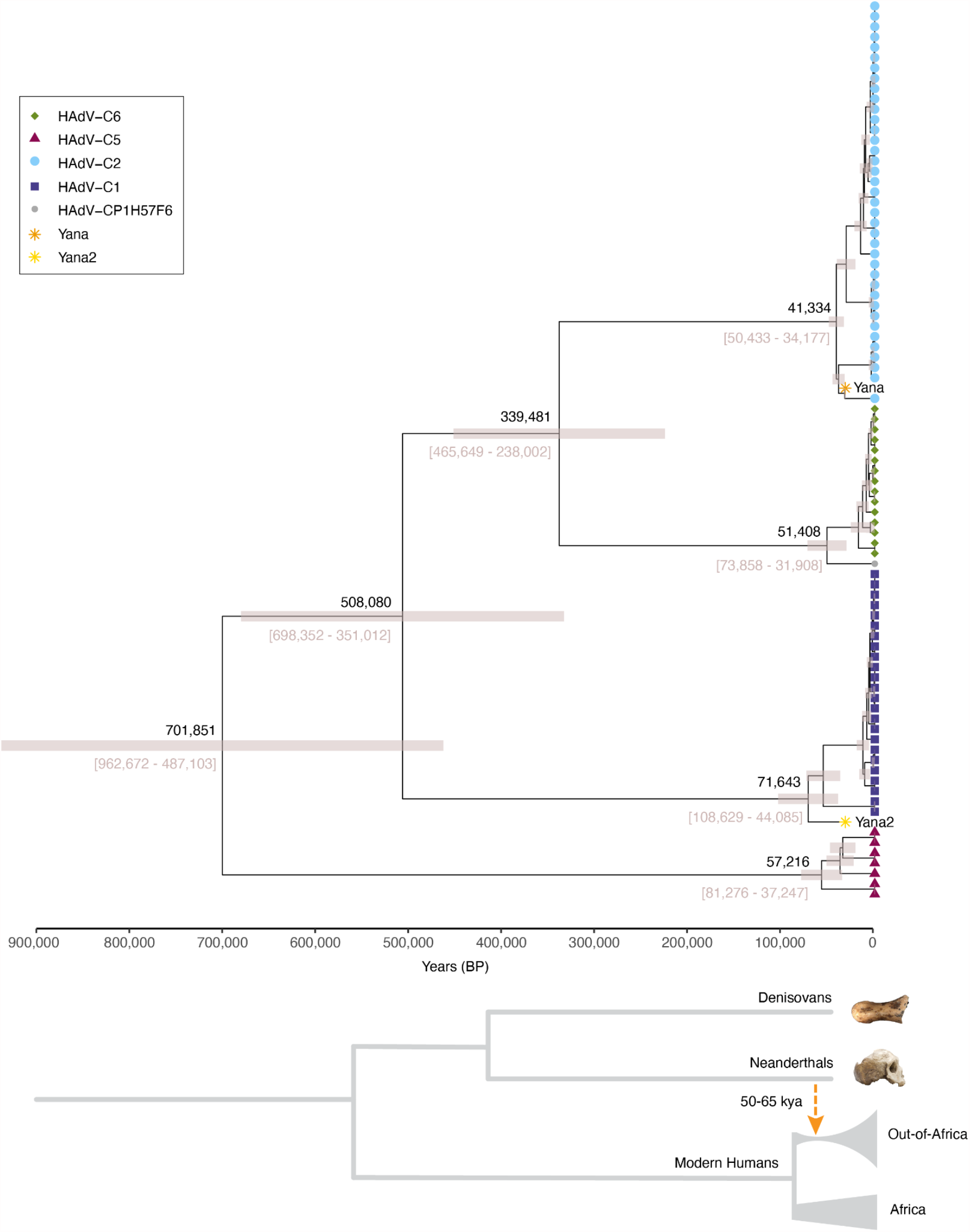
Dating the evolutionary history of HAdV-C. Phylogenetic tree of HAdV-C sequences with divergence dates and 95% HPD interval for major type divergences, estimated using BEAST2 and a coalescent Bayesian skyline model with a strict molecular clock. A comparative tree illustrating population divergence and admixture times for representative archaic and modern human groups is shown below the inferred HAdV-C phylogeny.

## Discussion

Recent progress in the recovery and analysis of ancient pathogen DNA has dramatically improved our understanding of the history of human infectious diseases(*13, 59*). In this study, we found the first molecular evidence that early modern humans in the Pleistocene hosted common childhood viral infections, such as adenovirus and herpesvirus. Our findings show that recovery of viral DNA and reconstruction of near-complete genomes is feasible from DNA extracted from human teeth dating back 31,600 years, pushing back the earliest evidence for human viral infections by ∼25,000 years. We retrieved the DNA of multiple viral species responsible for common childhood viral infections from the ancient samples, including at least four species of human herpesvirus (HHV1, HHV5, HHV6B and HHV7) as well as two genotypes of human adenovirus C (HAdV-C1, HAdV-C2). All of these viruses are capable of establishing life-long latent or persistent infections in humans(*60*–*62*), making it unlikely that the viral DNA recovered from the two milk teeth originated from active infections at the time of tooth shedding. Human cell types for latent HHV1 infections are typically neurons while other herpesviruses (HHV5, HHV6B, HHV7) and adenovirus HAdV-C analysed in this study are most likely found latent in immune cells(*63, 64*). Chromosomal integration has been shown to occur for HHV6B(*65*), but the low coverage of the virus genomes (Yana1 0.14X, Yana2 0.06X) compared to the respective human genomes (Yana1 25X, Yana2 7X(*45*)) indicate that the amplified viral DNA was most likely not from human genome integrated sites and is not an inherited chromosomally integrated copy. Taken together, our results demonstrate the power of deep sequencing of well-preserved ancient teeth to retrieve ancient viral DNA from likely low viral load, latent or persistent infections and as such to a record of previous virus infections of prehistoric individuals.

We reconstructed near-complete genomes of two 31,630-year-old HAdV-C viruses, revealing surprising new insights into human adenovirus diversity and evolution. All modern HAdV-C genomes sequenced to date are characterized by a conserved virus backbone, interspersed with regions of high diversity including the major capsid genes L3 (hexon) and L5 (fiber) as well as the host and immune-modulating early gene region E3(*34, 35, 39*). Both ancient genomes were found to share >97% average nucleotide identity with modern genomes along most of the conserved virus backbone, as well as with their respective genotype prototypes in the L3, E3 and L5 regions. Our findings reveal an unexpected genetic similarity of the two ancient adenovirus genomes with their most closely related modern adenoviral lineages, despite being separated by over 31,000 years of evolution. Notably, the modern adenovirus strain UFL_Adv6/2005/6 (KF268129.1) carries a gene region variant more closely related to the ancient Yana1 HAdV-C genome sample than to any other known modern adenovirus lineage. Similarly, the modern adenovirus EGY/E53/2001/2 (JX173081.1) contained a variant of the L3 region diverging basal to other modern HAdV-C L3 gene lineages, but most closely related to the L3 gene variant found in Yana1. Altogether the close genetic relatedness of the ancient HAdV-C adenovirus genome variants retrieved to the modern adenoviral genetic diversity suggests that the two newly discovered traces of ancient lineages have largely evaded lineage extinction and instead have persisted with humans through time since the Pleistocene. Interestingly, our findings are in contrast to recent studies of pathogen diversity for other viruses retrieved from ancient samples representing extinct clades not observed in present-day or recent viral populations (*24, 25, 27, 28*). Nevertheless, we also show that both nearly complete ancient adenoviral genomes differentiate from all 102 publicly available modern HAdV-C genome lineages reported thus far (*35*). The anticipated large ancient population size of HAdV-C viruses infecting humans in combination with persistent and/or latent infections, prolonged viral shedding, high rates of transmission and low virulence are the likely essential viral features, which have sustained the long common history and diversity of HAdV-C viruses with humans.

Apart from genetic drift caused by mutations, genomic recombination is the main force governing adenovirus evolution. Compared to other adenovirus species, HAdV-C has been suggested to be more restricted in its recombinational activity. For example, intra-type recombination between the major capsid genes L3 (hexon) and L5 (fiber) is frequently observed in HAdV-D, but is rarely reported in HAdV-C strains(*4, 34, 38, 66*). Our analysis of patterns of linkage disequilibrium confirmed strong linkage between the L3 and L5 as well the E3 regions, which all exhibit deeply diverged gene trees of co-segregating genotypes. However, despite their co-segregation with the major HAdV-C genotypes, gene trees for L3 and L5 show distinct topologies, indicating an ancient recombination event in the history of currently circulating types. Our finding that both ancient genomes clustered with their respective closest type demonstrate that currently circulating linkage types were already established over 31,000 ybp, thereby suggesting a strong functional association between hexon and fiber protein types. The remaining viral genome shows low levels of LD, consistent with an essentially freely recombining sequence. Highly diverse gene regions on a genomic backbone of freely recombining regions have also been observed for other large dsDNA viruses causing low virulence and latent infections such as HHV5 (cytomegalovirus), therefore suggesting similar genomic evolutionary dynamics(*67*).

Large dsDNA viruses are generally characterized by lower mutation rates than other virus classes such as RNA viruses(*42, 43*). As a consequence, many species of dsDNA viruses, including herpesviruses and adenoviruses, are commonly thought to have evolved through a mechanism of long-term association and co-divergence with their host species. Previous studies have estimated low rates of molecular evolution, on the order of 10^−8^ to 10^−9^ substitutions per site per year (s/s/y), consistent with this notion(*43, 68*–*70*). However, these rates have often been derived using phylogenies calibrated under the assumption of host co-divergence, due to the absence of direct fossil calibration points. When heterochronous samples were included in the analyses, surprisingly high rates of evolution (10^−4^ to 10^−6^ s/s/y) were found, conflicting with the long-term co-divergence scenario(*12*). A likely explanation for this discrepancy lies in the shallow time depth of the available virus genomes used, which date back ∼70 years or less. The ancient virus genomes presented in this study circumvent this issue by providing, for the first time, direct molecular evidence of the long-term association of humans with herpesvirus and adenovirus, dating back to the Pleistocene.

Our molecular divergence analysis using BEAST2(*55, 56*) suggests a very low rate of evolution in HAdV-C, with an estimated substitution rate of 2.52 × 10^−10^ s/s/y, consistent with a species co-divergence scenario. We demonstrate the value of including ancient genomes in dating divergence times between viruses, especially for species with a slow mutation rate. This is also highlighted by the estimates of tMRCA which would have pointed to a recent origin (7297 BP [-11,801 - 2,388]) had we only included modern isolates. This is in stark contrast with the much older estimates of 726,828 BP [2,585,687-152,225] obtained when including the ancient genomes. Furthermore, we show the importance of careful treatment of alignments to exclude recombinant or putatively artefactual mutations prior to the conduction of temporal analyses. It is important to note that we do not obtain precise divergence dates (leading to the large confidence intervals observed), however, the three demographic coalescent models tested (constant, exponential and Bayesian skyline) all converge on comparable tMRCA point estimates (Fig. S44) with posterior distributions significantly different from those recovered when sampling from the prior.

Our analyses point to a strikingly ancient origin and deep divergence of the currently circulating HAdV-C genotypes. Our estimated time to the common ancestor of all HAdV-C lineages dates back ∼700,000 years, and all estimated split times between the major genotypes sampled today were older than 300,000 years (Fig. 6). Recent genomic analyses indicate a separation between modern and archaic (i.e. Neanderthal and Denisovan) human ancestors between 550,000 to 700,000 ybp, followed by the establishment of present-day human genetic diversity within the last ∼250,000 years(*71, 72*). Our results thus suggest that the current HAdV-C genetic diversity pre-dates modern human origins, and likely originated within our hominin ancestors. Whether this diversity is the result of divergent virus lineages migrating out-of-Africa with their human hosts, or cross-species transmission from archaic humans during admixture outside of Africa remains unknown. Intriguingly, our estimates of within-genotype divergence times fall only within the last ∼70,000 years, which would be consistent with the latter scenario as the main Neanderthal admixture event into the early out-of-Africa population has been dated to the same time period(*57, 58*). A similar transmission from archaic to modern humans has also been suggested for human papillomavirus 16, although based solely on analyses of present-day virus genomes (*69, 73*). The lack of present-day geographic structure further supports the deep divergence. Future studies on Neanderthal and Denisovan viral infections may resolve these questions, and our study demonstrates the feasibility of Pleistocene viral phylogenomics.

## Acknowledgments

We thank the staff at the GeoGenetics Sequencing Core (Copenhagen; seqcenter.ku.dk), and Eske Willerslev for funding support and scientific discussions.

## Author contributions

Conceptualization: SHN, MS, LVD, FB

Methodology: SHN, MS, LVD, FB, LV

Investigation: SHN, MS, LVD

Visualization: SHN, MS

Funding acquisition: MS

Project administration: SHN, MS

Supervision: MS

Writing – original draft: SHN, MS

Writing – review & editing: SHN, MS, LVD, FB, CJH, AGP, MEA, LV, AM, EP, VC, PN, VP, VNP

## Data and materials availability

Sequencing data and consensus genomes are available in the European Nucleotide Archive under project number XXXX

