## supplemental material for "31,600-year-old human virus genomes support a Pleistocene origin for common childhood infections"

### Materials and Methods

Screening and ancient DNA authentication

Screening for ancient virus DNA was carried out on previously generated whole-genome shotgun sequencing data *(45)*. For individual Yana2, we performed additional targeted enrichment of viral sequences on a single dual-indexed sequencing library, with hybridization and sequencing on HiSeq4000 platform performed as described previously *(25)*.We used the metagenomic classifier Kraken (version 1.0)(*46*), using a custom-built database of all microbial and viral genomes in the RefSeq database (February 2017) for in-silico pathogen screening. Following identification of putative ancient herpesvirus and adenovirus hits in both individuals, all reads were aligned to diverse sets of reference genomes with bowtie2 (version 2.3.2)(*47*), using both end-to-end as well as local alignment options. We used a modification of the 'very sensitive' preset allowing for a mismatch in the alignment seed ('-N 1' option) to maximize the number of putative virus reads aligned. Ancient DNA authentication was carried out by assessing patterns of typical ancient DNA miscoding lesions using mapDamage (version 2.0.8-dirty)(*74*), evenness of coverage along the virus genomes, as well as distributions of edit distances against different reference genomes.

Adenovirus genome reconstruction

Reconstruction of ancient HAdV consensus genomes was carried out using read alignments against all available HAdV-C reference genomes (n=44) in NCBI (November 2018). Following initial alignment against each reference genome, PCR duplicates were removed and reads were filtered for mapping quality (MQ) ≥ 30 using samtools(*75*). We further removed reads with read length >70 bp for Yana1 and > 60 bp for Yana2, based on the observed read length distributions from mapDamage (Supplementary Figure 4, 6). We called variants using bcftools (version 1.9-94-g9589876)(*75, 76*) against each reference genome and used them to build initial consensus sequences for both ancient samples. For the higher coverage Yana1 sample, we required that included positions were covered by a minimum of three reads, and with a minimum quality ('QUAL' field) ≥ 20. For the lower coverage Yana2 we included all sites to maximize coverage across the virus genome. The final consensus dataset was constructed following multiple alignment of the individual consensus genomes.

Diversity and recombination

A multiple sequence alignment of the two ancient genomes and a set of 85 modern HAdV-C isolates (NCBI April 2019) was obtained using MAFFT (version 7.427)(*77*), with 1000 iterations and local pairs. The alignment was manually checked for alignment errors in AliView (version 1.25)(*78*). Single-nucleotide variants for population genetic analysis were extracted from the alignment using snp-sites (version 2.4.1)(*79*), using JX173077.1 as an internal pseudo-reference, and subsequently used for genetic similarity analyses. Linkage disequilibrium was determined using tomahawk (version 0.7.0). Chromosome painting analyses were carried out using Chromopainter (version 2)(*50*), assuming a uniform recombination map and following EM estimation of mutation and switch rates. We performed painting analyses using different sets of donor and recipient genomes, including an all against all analysis, as well as different subsets with and without the ancient samples included in the donor panel. Results were visualized using R and the tidyverse suite of packages(*80*).

Phylogenetic analyses

Maximum likelihood phylogenetic trees were built with RAxML (version 8.2.11)(*51*) using a GTR+I+G model (i.e., gamma-distributed rate heterogeneity and an estimated proportion of invariant sites)(*81*), and 100 bootstrap replicates. For all gene and CDS trees, invariant sites were not included. Phylogenies were rooted using Simian adenovirus 43 (accession FJ025900.1) as an outgroup, which was aligned to the HAdV-C alignment using MAFFT with '—add' and '—keeplength' options.

Homoplasies were identified using the R package HomoplasyFinder(*82*), using both the alignment and a maximum parsimony tree obtained in MEGA with the 'complete deletion' specification (version 7.0.26). The detectable temporal signal was explored using Phylo STemS(*53*) using maximum likelihood trees obtained from the raw alignment, a homoplasy filtered alignment and a further alignment which also excluded singleton positions. Root-to-tip analyses and divergence dating were carried out using the BactDating(*54*) package in R, as well as BEAST version 2.6.1(*54, 55*). Model averaging was conducted in BModelTest (posterior support model 121323, 42,42%; model 121343, 28.25%; Supplementary Figure 39). All other priors were set to default values.

Three possible coalescent models (constant, exponential, Bayesian skyline) were tested under both strict and relaxed priors on the molecular clock rate. MCMC chains were run until convergence which was inspected in Tracer and required effective sample size (ESS) value >200. With a relaxed clock, 1.9 billion iterations were required, whereas assuming a strict clock model, all converged with 880 million iterations or even less, though the majority were run for the same chain length for model comparison (Supplementary Table 6). Each strict clock model was run three times with different seeds and concluded in the same result. The best raw and marginal likelihood was calculated with PathSampler requiring 35 steps and 700.000 iterations, selecting the Bayesian Skyline model as having the highest support. This model was also run without the data 'sampling from the prior' to ensure the posterior differed (with data 3.797E6 and without data -325.887). TreeAnnotator was used to summarise the tree samples as the maximum credibility tree, discarding 10% as burn-in. Trees were generated for both the mean and median heights and were highly consistent. Mean heights were used for tree visualization in R with ggtree version 1.16.6(*83*).

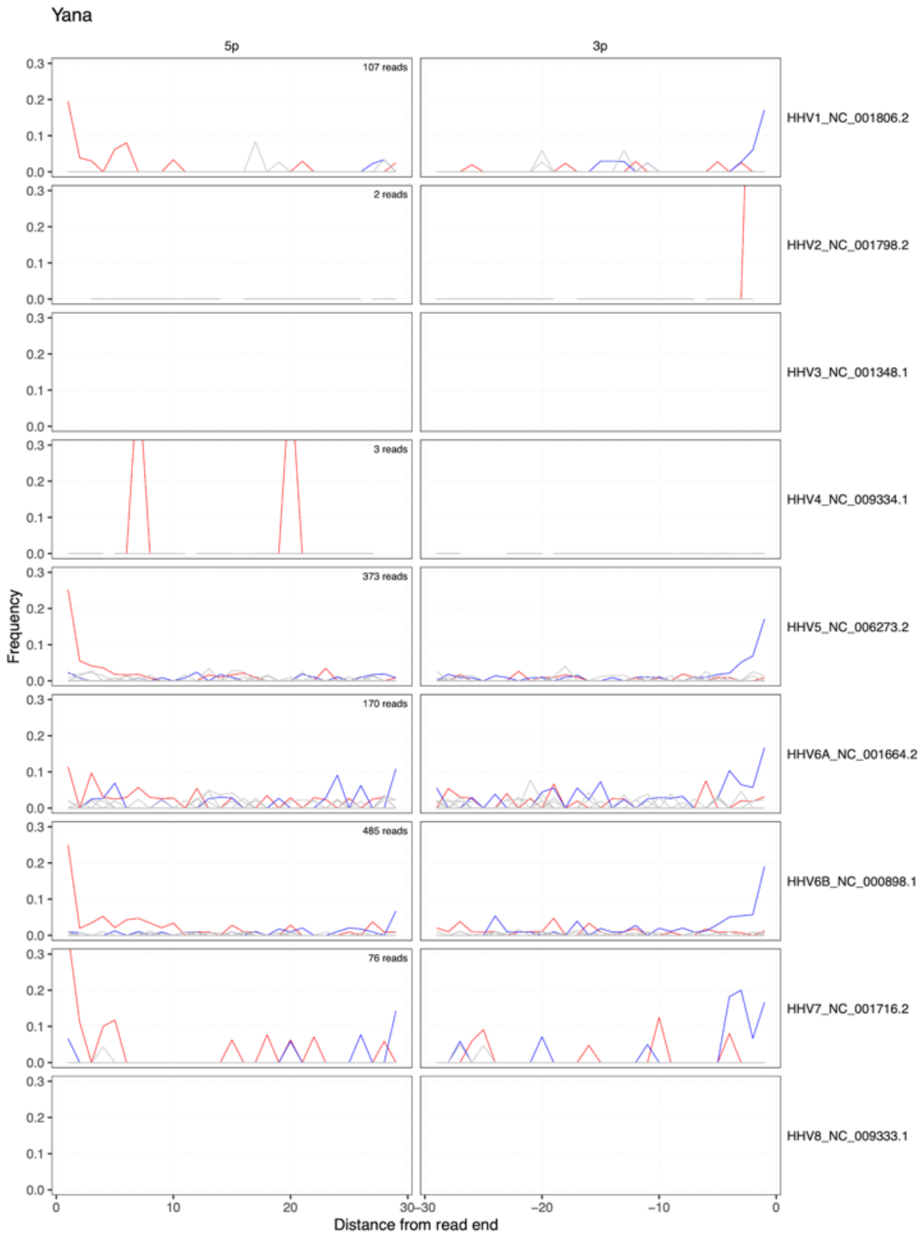

**Fig. S1. Herpesvirus ancient DNA damage patterns for individual Yana1.** Rates of nucleotide substitutions for mapping against nine herpesvirus reference genomes, as a function of distance from the read ends. Coloured lines indicate cytosine to thymine (red) and guanine to adenine (blue) transitions.

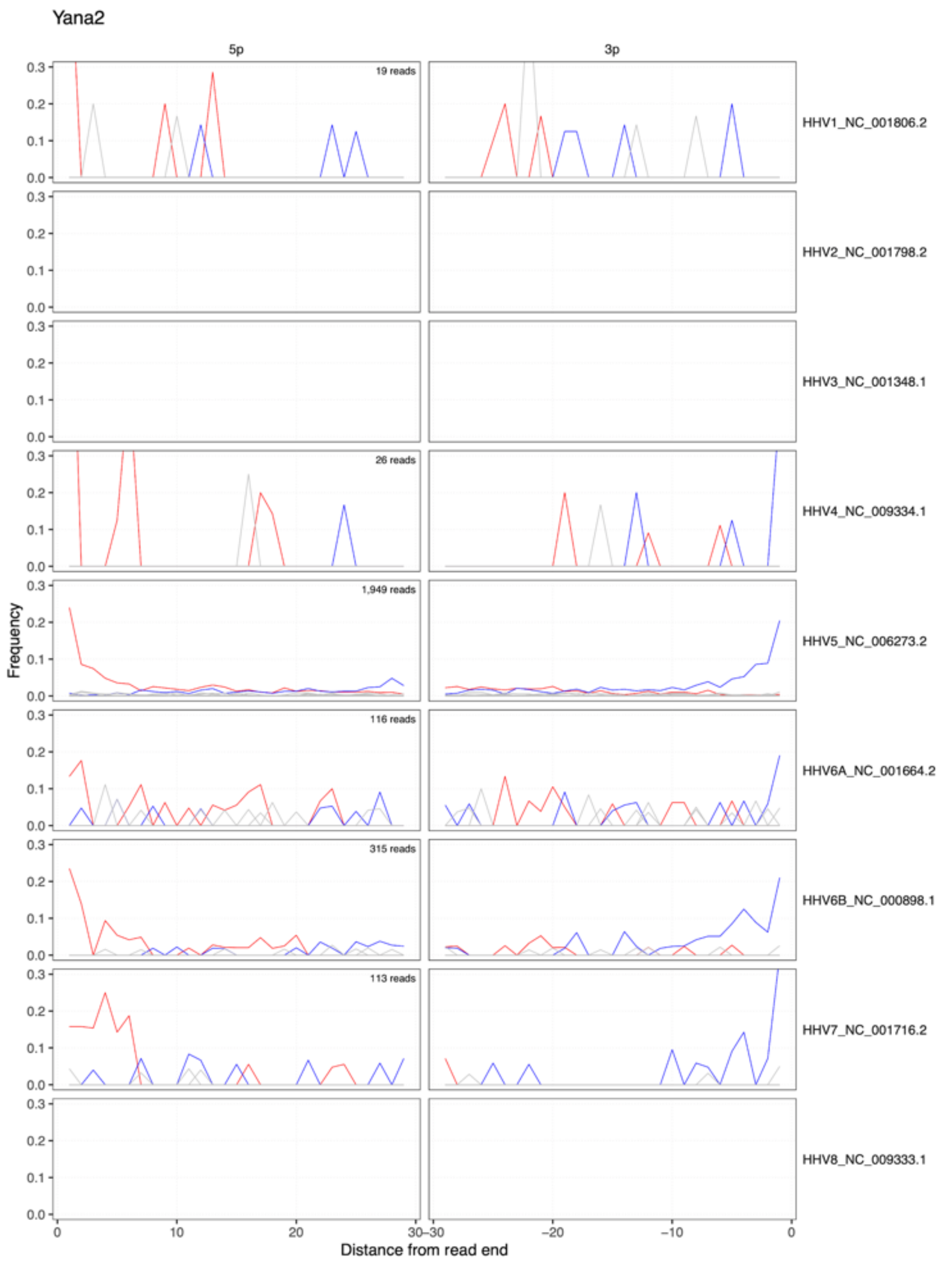

**Fig. S2. Herpesvirus ancient DNA damage patterns for individual Yana2.** Rates of nucleotide substitutions for mapping against nine herpesvirus reference genomes, as a function of distance from the read ends. Coloured lines indicate cytosine to thymine (red) and guanine to adenine (blue) transitions.

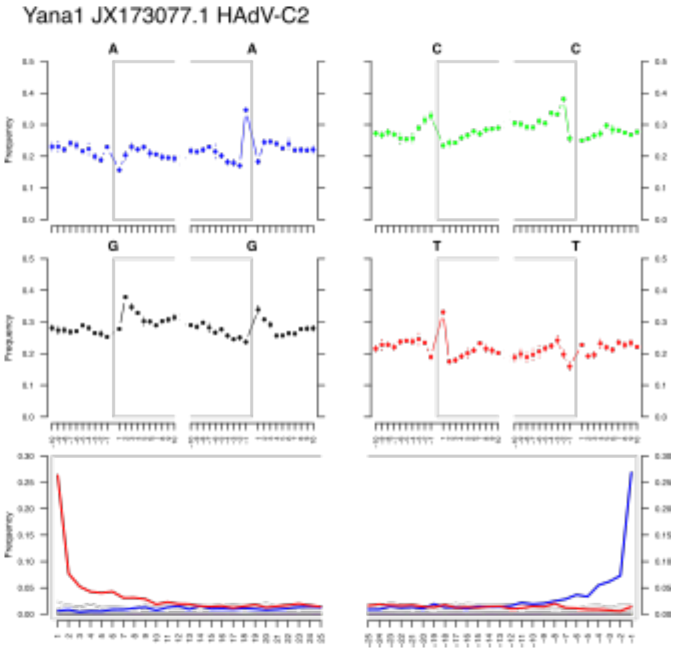

**Fig. S3. Adenovirus ancient DNA authentication for individual Yana1**. Top, nucleotide compositions of genomic regions for read ends and flanking positions. Bottom, cytosine to thymine (red) and guanine to adenine (blue) transitions.

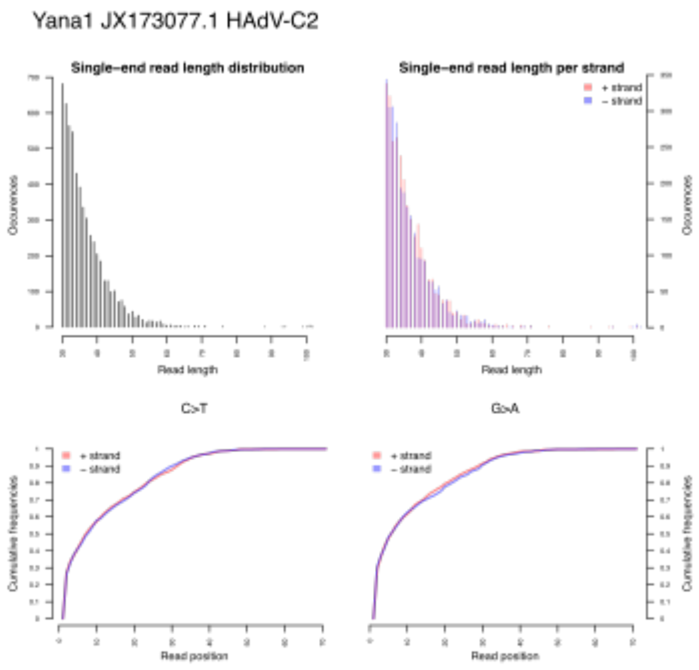

**Fig. S4. Yana1 read length distribution** and cumulative frequency of C-T and G-A misincorporations for individual Yana1.

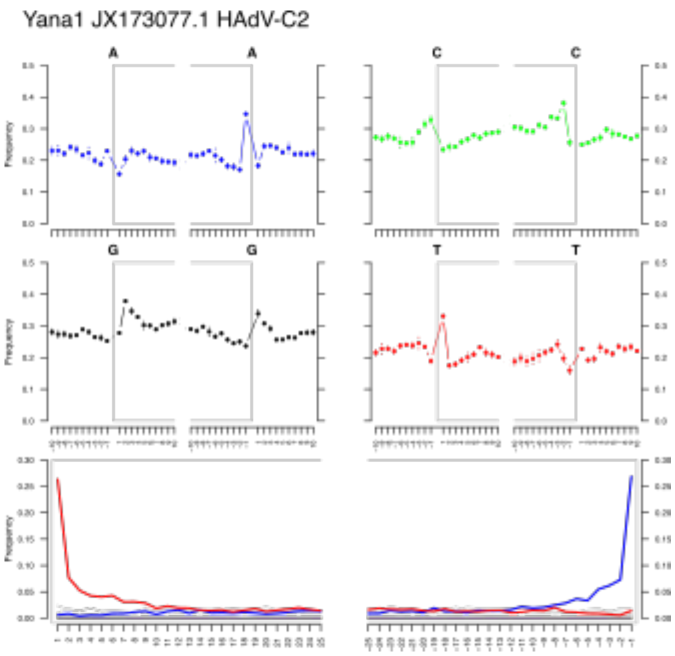

**Fig. S5. Adenovirus ancient DNA authentication for individual Yana2**. Top, nucleotide compositions of genomic regions for read ends and flanking positions. Bottom, cytosine to thymine (red) and guanine to adenine (blue) transitions.

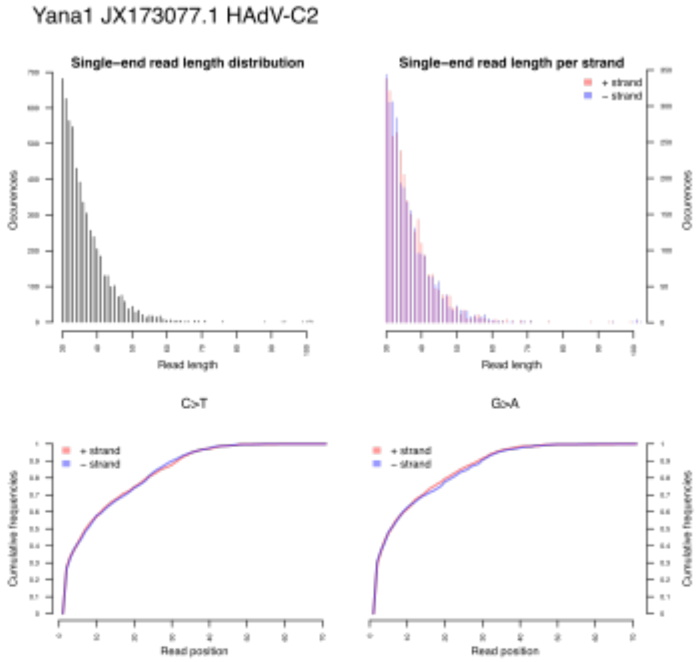

**Fig. S6. Yana2 read length distribution** and cumulative frequency of C-T and G-A misincorporations for individual Yana2.

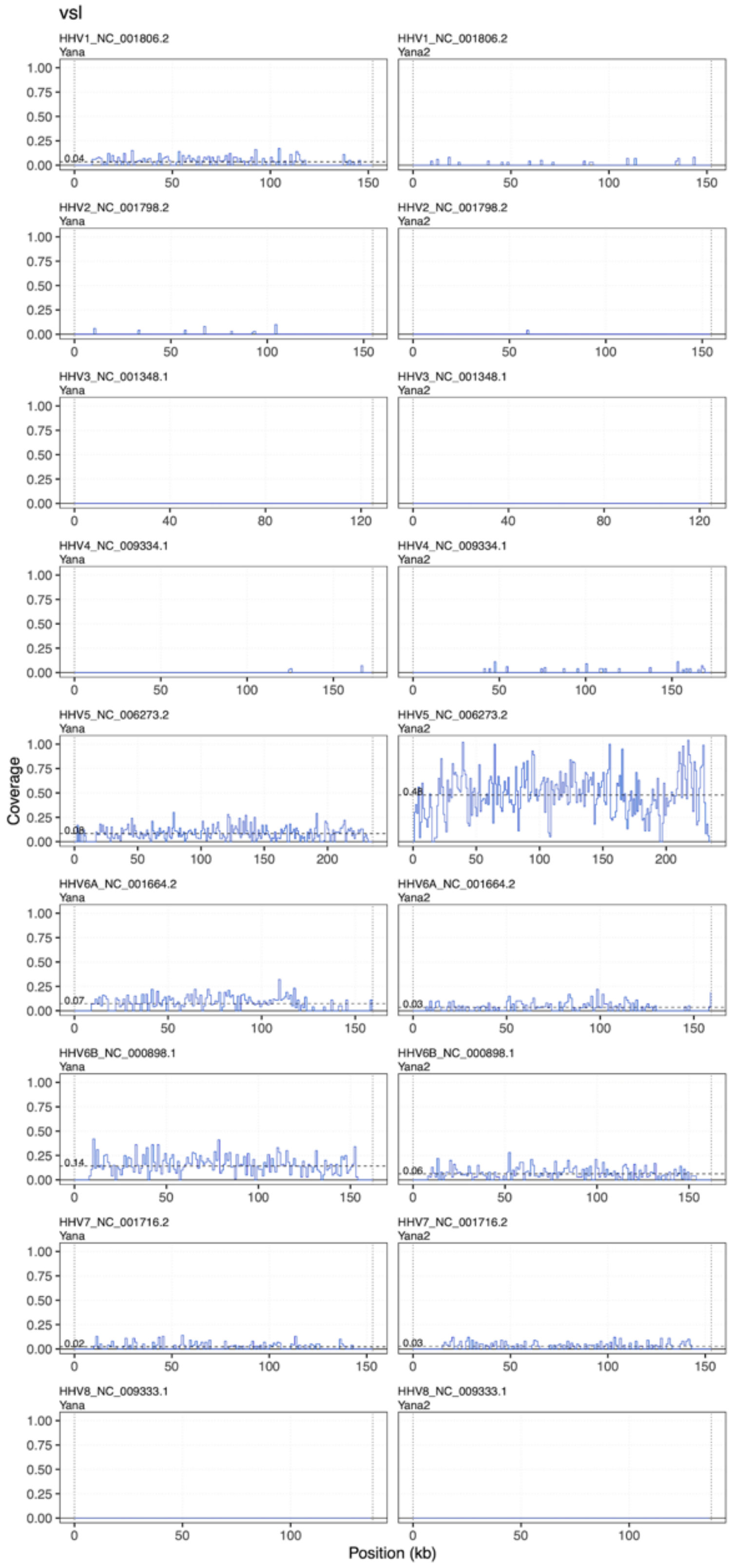

**Fig. S7. Genomic coverage for ancient herpesvirus.** Average genomic coverage across nine herpesvirus reference genomes (1 kbp windows) for the two ancient individuals, using modified 'very sensitive' preset and end-to-end alignment in bowtie2. Dashed line indicates genome-wide average coverage.

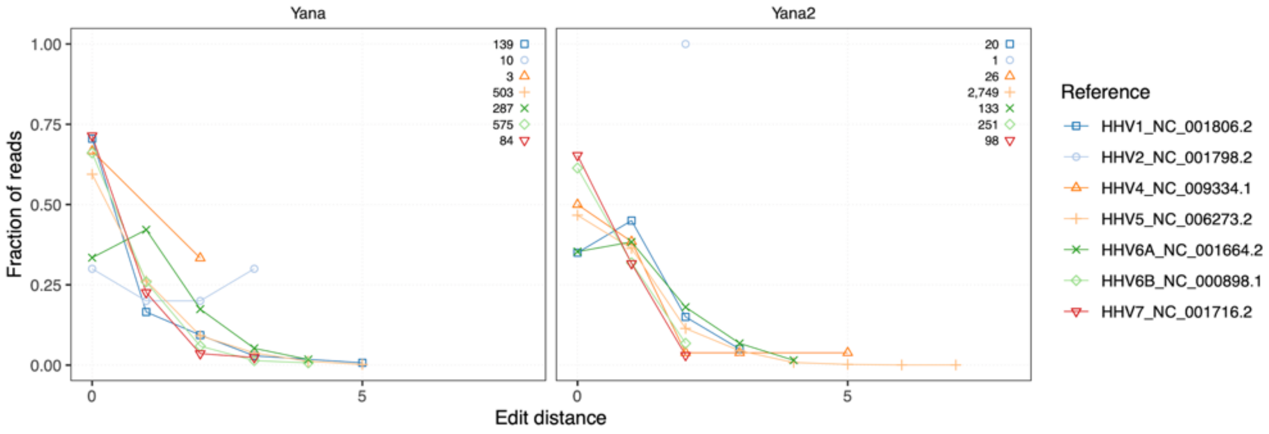

**Fig. S8. Read edit distance distributions for ancient herpesvirus.** Coloured lines indicate edit distance (number of mismatches) distributions for reads mapped against seven different herpesvirus reference genomes (modified 'very sensitive' preset, local alignment mode). Numbers indicate the total number of reads mapped after filters for each reference genome.

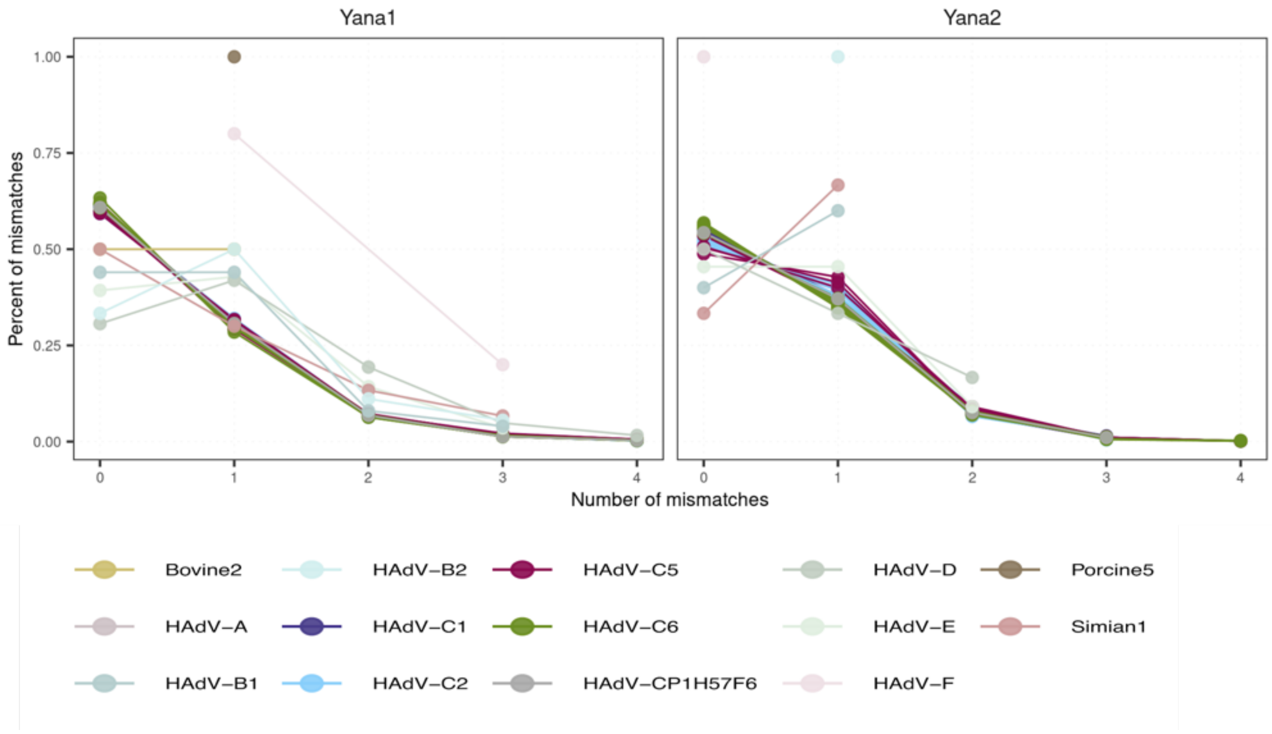

**Fig. S9. Read edit distance distributions for ancient adenovirus.** Coloured lines indicate distributions of edit distance (number of mismatches) for reads mapped against different HAdV reference genomes (modified 'very sensitive' preset, local alignment mode, after removal of duplicates and mapping quality below 30).

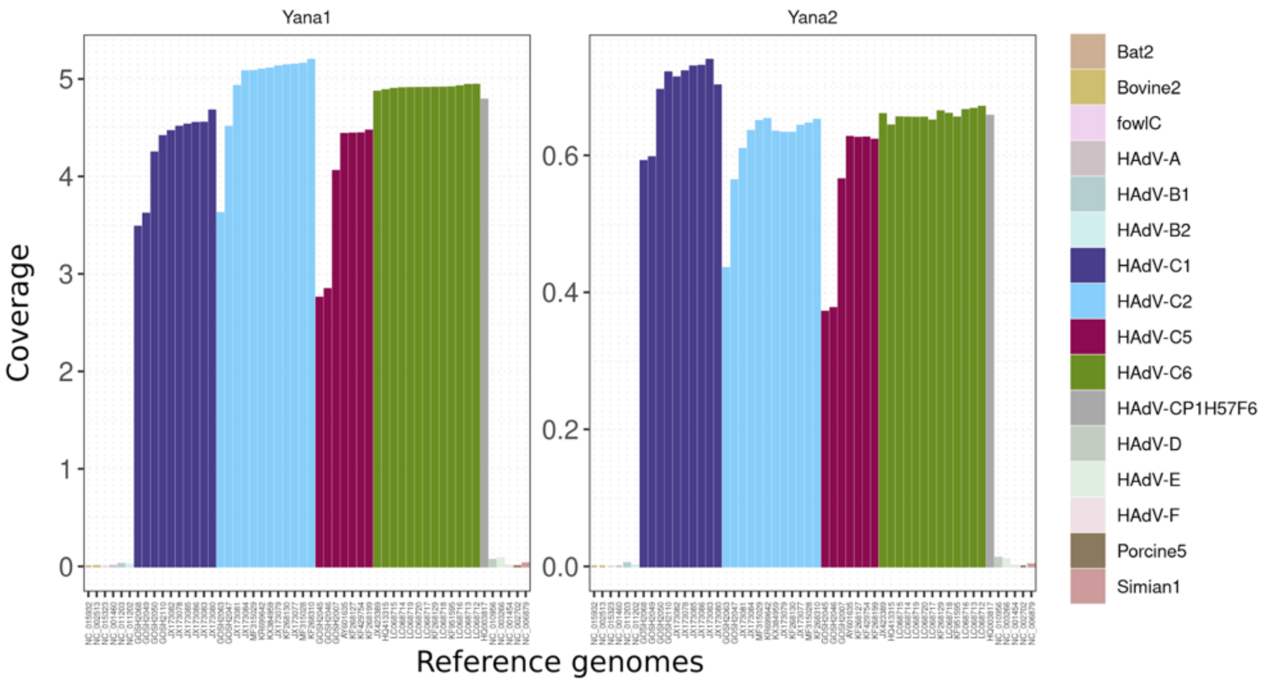

**Fig. S10. Genomic coverage for HAdV reference genomes.** Average coverage of adenovirus mappings for Yana1 (left) and Yana2 (right). Different HAdV species as well as HAdV-C genotypes are indicated by coloured bars.

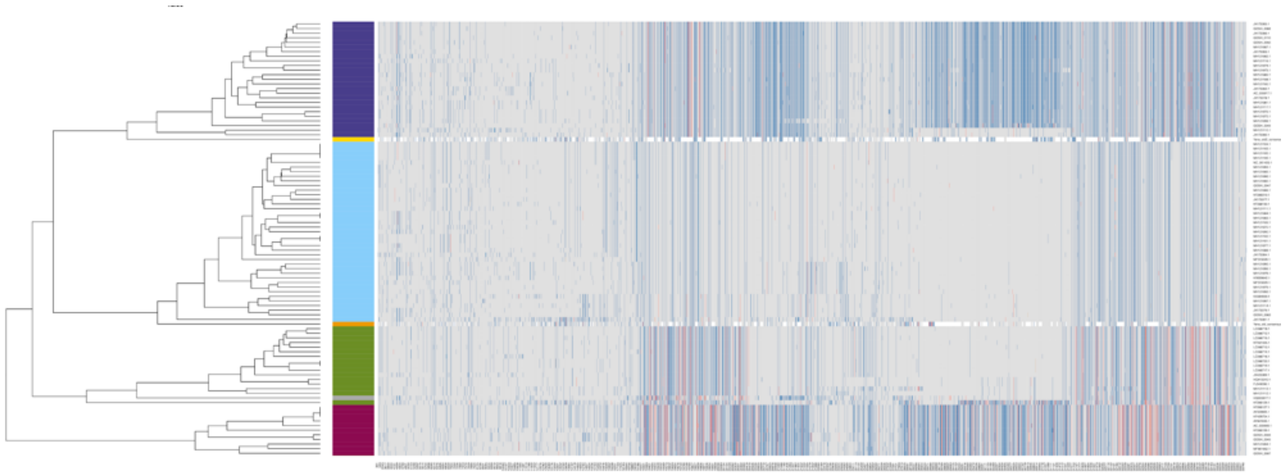

**Fig. S11. Genetic diversity of HAdV-C.** Heatmap visualization of genetic variants in the HAdV-C multiple sequence alignment, for all 3,634 variants including missing genotypes in the ancient genomes. Genotypes for SNPs are coloured according to minor allele frequency. Dendrogram shows the result of hierarchical clustering of modern and ancient genomes based on observed genotypes, with major types indicated by coloured bars.

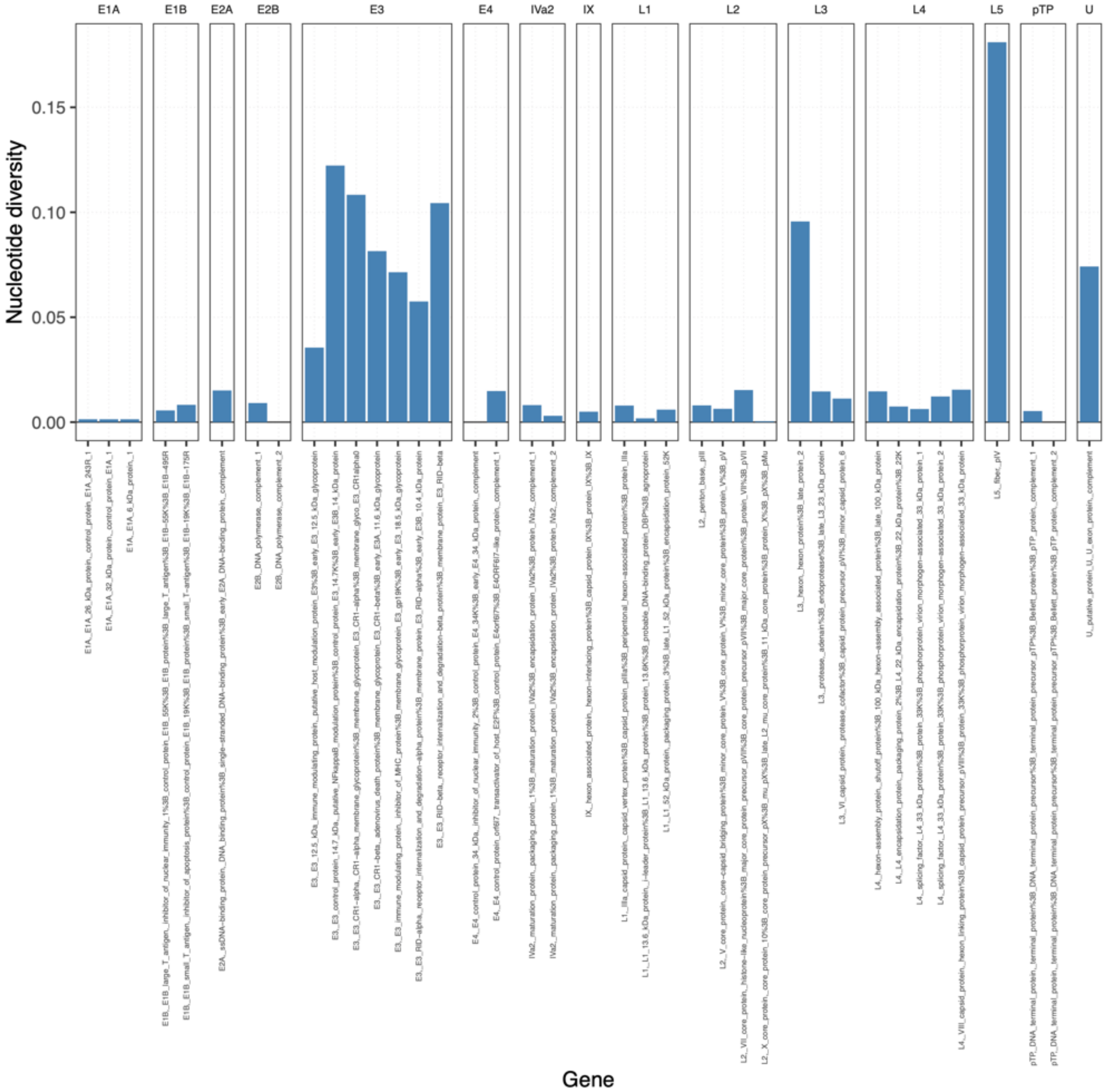

**Fig. S12. Genetic diversity of HAdV-C genes.** Bar plot showing nucleotide diversity for HAdV-C genes, grouped by transcript unit.

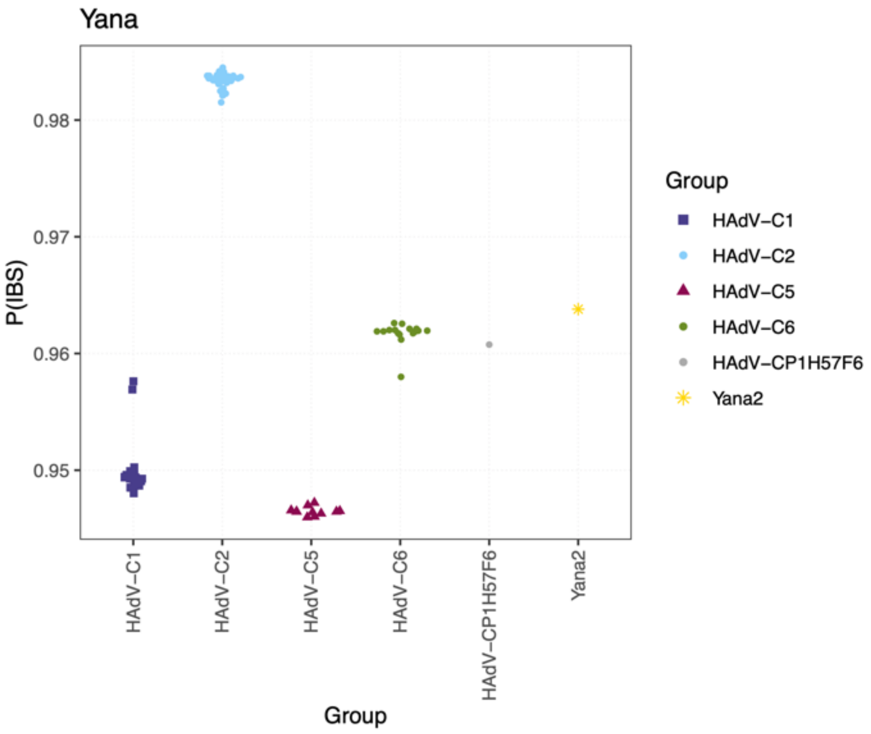

**Fig. S13. Genetic affinities of Yana1 virus genome.** Plot shows overall genetic similarity measured as a fraction of genomic sites shared identical-by-state (IBS), between Yana1 and other genomes. HAdV-C types are indicated by different plot symbols and colours.

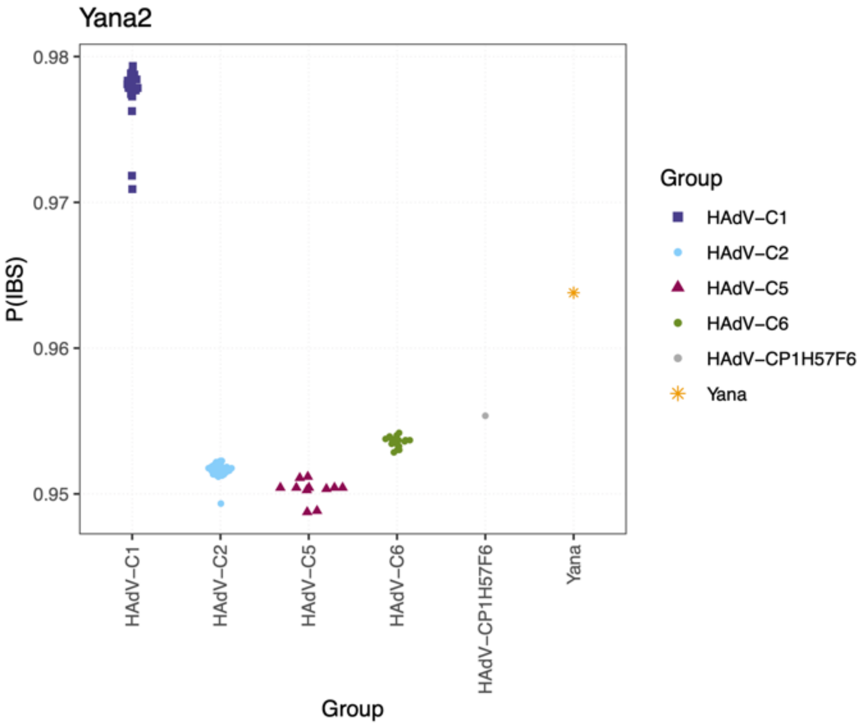

**Fig. S14. Genetic affinities of Yana2 virus genome.** Plot shows overall genetic similarity measured as a fraction of genomic sites shared identical-by-state (IBS), between Yana2 and other genomes. HAdV-C types are indicated by different plot symbols and colours.

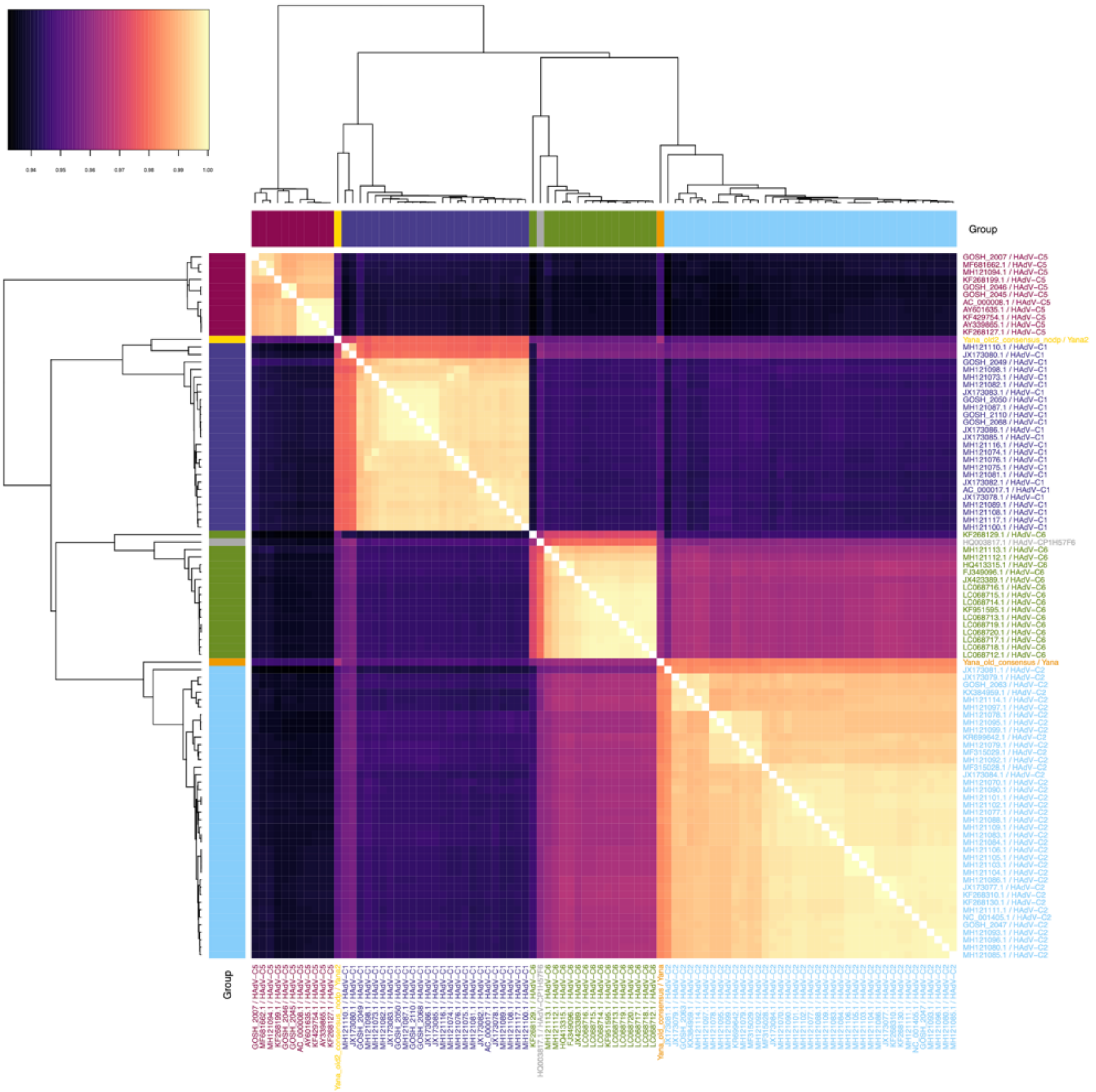

**Fig. S15. Genetic similarity of HAdV-C genomes.** Heatmap showing overall genetic similarity between pairs of virus genomes. Dendrogram shows the result of hierarchical clustering based on nucleotide identity, and coloured bars indicate HAdV-C types.

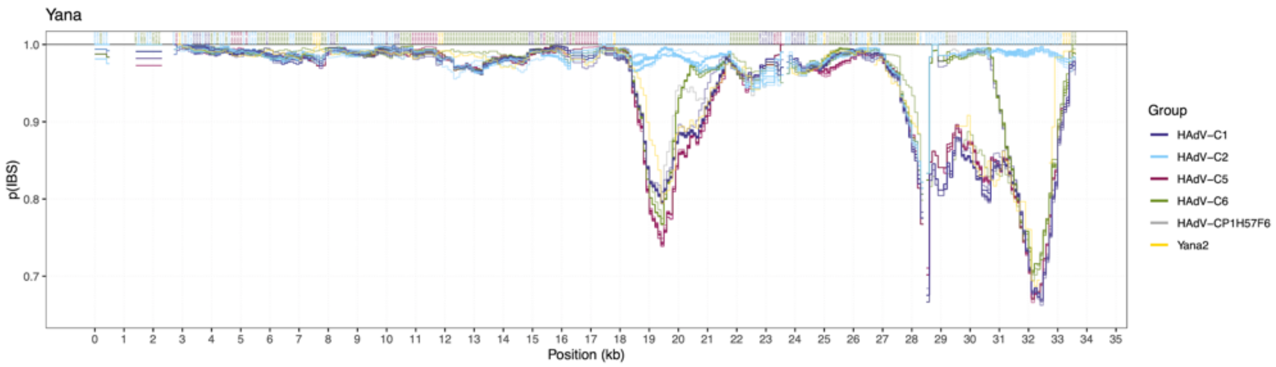

**Fig. S16. Pairwise genetic similarity along Yana1 genome.** Coloured lines indicate average nucleotide identity (ANI) in 1kb sliding windows (100 bp step size) between the Yana1 ancient genome and all other genomes. For each window, the accession for the most similar comparison genome is indicated along the top of the plot.

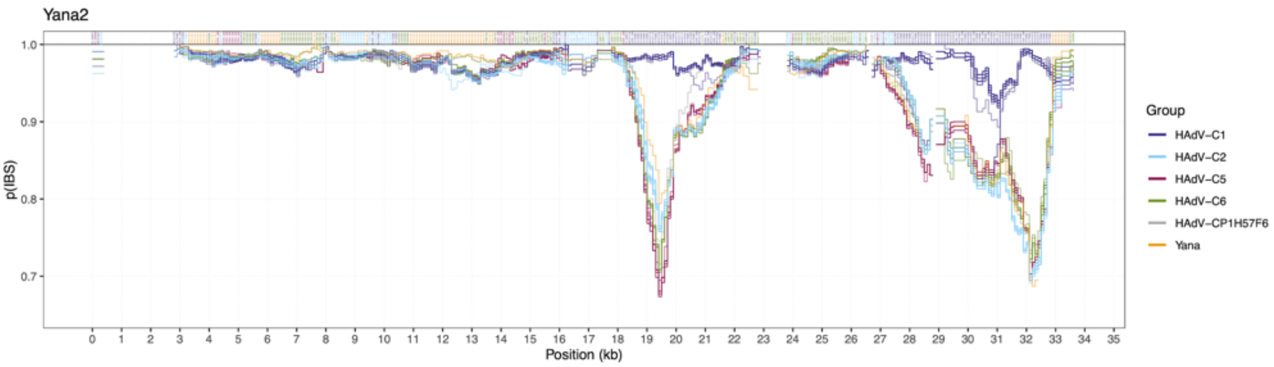

**Fig. S17. Pairwise genetic similarity along Yana2 genome.** Coloured lines indicate average nucleotide identity (ANI) in 1kb sliding windows (100 bp step size) between the Yana2 ancient genome and all other genomes. For each window, the accession for the most similar comparison genome is indicated along the top of the plot.

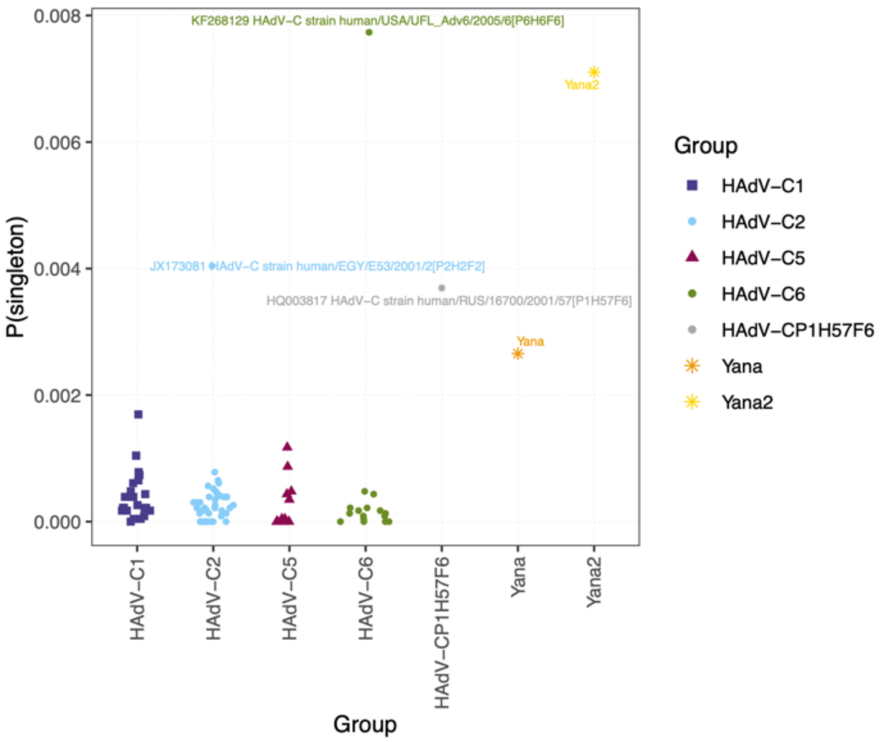

**Fig. S18. Private genetic variants in HAdV-C genomes.** Plot shows the fraction of private variants among all sites for HAdV-C genomes. The respective HAdV-C types are indicated by different plot symbols and colours, and genomes with high rates of singleton variants are labelled.

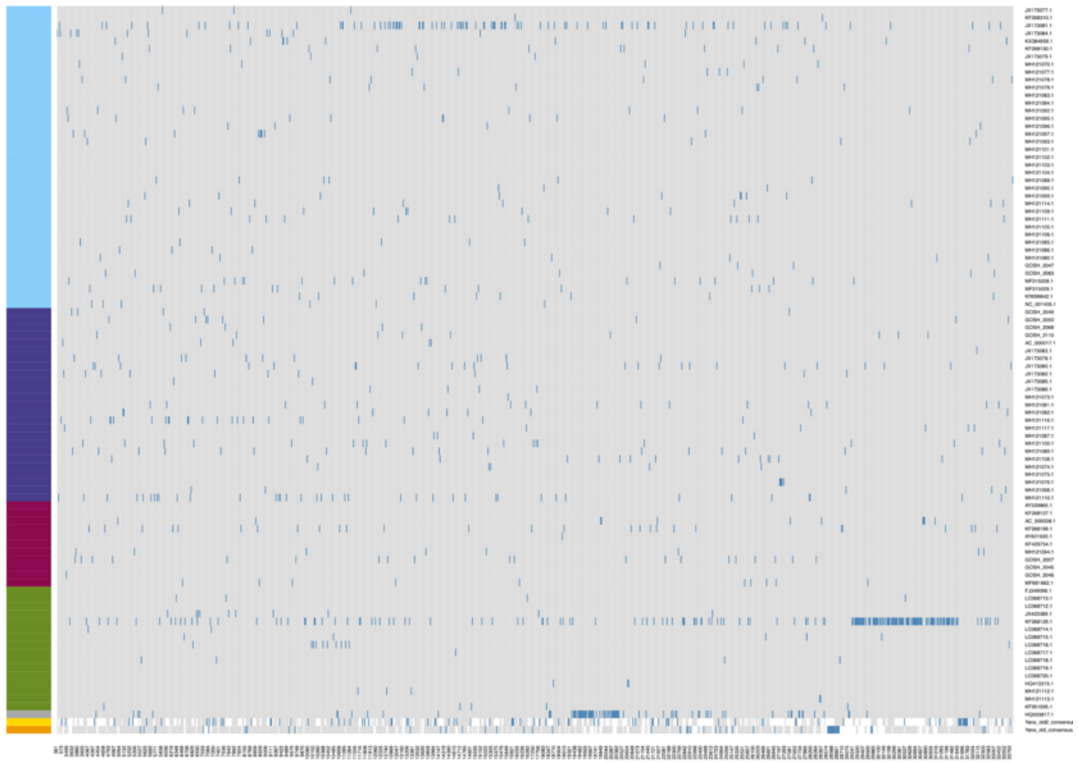

**Fig. S19. Genomic distribution of singleton variants.** Heatmap showing genomic distributions of singleton variants (highlighted in blue) across all HAdV-C genomes.

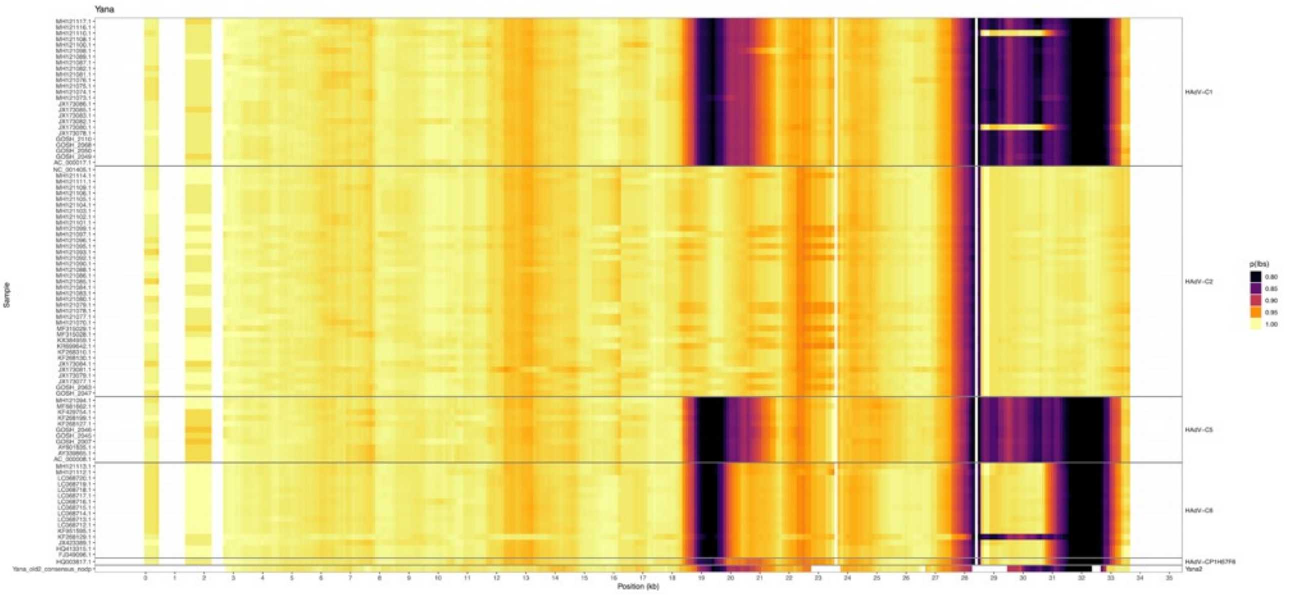

**Fig. S20. Pairwise genetic similarity for Yana1 genome.** Heatmap showing average nucleotide identity (ANI) in 1kb sliding windows (100 bp step size) between the Yana1 ancient genome and all other HAdV-C genomes and Yana2, grouped by type. Recombinant sequences can be identified by local within-type differences in similarity against Yana1.

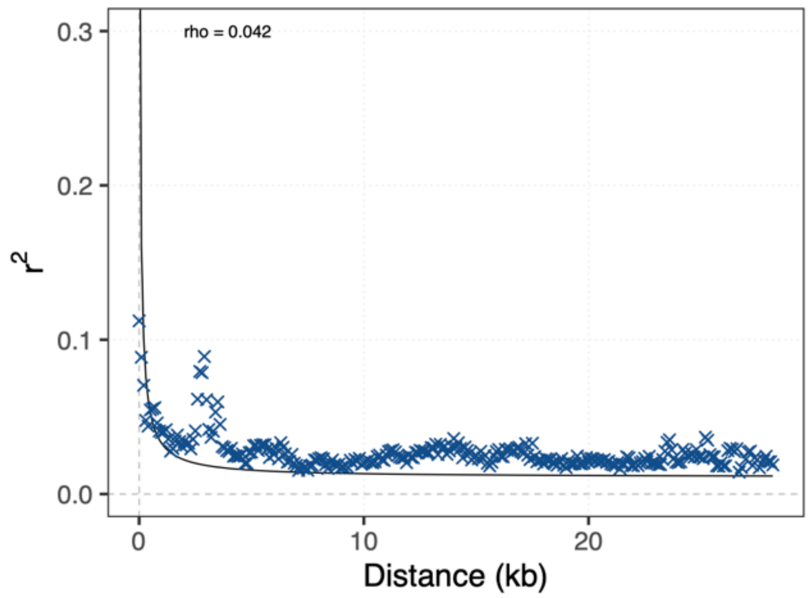

**Fig. S21. Decay of linkage disequilibrium.** LD decay curve showing average pairwise $r^2$ as a function of the distance between SNPs (bins of 100 bp), for genomic regions excluding L3, E3 and L5 transcript units. The black curve indicates the theoretically expected decay from the estimated population recombination rate (indicated in plot).

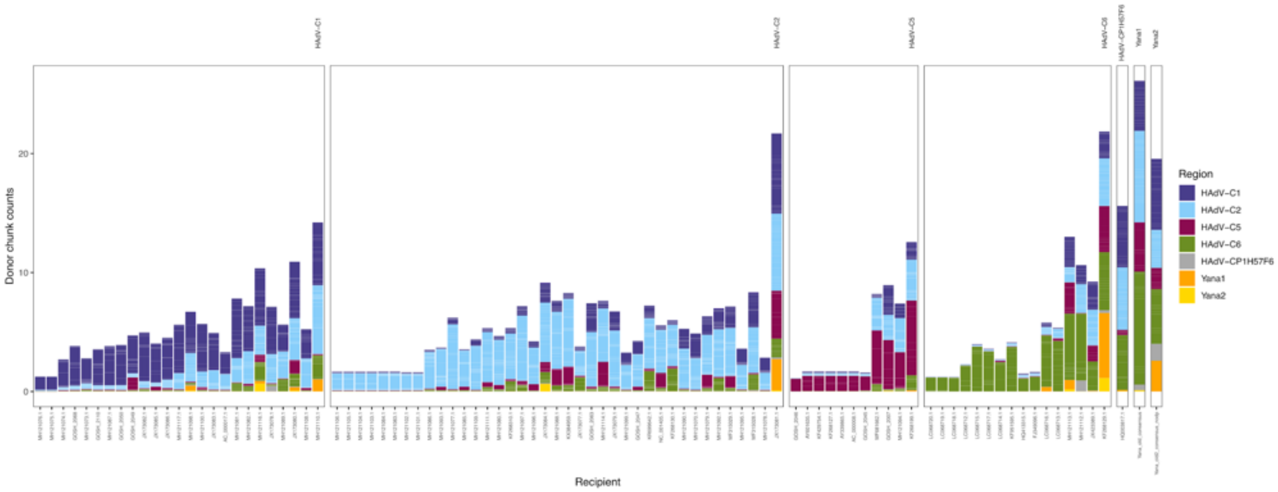

**Fig. S22. Chromopainter, chunk counts,** all against all

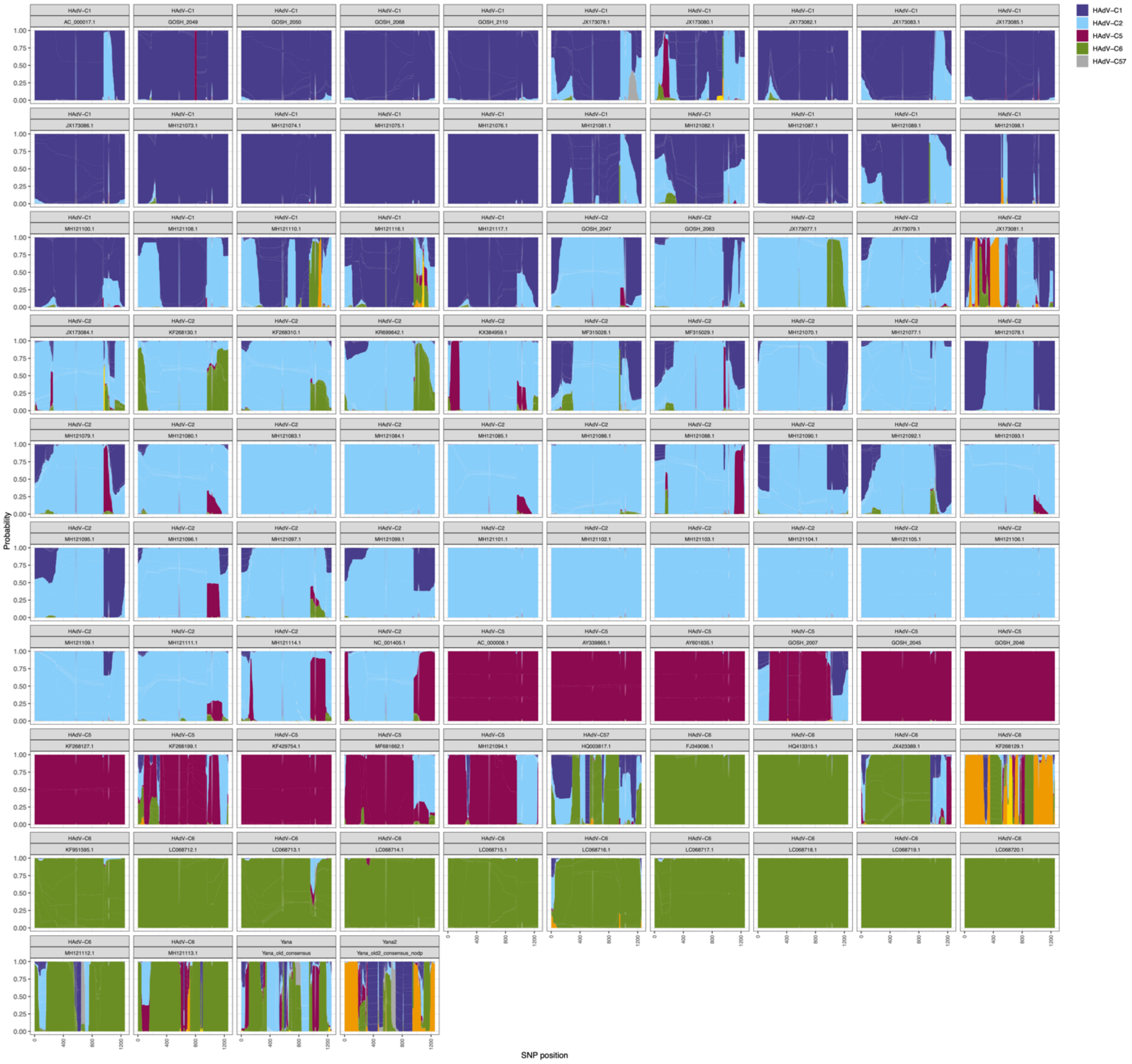

**Fig. S23. Chromopainter, local ancestry,** all against all.

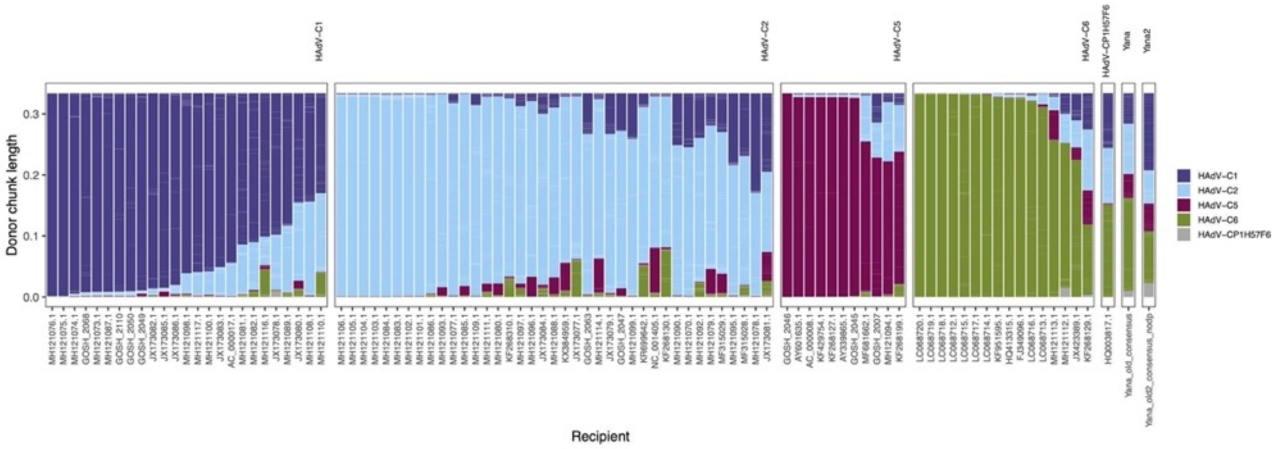

**Fig. S24. Chromopainter, chunk length**, all but ancient samples donate.

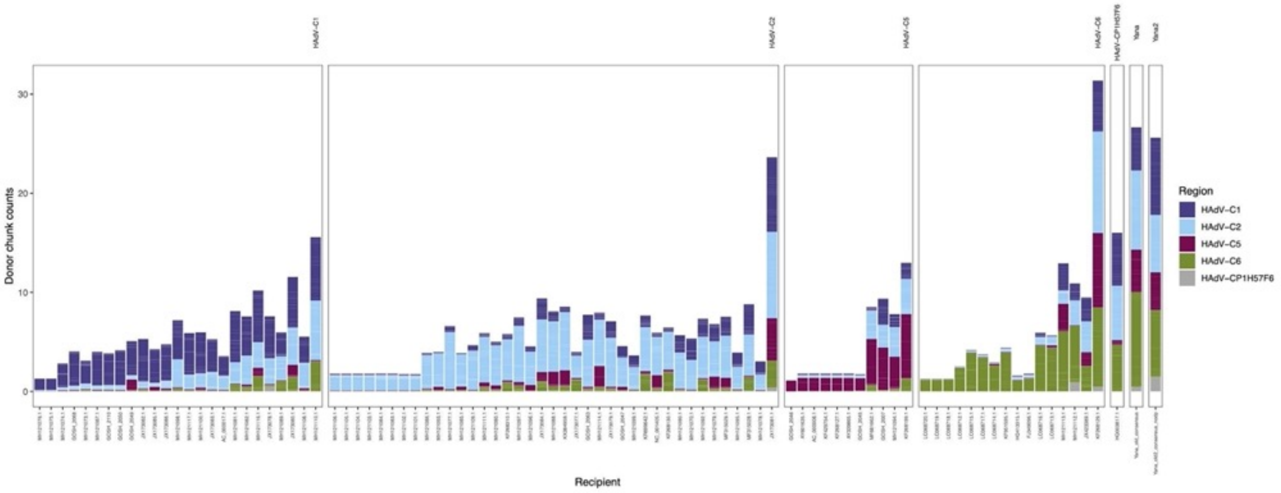

**Fig. S25. Chromopainter, chunk counts**, all but ancient samples donate.

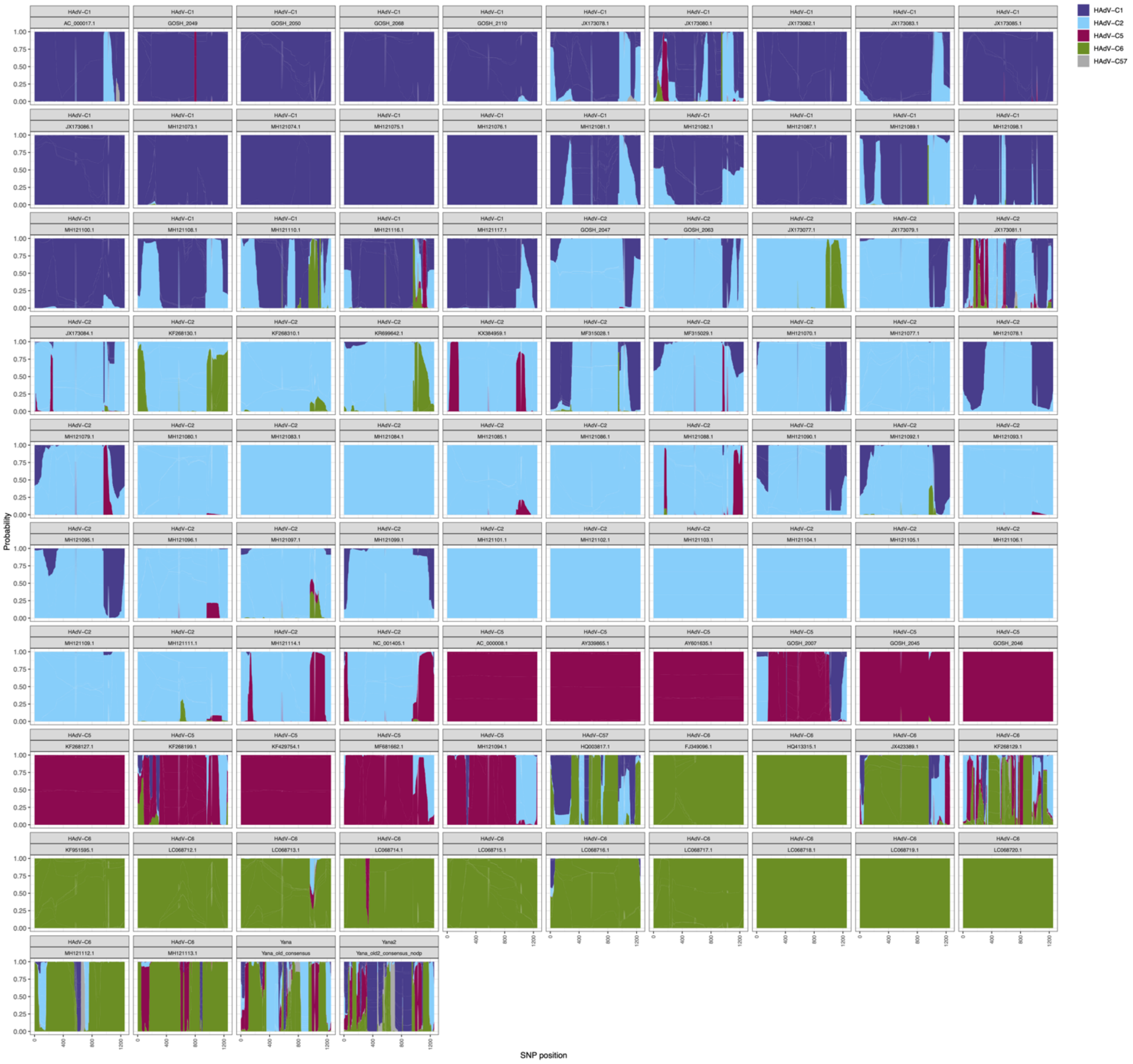

**Fig. S26. Chromopainter, local ancestry**, all but ancient samples donate.

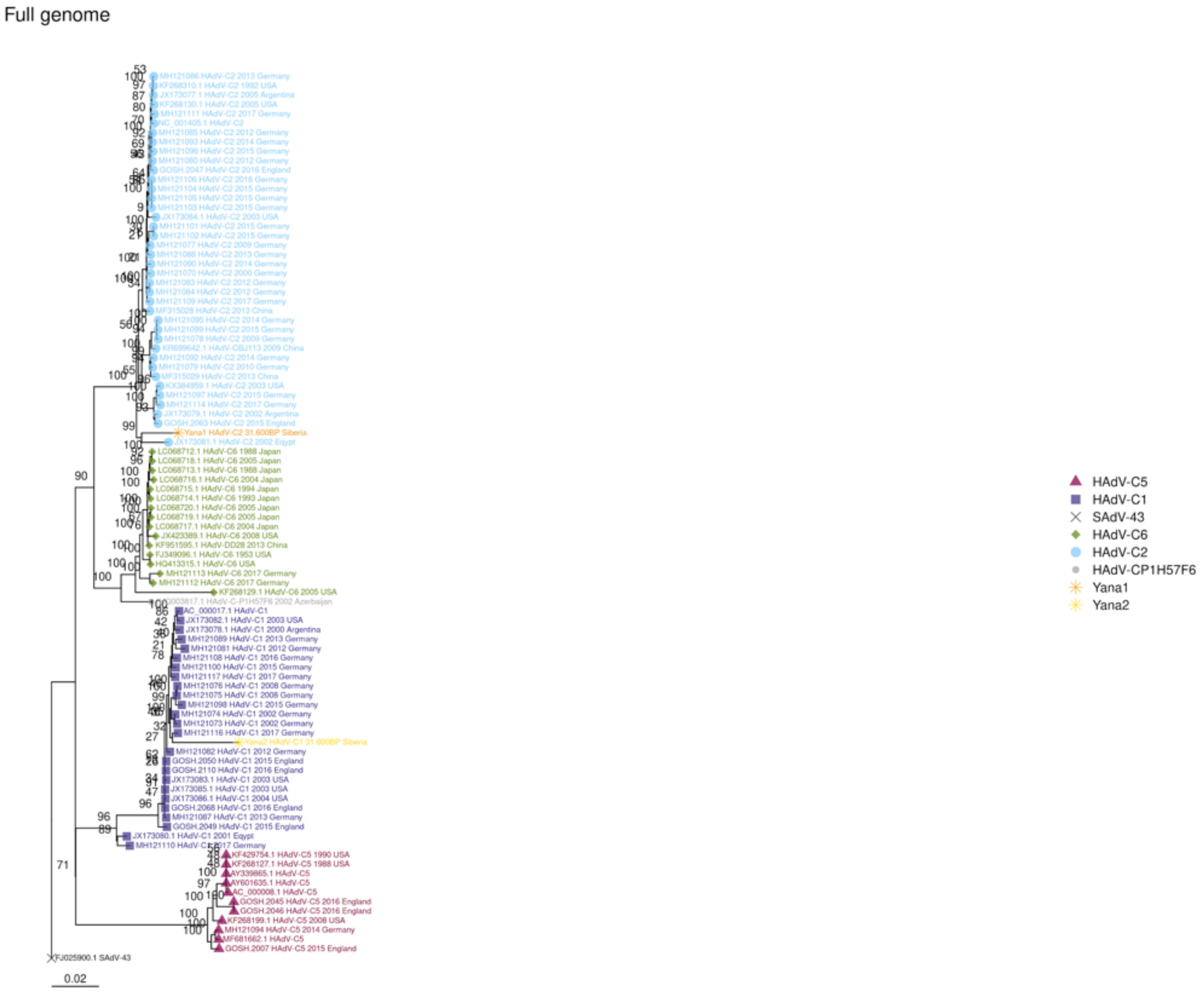

**Fig. S27. Whole genome maximum likelihood tree**, model GTR+G+I, outgroup branch removed for visualization.

E1A

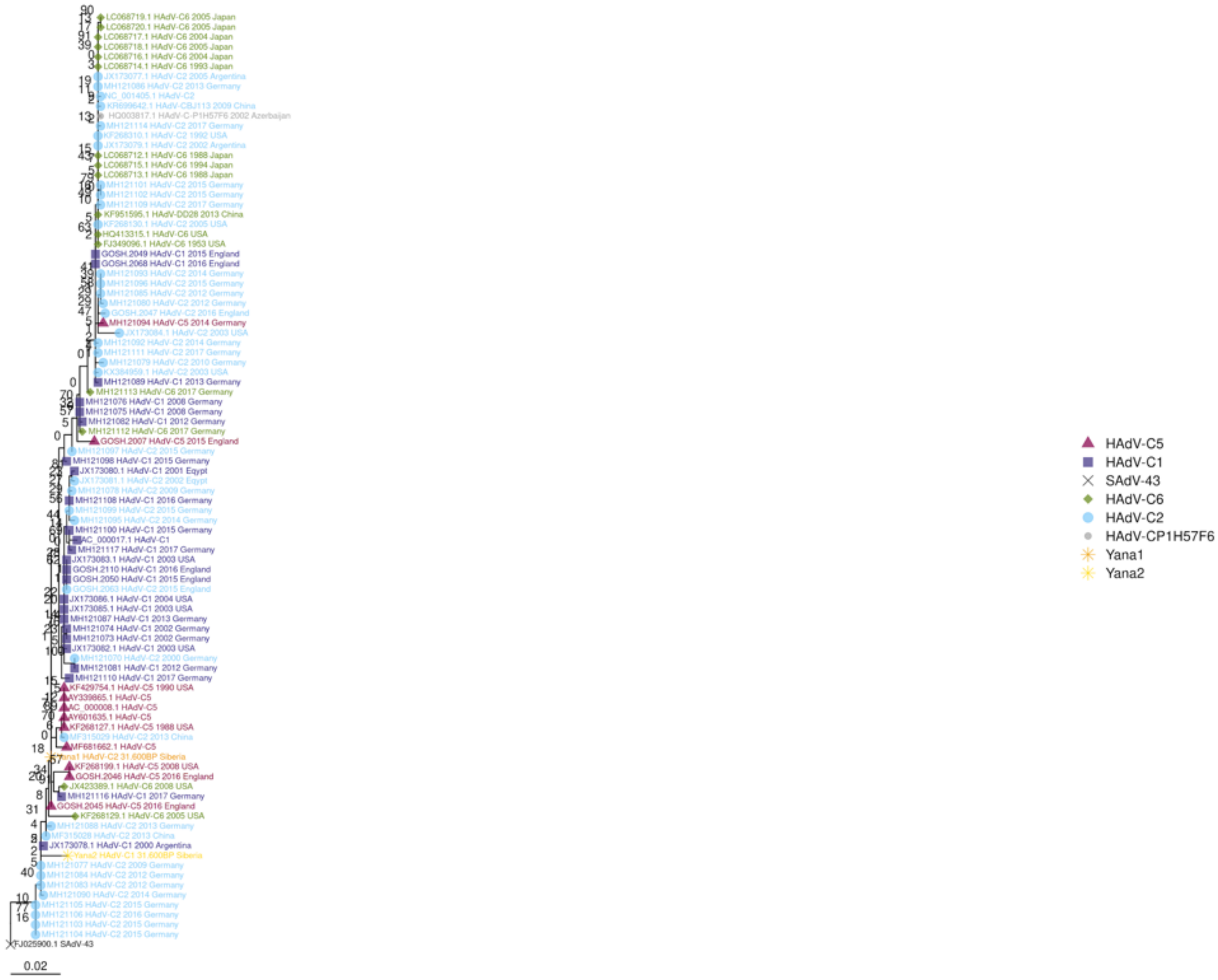
HAdV-C5
HAdV-C1
SAdV-43
HAdV-C6
HAdV-C2
HAdV-CP1H57F6
Yana1
Yana2
0.02

E1B

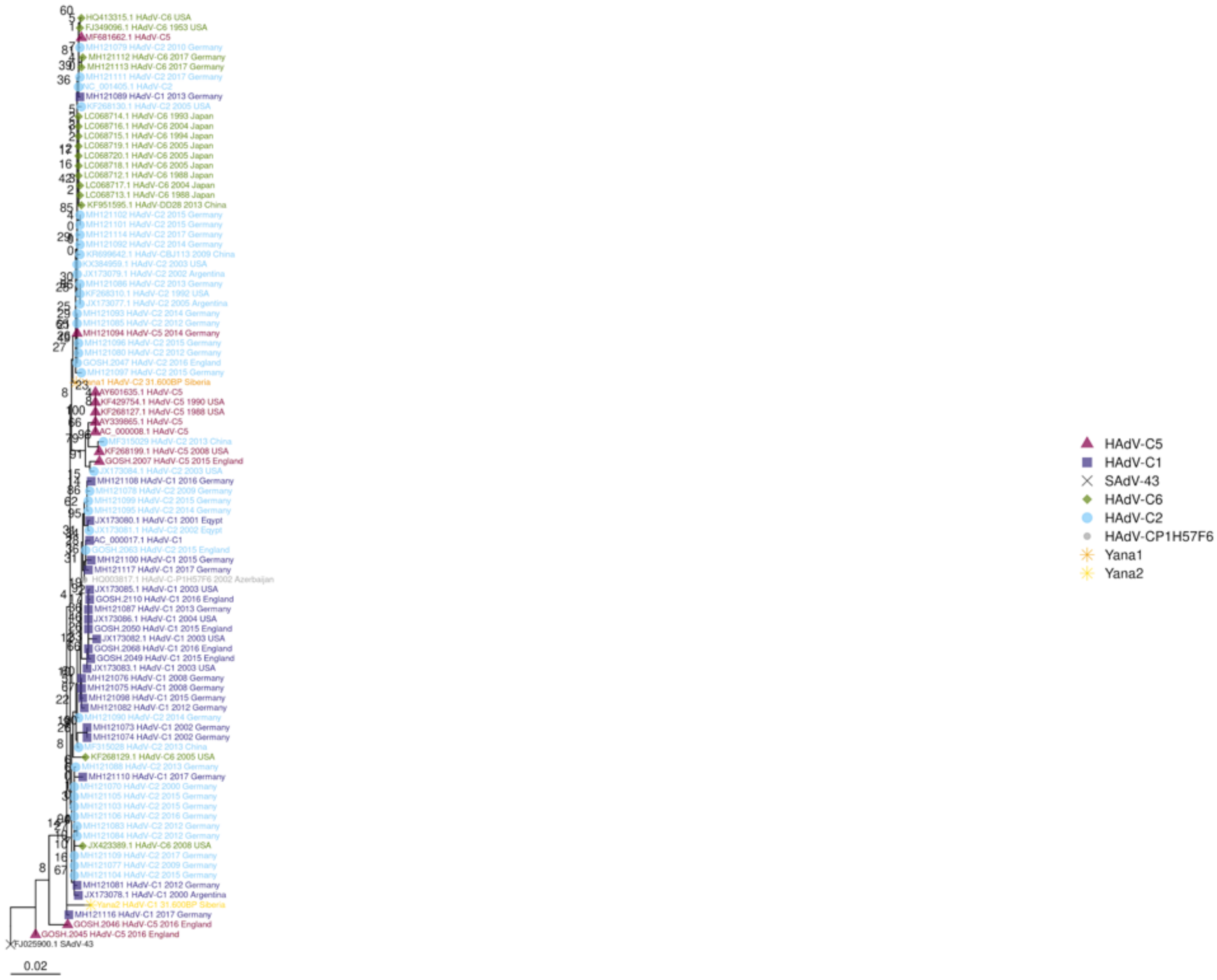
HAdV-C5
HAdV-C1
SAdV-43
HAdV-C6
HAdV-C2
HAdV-CP1H57F6
Yana1
Yana2
0.02

E2A

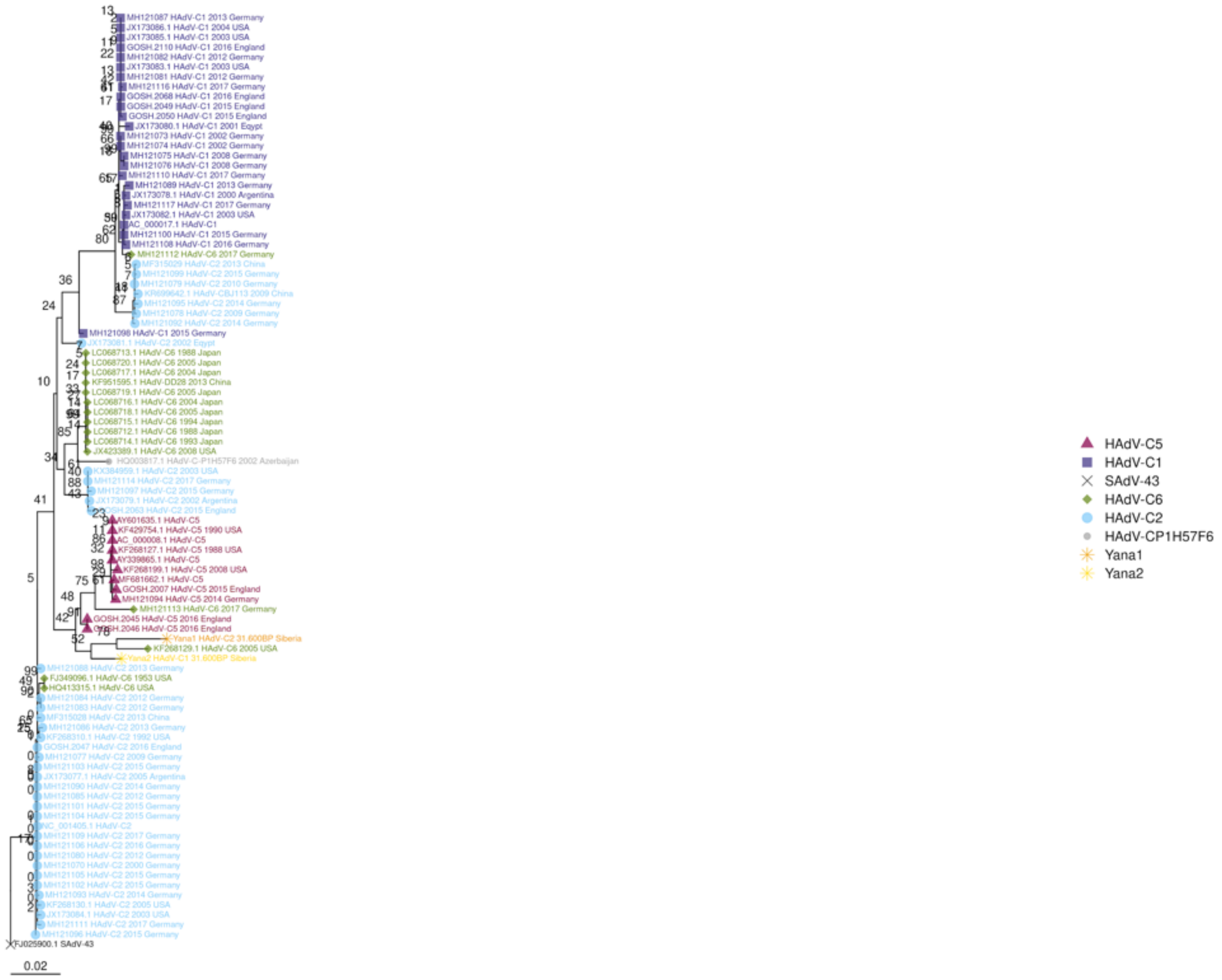

0.02
HAdV-C5
HAdV-C1
SAdV-43
HAdV-C6
HAdV-C2
HAdV-CP1H57F6
Yana1
Yana2

E2B

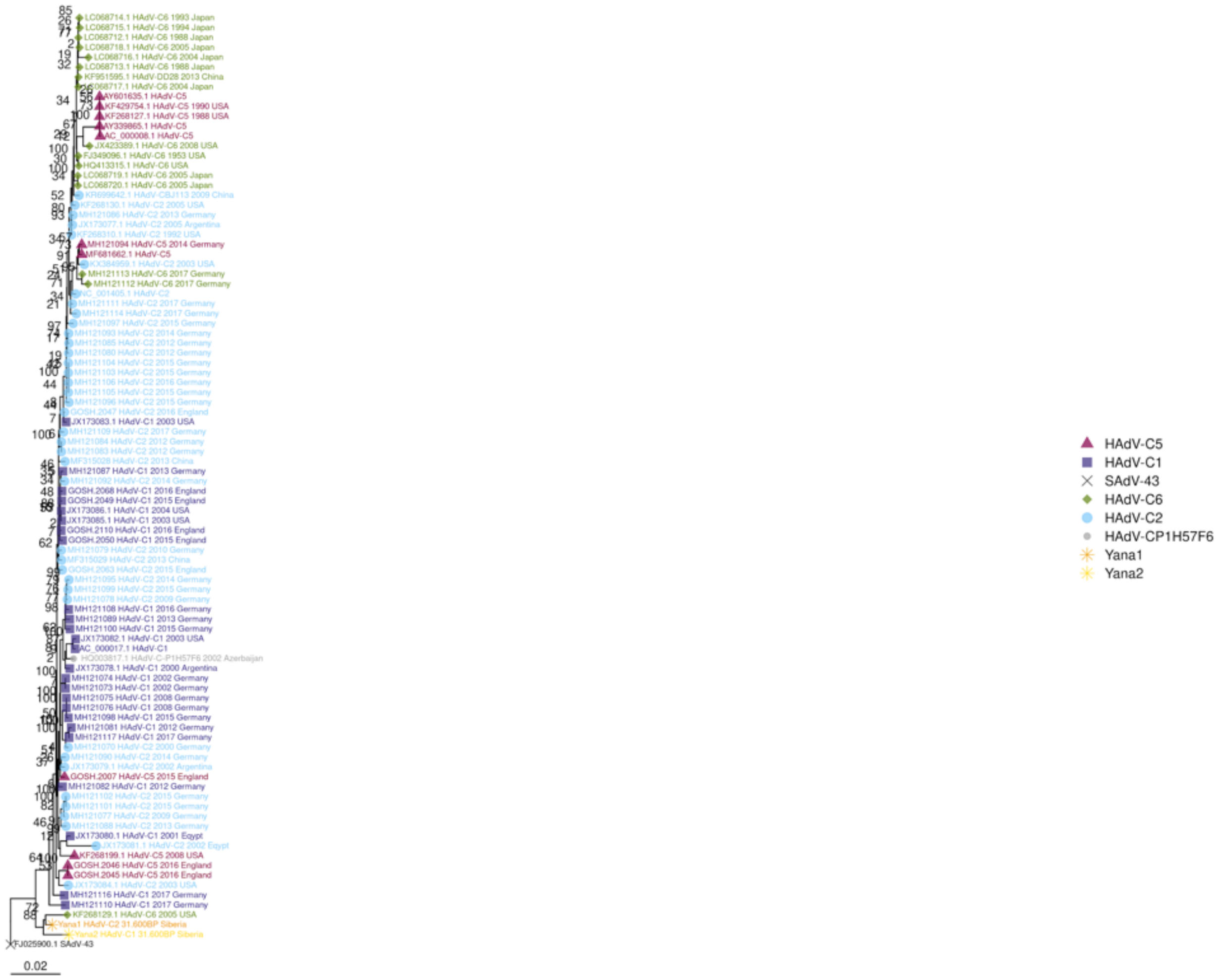
HAdV-C5
HAdV-C1
SAdV-43
HAdV-C6
HAdV-C2
HAdV-CP1H57F6
Yana1
Yana2
0.02

E3

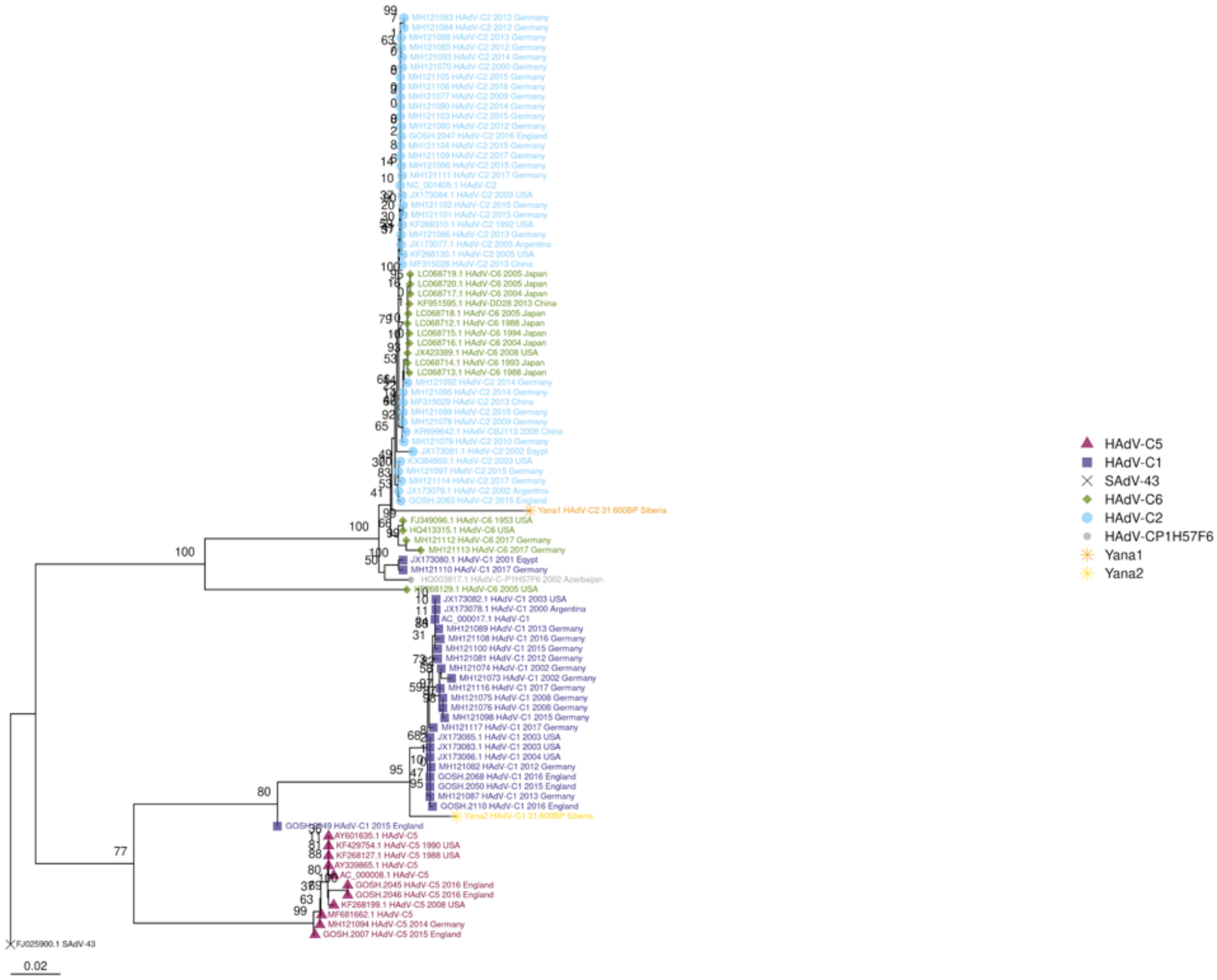
HAdV-C5
HAdV-C1
SAdV-43
HAdV-C6
HAdV-C2
HAdV-CP1H57F6
Yana1
Yana2
0.02

E4

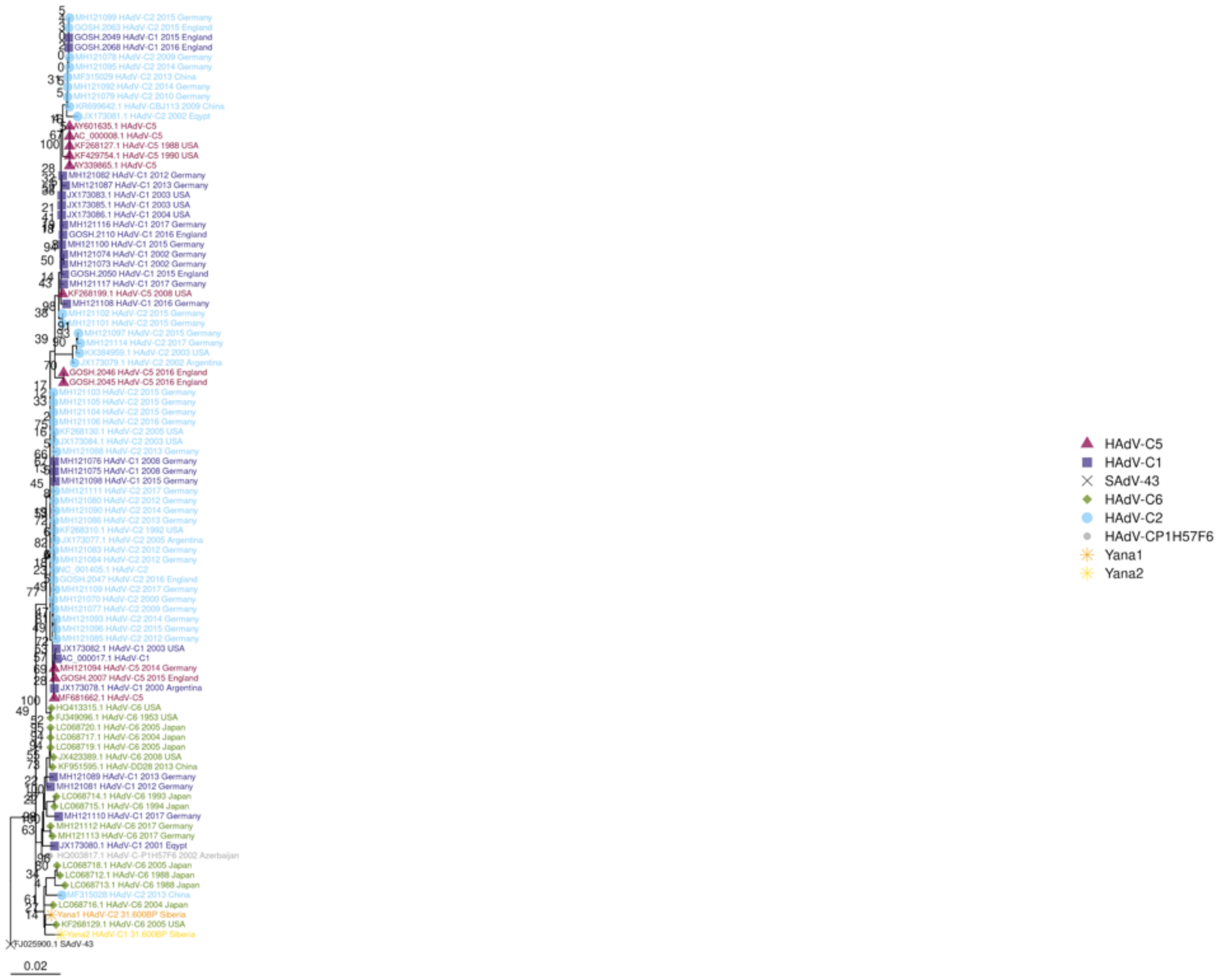
HAdV-C5
HAdV-C1
SAdV-43
HAdV-C6
HAdV-C2
HAdV-CP1H57F6
Yana1
Yana2
0.02

IVa2

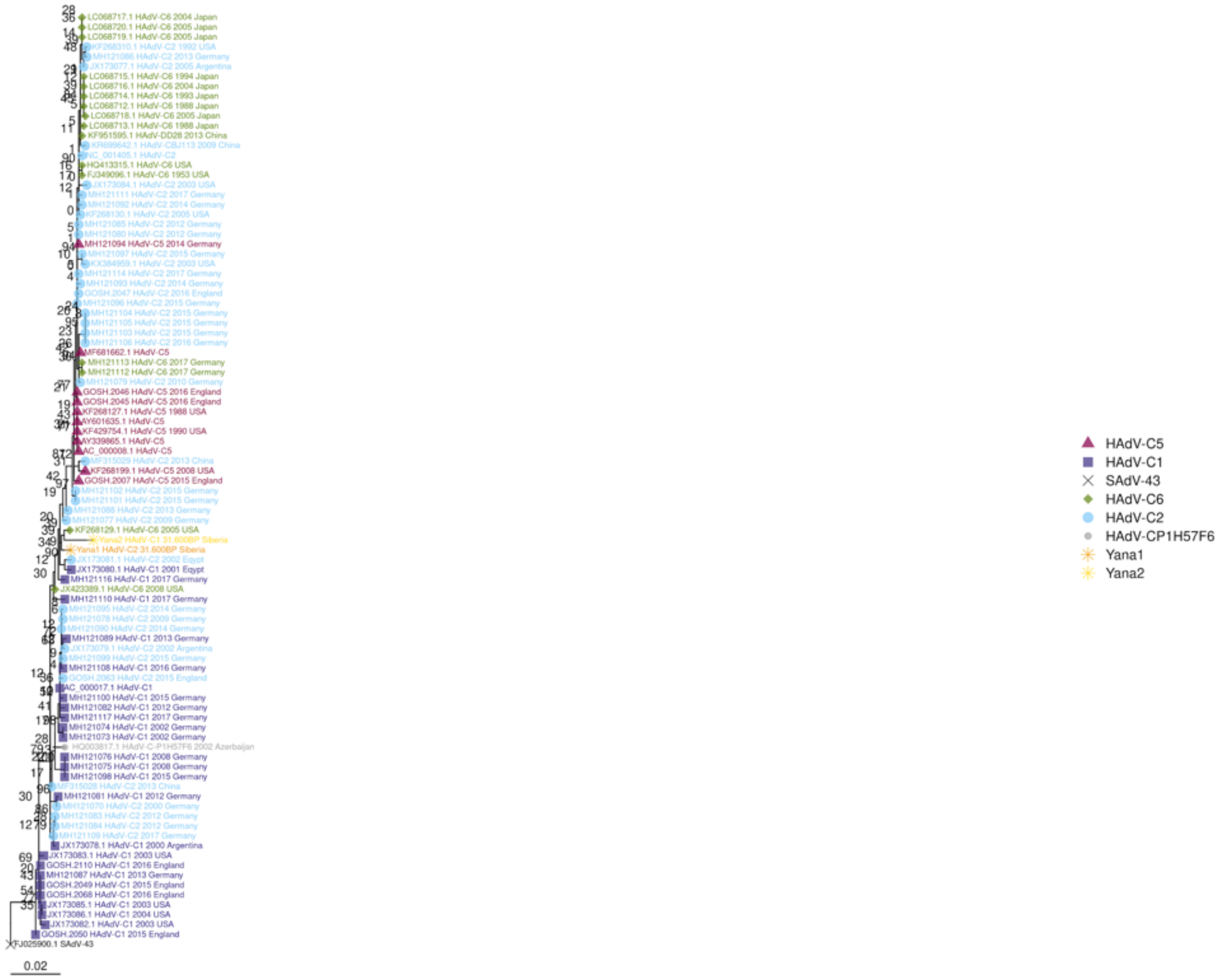
HAdV-C5
HAdV-C1
SAdV-43
HAdV-C6
HAdV-C2
HAdV-CP1H57F6
Yana1
Yana2
0.02

IX

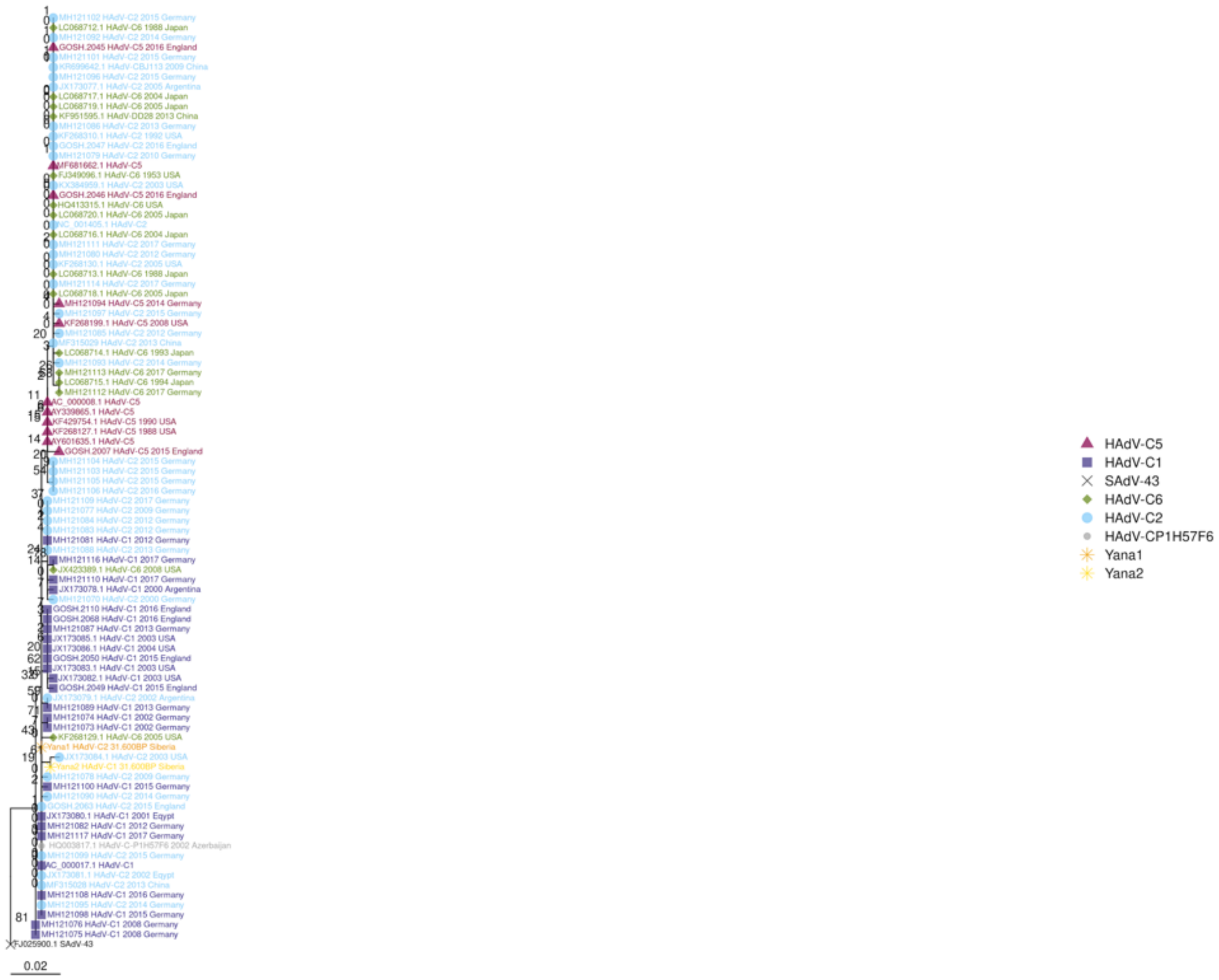
HAdV-C5
HAdV-C1
SAdV-43
HAdV-C6
HAdV-C2
HAdV-CP1H57F6
Yana1
Yana2
0.02

L1

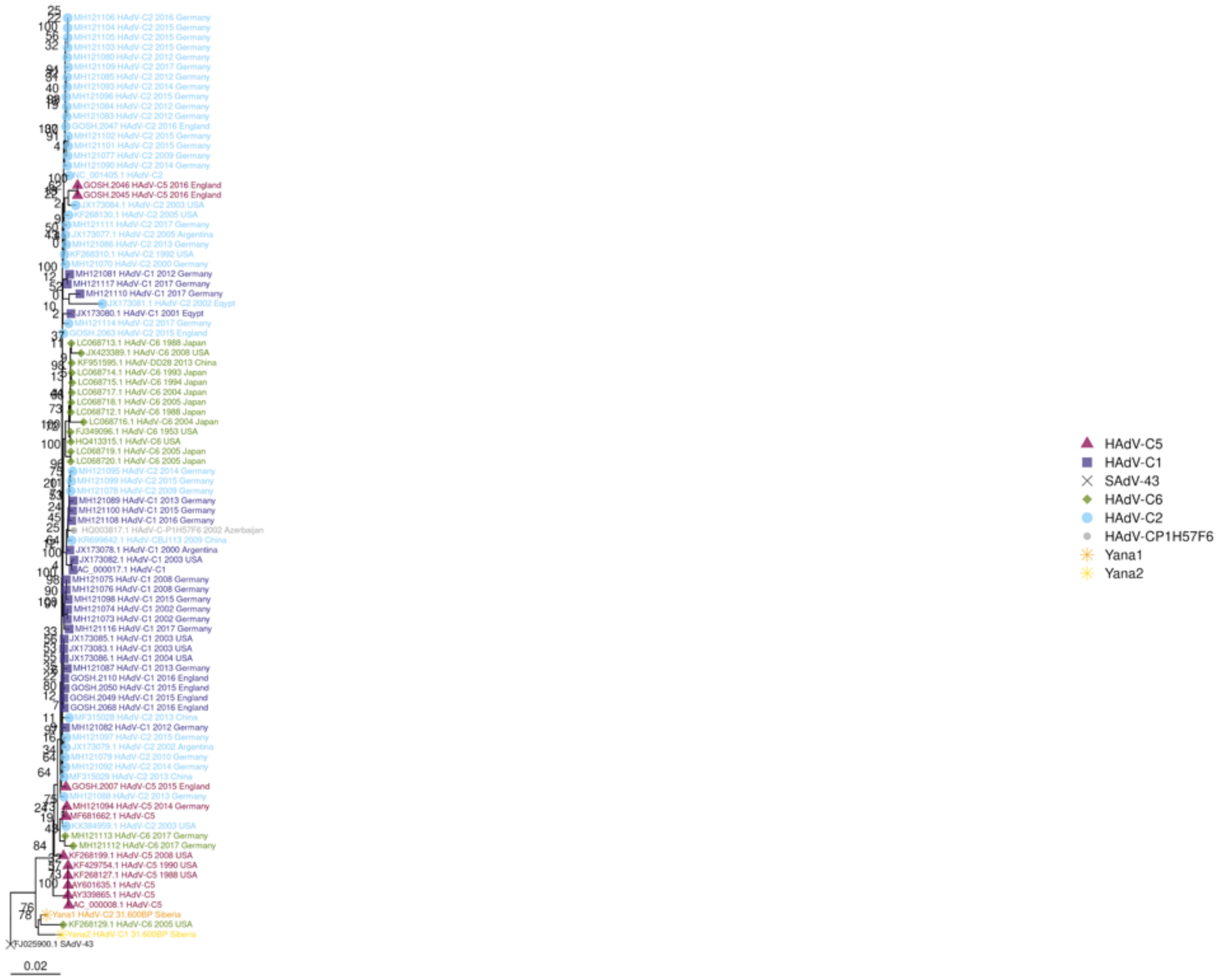
HAdV-C5
HAdV-C1
SAdV-43
HAdV-C6
HAdV-C2
HAdV-CP1H57F6
Yana1
Yana2
0.02

L2

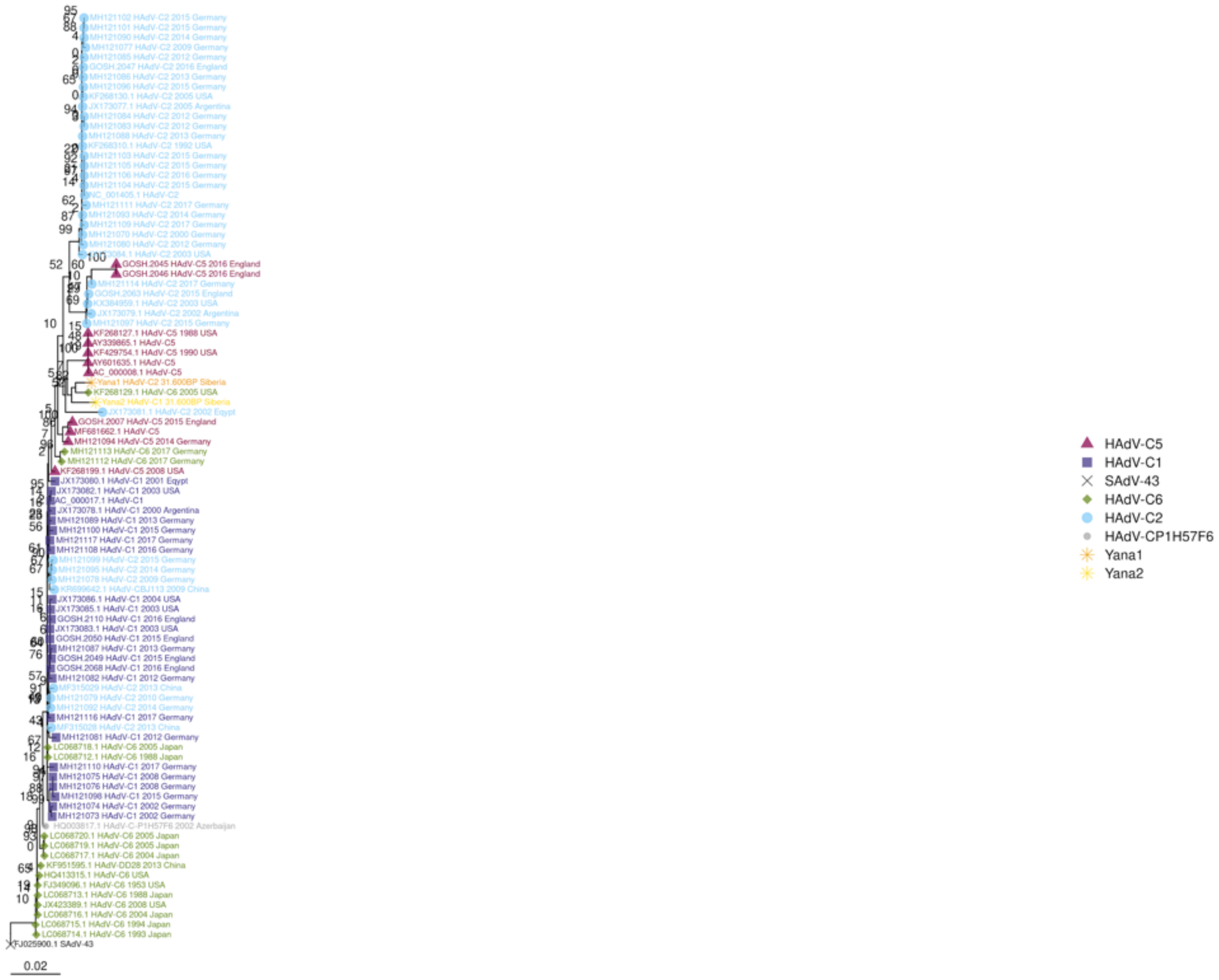

HAdV-C5
HAdV-C1
SAdV-43
HAdV-C6
HAdV-C2
HAdV-CP1H57F6
Yana1
Yana2
0.02

L3

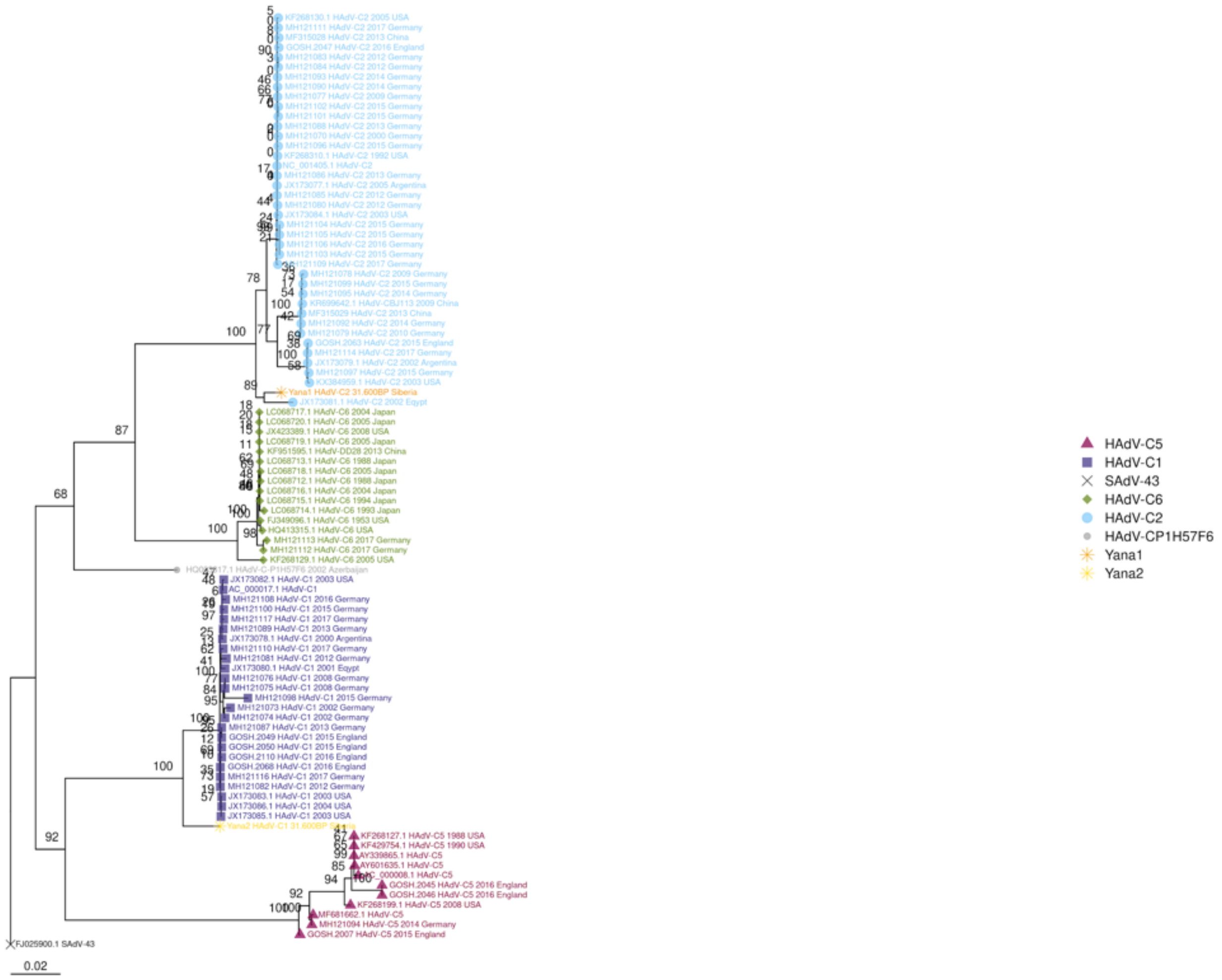

HAdV-C5
HAdV-C1
SAdV-43
HAdV-C6
HAdV-C2
HAdV-CP1H57F6
Yana1
Yana2
0.02

L4

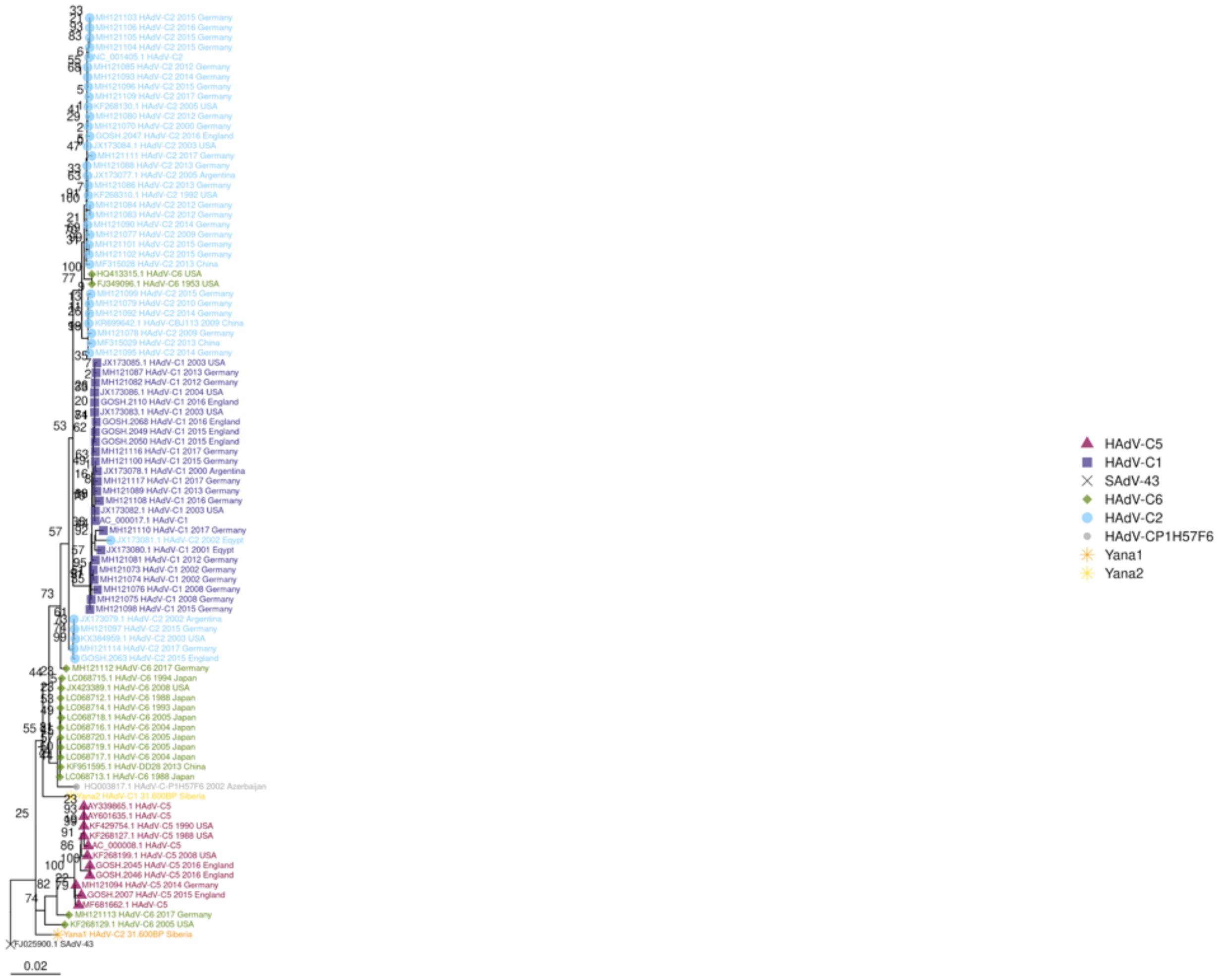
MH121103 HAdV-C2 2015 Germany
MH121106 HAdV-C2 2016 Germany
MH121105 HAdV-C2 2015 Germany
MH121104 HAdV-C2 2015 Germany
NC_001405.1 HAdV-C2
MH121085 HAdV-C2 2012 Germany
MH121093 HAdV-C2 2014 Germany
MH121096 HAdV-C2 2015 Germany
MH121109 HAdV-C2 2017 Germany
KF268130.1 HAdV-C2 2005 USA
MH121080 HAdV-C2 2012 Germany
MH121070 HAdV-C2 2000 Germany
GOSH.2047 HAdV-C2 2016 England
JX173084.1 HAdV-C2 2003 USA
MH121111 HAdV-C2 2017 Germany
MH121088 HAdV-C2 2013 Germany
JX173077.1 HAdV-C2 2005 Argentina
MH121086 HAdV-C2 2013 Germany
KF268310.1 HAdV-C2 1992 USA
MH121084 HAdV-C2 2012 Germany
MH121083 HAdV-C2 2012 Germany
MH121090 HAdV-C2 2014 Germany
MH121077 HAdV-C2 2009 Germany
MH121101 HAdV-C2 2015 Germany
MH121102 HAdV-C2 2015 Germany
MF315028 HAdV-C2 2013 China
HQ413315.1 HAdV-C6 USA
FJ349096.1 HAdV-C6 1953 USA
MH121099 HAdV-C2 2015 Germany
MH121079 HAdV-C2 2010 Germany
MH121092 HAdV-C2 2014 Germany
KR699642.1 HAdV-CBJ113 2009 China
MH121078 HAdV-C2 2009 Germany
MF315029 HAdV-C2 2013 China
MH121095 HAdV-C2 2014 Germany
JX173085.1 HAdV-C1 2003 USA
MH121087 HAdV-C1 2013 Germany
MH121082 HAdV-C1 2012 Germany
JX173086.1 HAdV-C1 2004 USA
GOSH.2110 HAdV-C1 2016 England
JX173083.1 HAdV-C1 2003 USA
GOSH.2068 HAdV-C1 2016 England
GOSH.2049 HAdV-C1 2015 England
GOSH.2050 HAdV-C1 2015 England
MH121116 HAdV-C1 2017 Germany
MH121100 HAdV-C1 2015 Germany
JX173078.1 HAdV-C1 2000 Argentina
MH121117 HAdV-C1 2017 Germany
MH121089 HAdV-C1 2013 Germany
MH121108 HAdV-C1 2016 Germany
JX173082.1 HAdV-C1 2003 USA
AC_000017.1 HAdV-C1
MH121110 HAdV-C1 2017 Germany
JX173081.1 HAdV-C2 2002 Egypt
JX173080.1 HAdV-C1 2001 Egypt
MH121081 HAdV-C1 2012 Germany
MH121073 HAdV-C1 2002 Germany
MH121074 HAdV-C1 2002 Germany
MH121076 HAdV-C1 2008 Germany
MH121075 HAdV-C1 2008 Germany
MH121098 HAdV-C1 2015 Germany
JX173079.1 HAdV-C2 2002 Argentina
MH121097 HAdV-C2 2015 Germany
KX384959.1 HAdV-C2 2003 USA
MH121114 HAdV-C2 2017 Germany
GOSH.2063 HAdV-C2 2015 England
MH121112 HAdV-C6 2017 Germany
LC068715.1 HAdV-C6 1994 Japan
JX423389.1 HAdV-C6 2008 USA
LC068712.1 HAdV-C6 1988 Japan
LC068714.1 HAdV-C6 1993 Japan
LC068718.1 HAdV-C6 2005 Japan
LC068716.1 HAdV-C6 2004 Japan
LC068720.1 HAdV-C6 2005 Japan
LC068719.1 HAdV-C6 2005 Japan
LC068717.1 HAdV-C6 2004 Japan
KF951595.1 HAdV-DD28 2013 China
LC068713.1 HAdV-C6 1988 Japan
HQ003817.1 HAdV-C-P1H57F6 2002 Azerbaijan
Yana2 HAdV-C1 31.600BP Siberia
AY339865.1 HAdV-C5
AY601635.1 HAdV-C5
KF429754.1 HAdV-C5 1990 USA
KF268127.1 HAdV-C5 1988 USA
AC_000008.1 HAdV-C5
KF268199.1 HAdV-C5 2008 USA
GOSH.2045 HAdV-C5 2016 England
GOSH.2046 HAdV-C5 2016 England
MH121094 HAdV-C5 2014 Germany
GOSH.2007 HAdV-C5 2015 England
MF681662.1 HAdV-C5
MH121113 HAdV-C6 2017 Germany
KF268129.1 HAdV-C6 2005 USA
Yana1 HAdV-C2 31.600BP Siberia
FJ025900.1 SAdV-43
0.02
HAdV-C5
HAdV-C1
SAdV-43
HAdV-C6
HAdV-C2
HAdV-CP1H57F6
Yana1
Yana2

L5

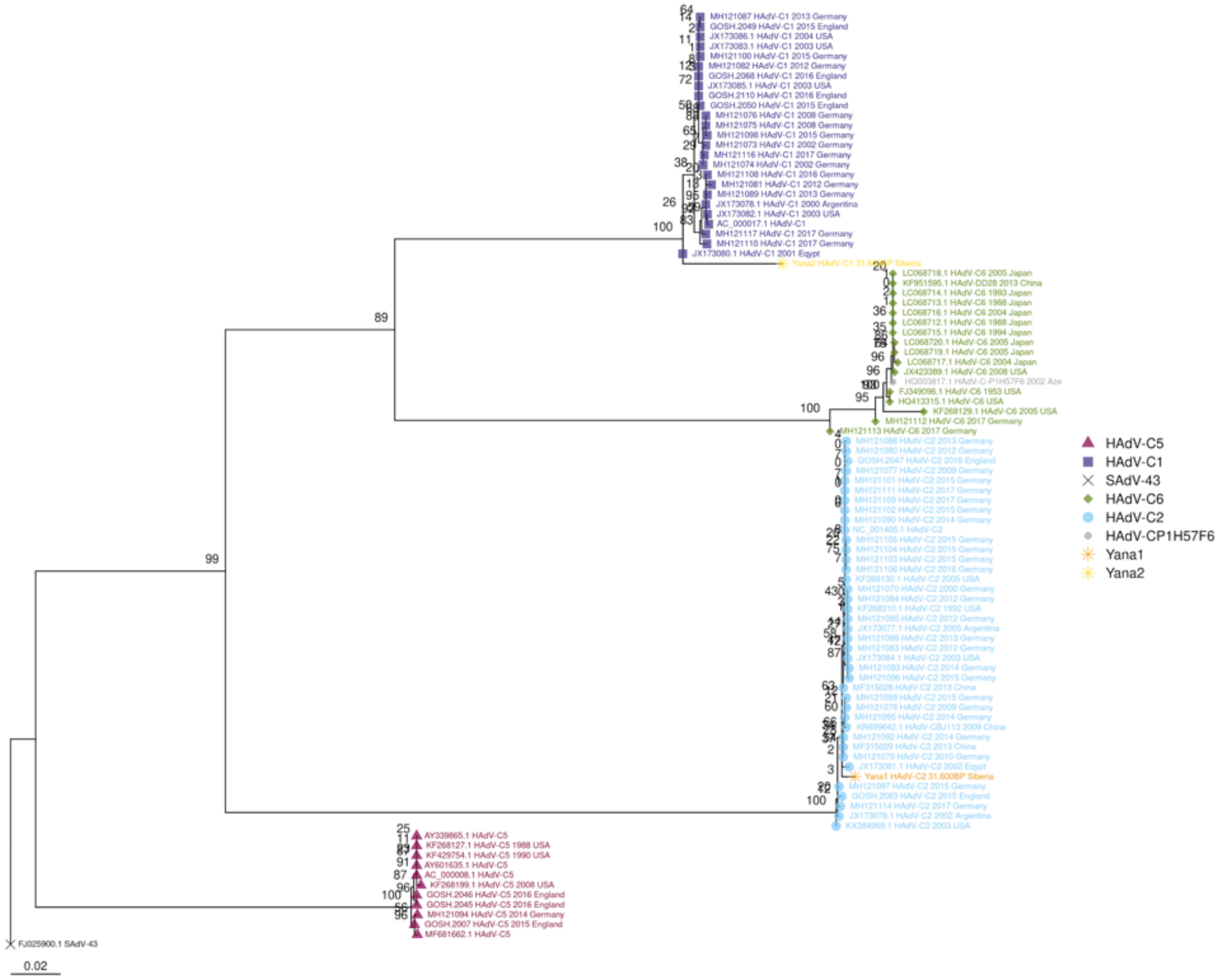

HAdV-C5
HAdV-C1
SAdV-43
HAdV-C6
HAdV-C2
HAdV-CP1H57F6
Yana1
Yana2
0.02

pTP

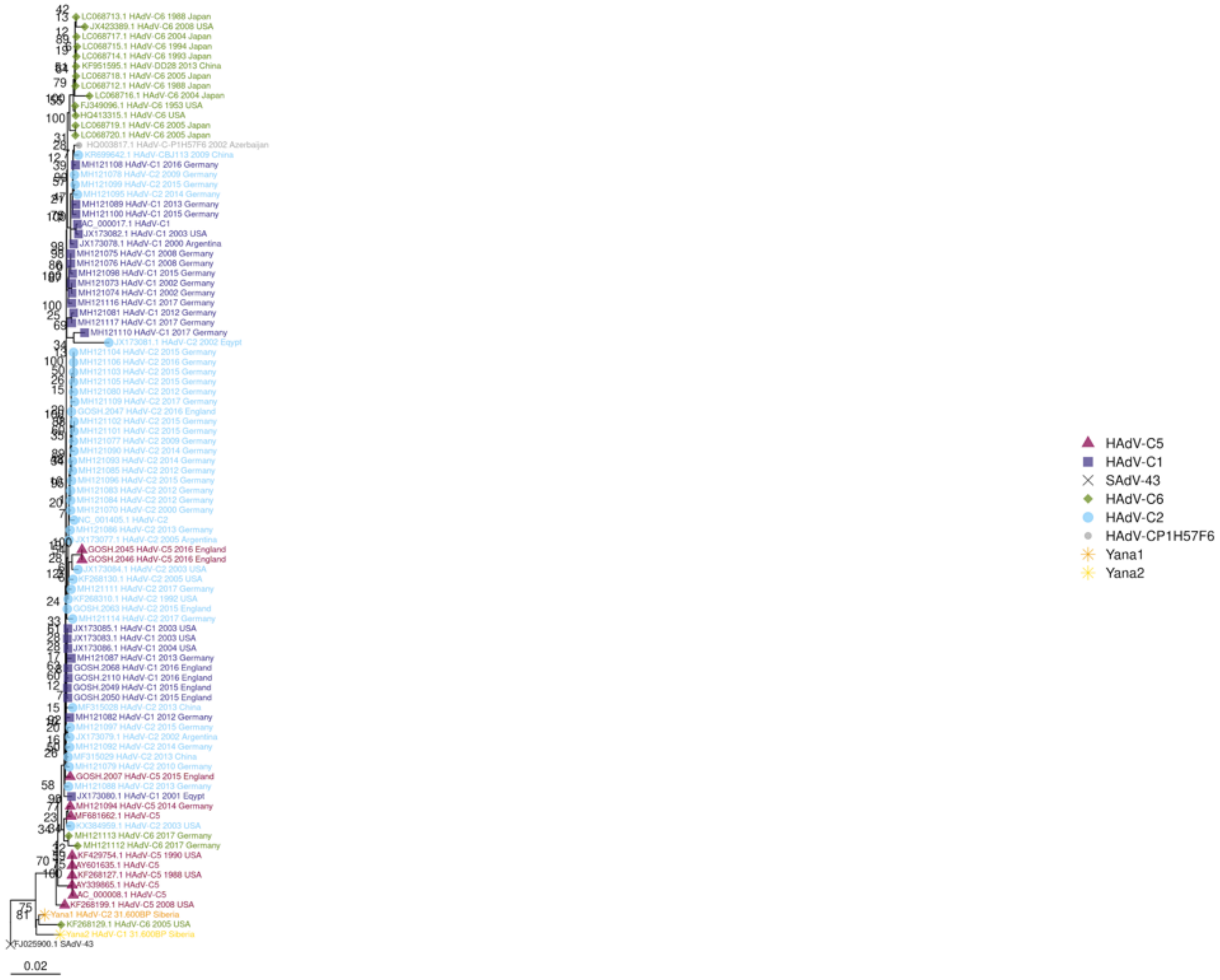
HAdV-C5
HAdV-C1
SAdV-43
HAdV-C6
HAdV-C2
HAdV-CP1H57F6
Yana1
Yana2
0.02

U

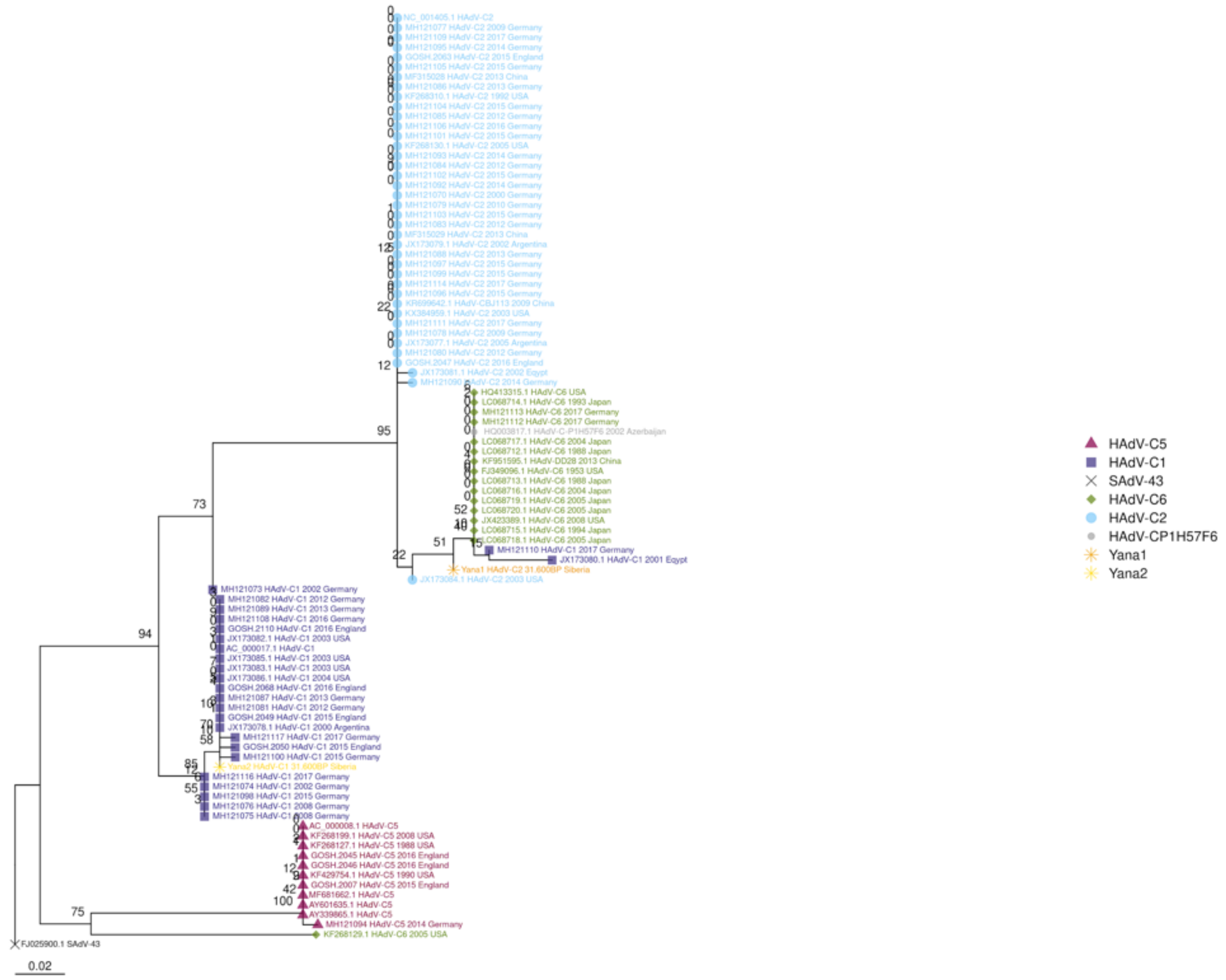

**Fig. S28. Maximum likelihood gene trees**, model GTR+G-I, outgroup branch removed for visualization.

E1A_6_kDa_protein

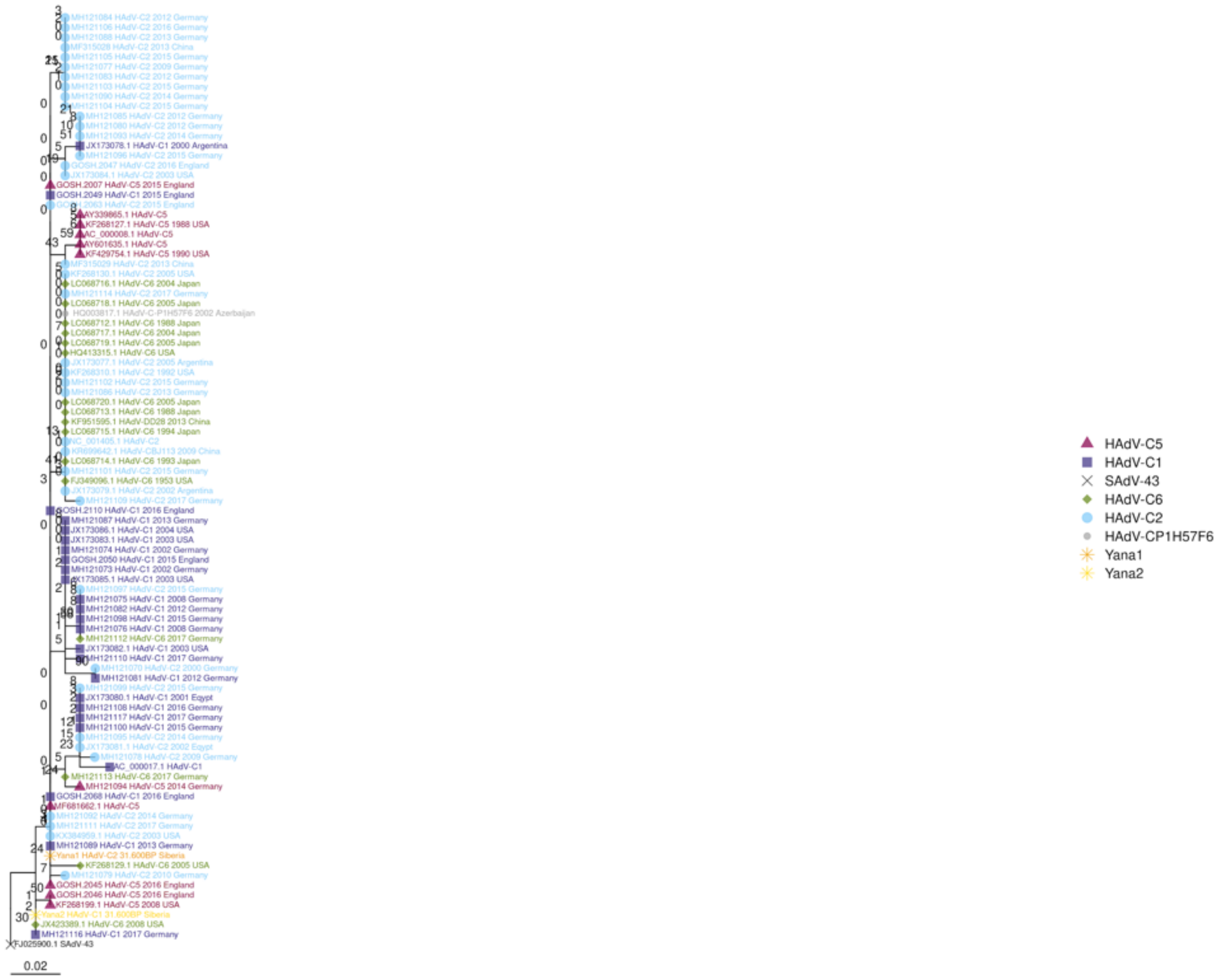

E1A_26_kDa_protein_control_protein_E1A_243R

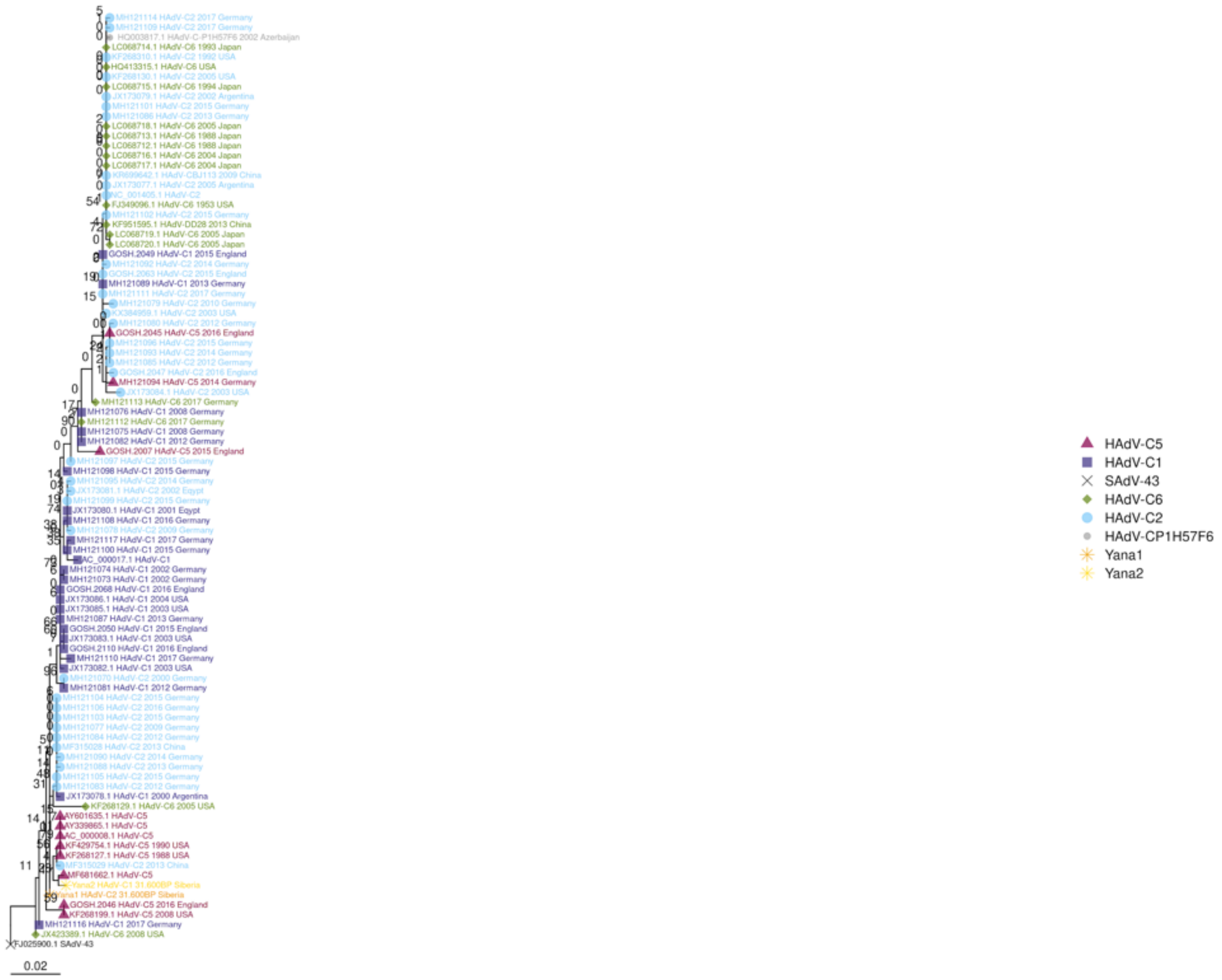

E1A_32_kDa_protein_control_protein_E1A

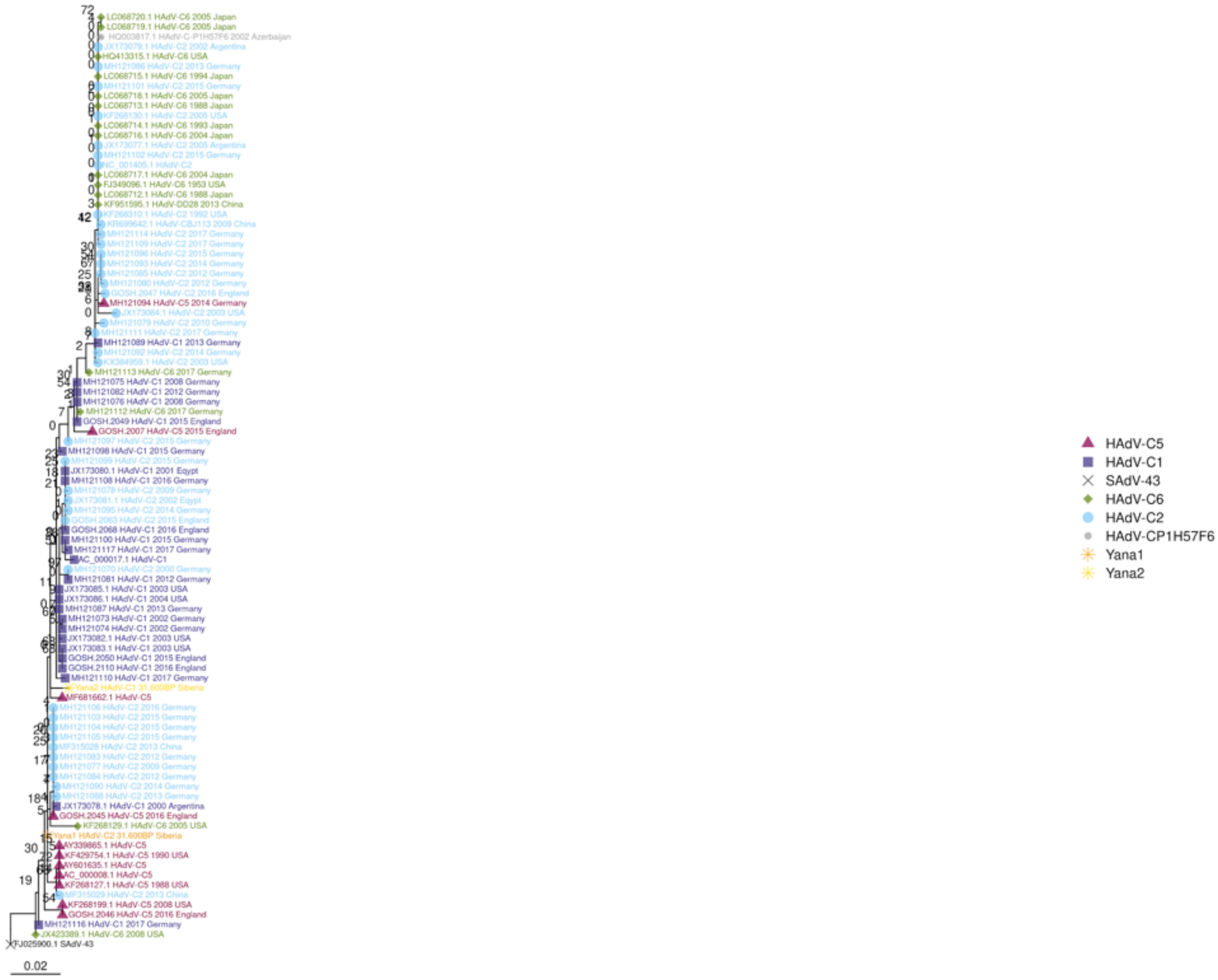

E1B_large_T_antigen

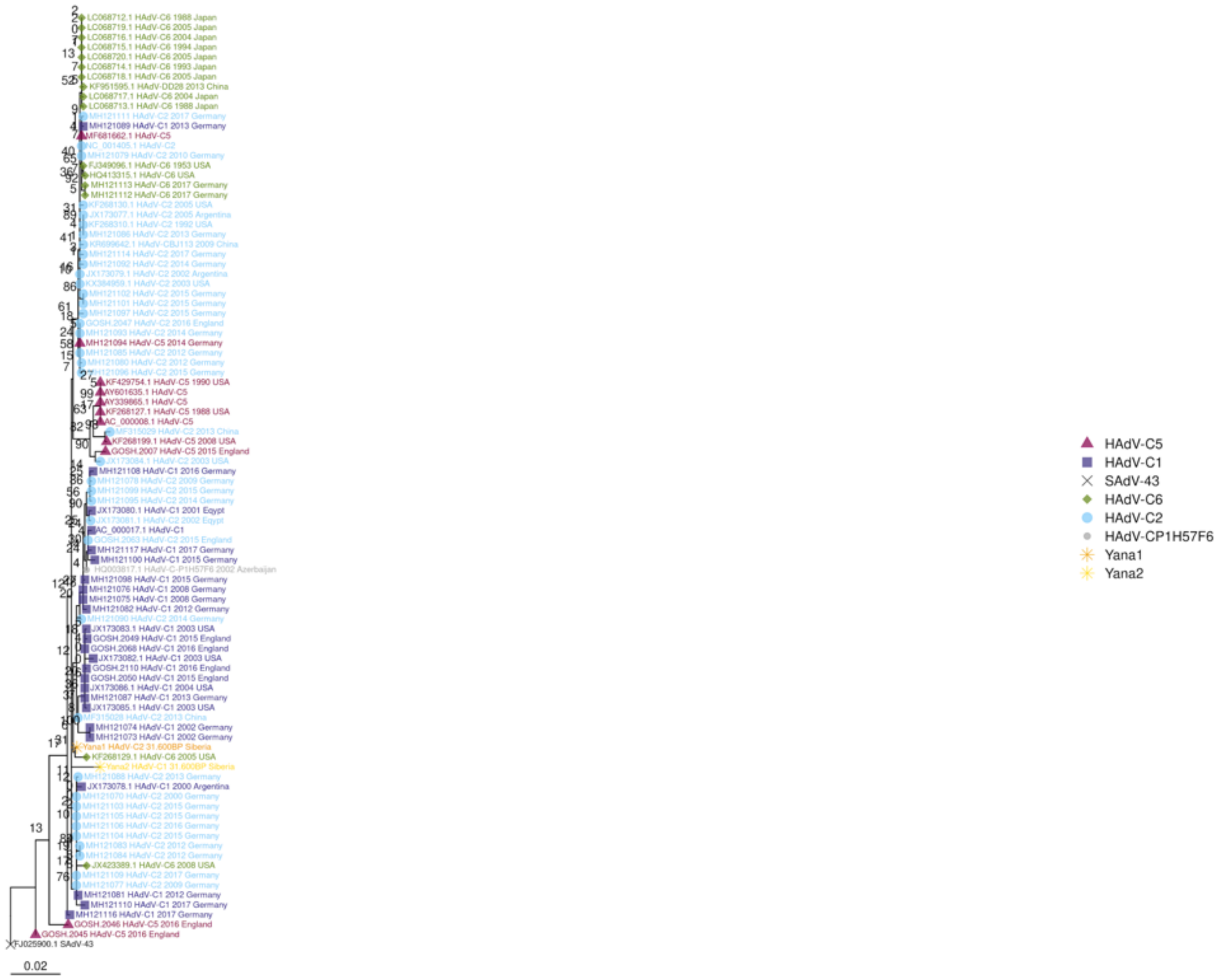

E1B,_E1B_small_T_antigen

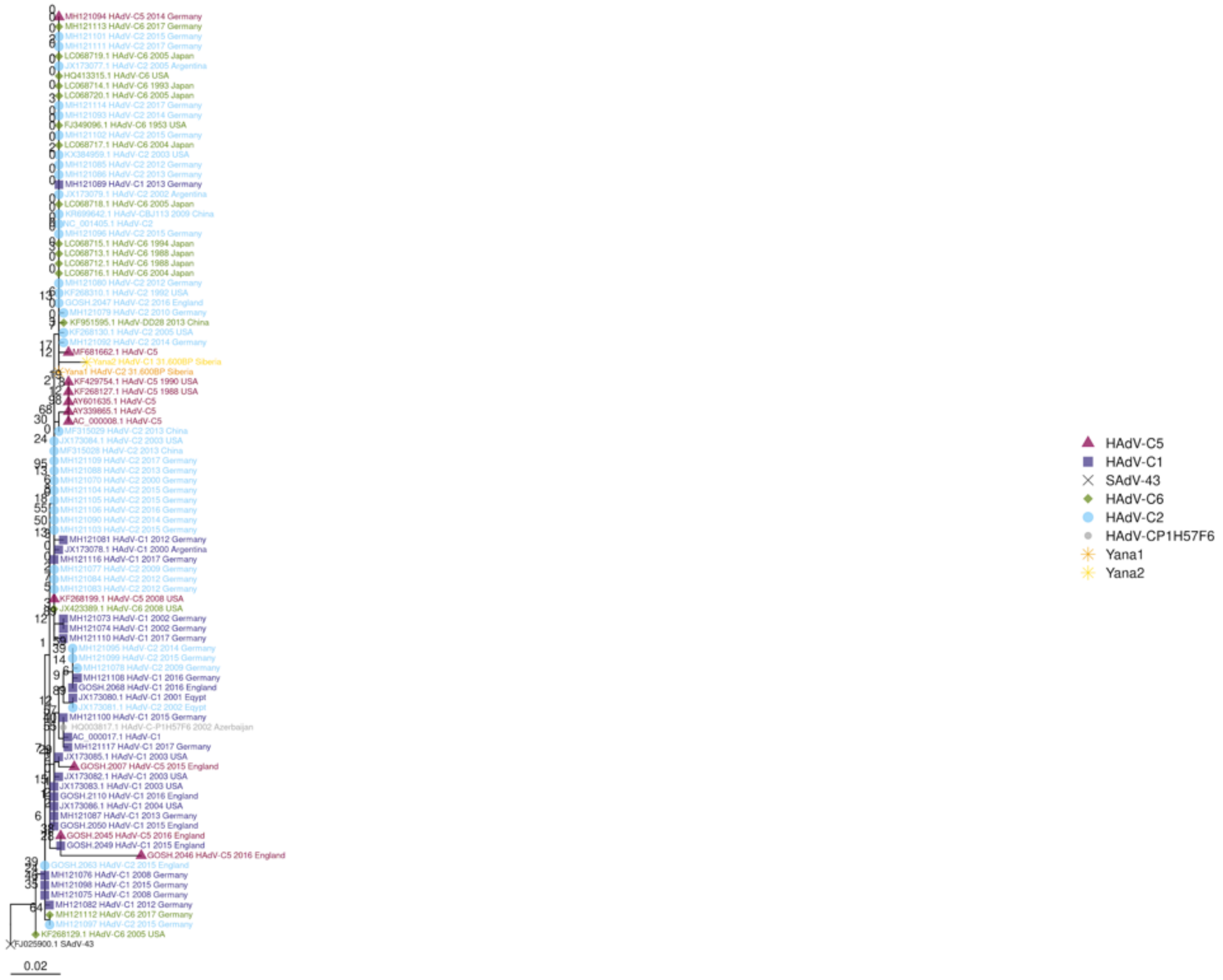

E2A_ssDNA-binding_protein

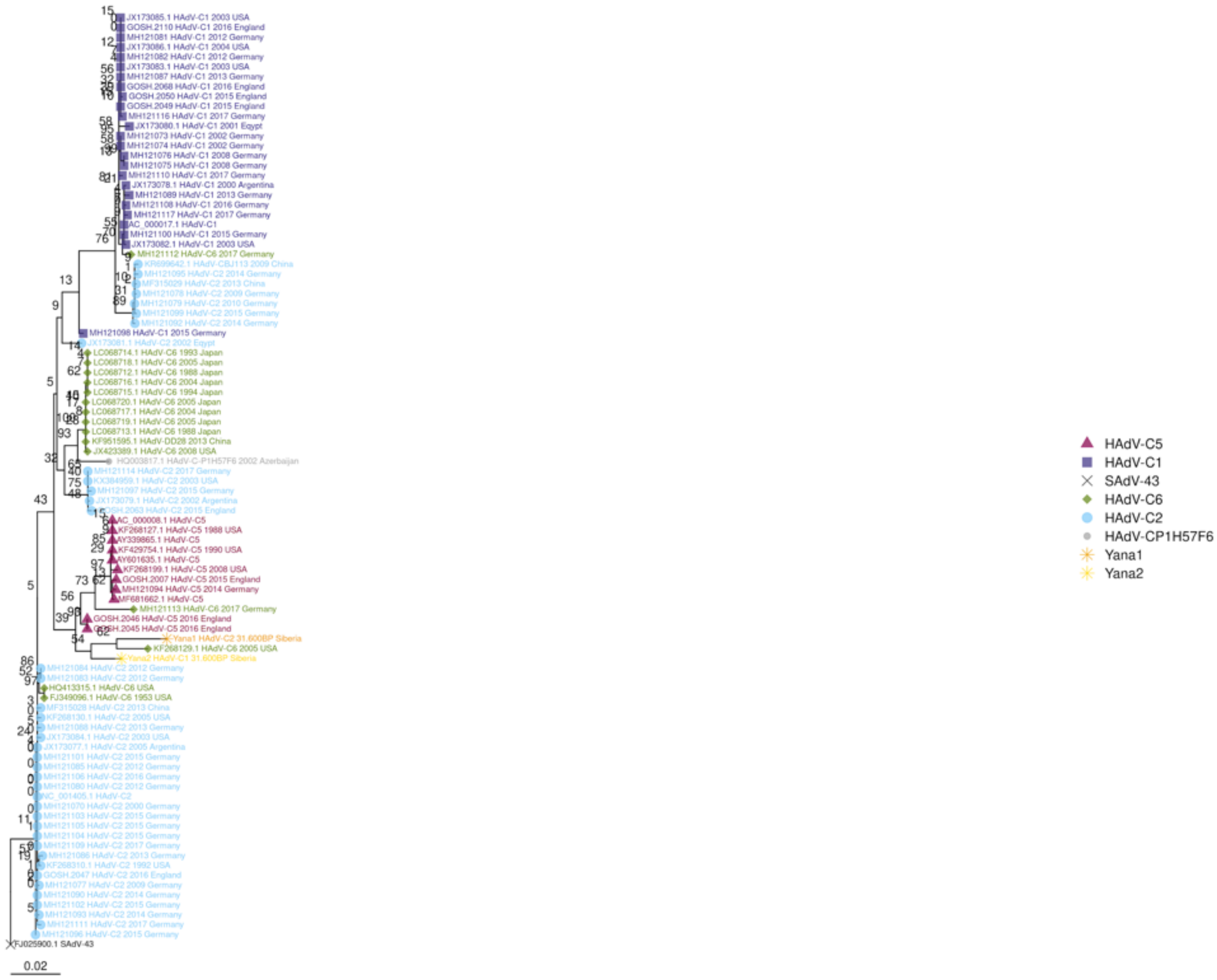

E2B_DNA_polymerase_complement

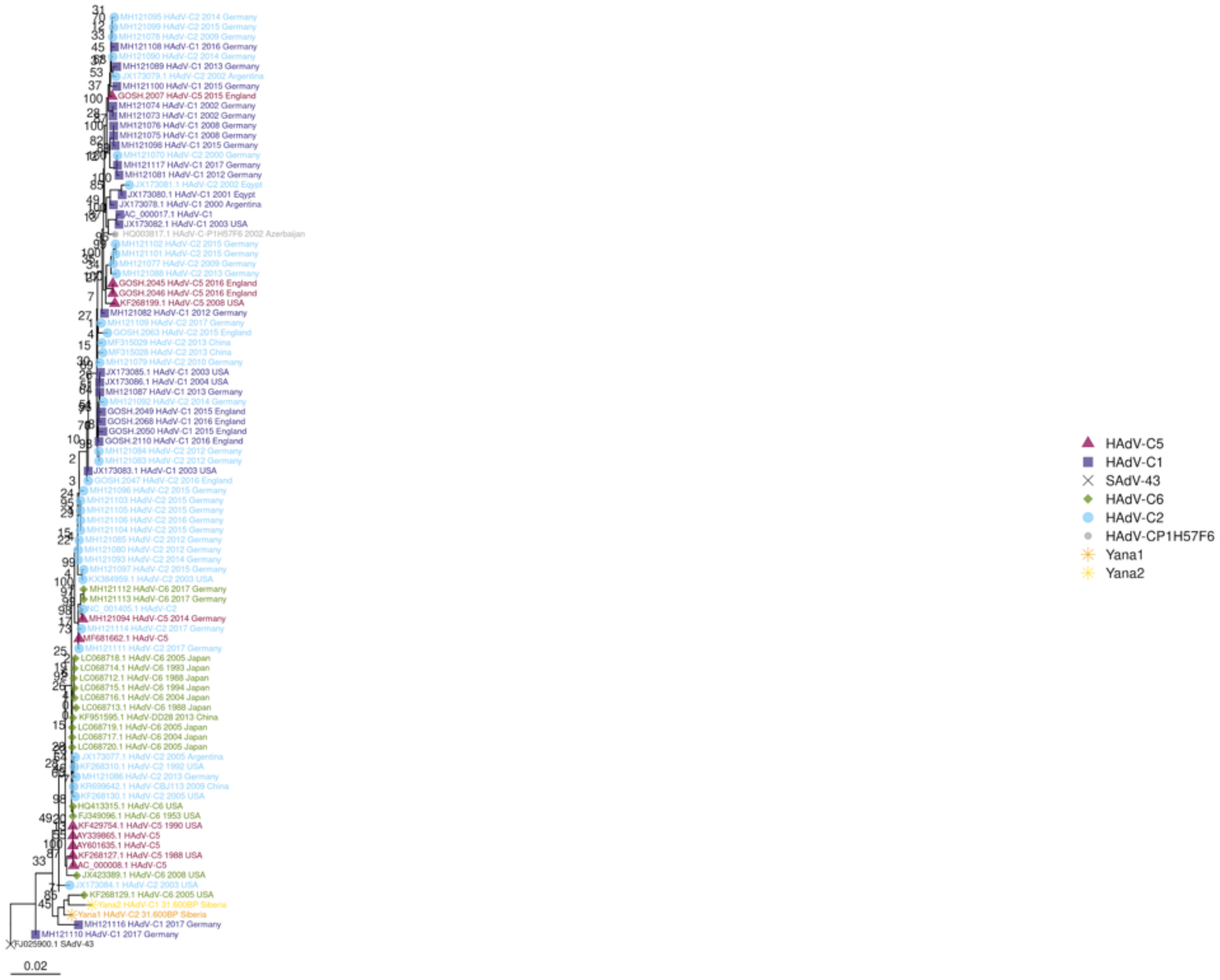

E3_12

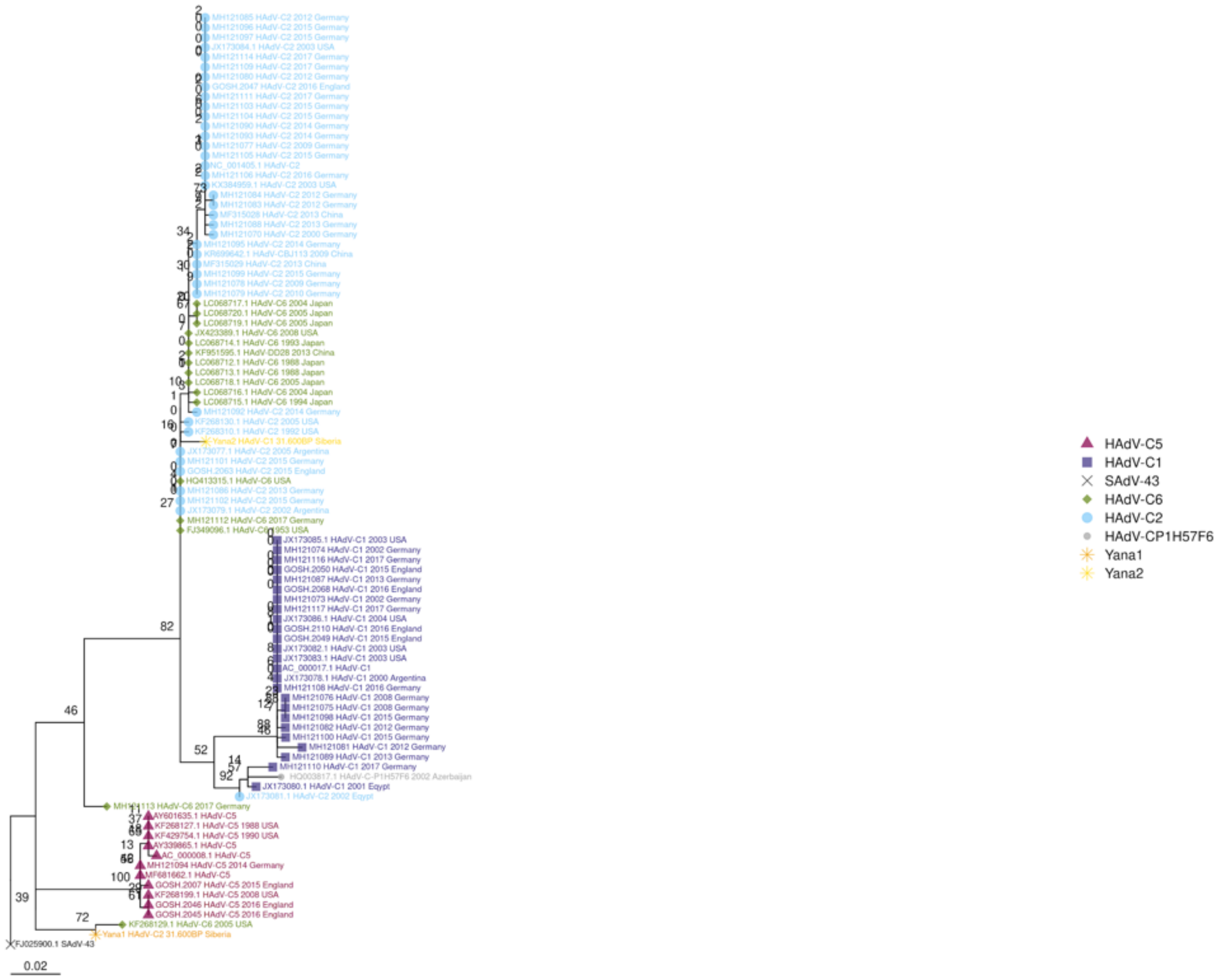
HAdV-C5
HAdV-C1
SAdV-43
HAdV-C6
HAdV-C2
HAdV-CP1H57F6
Yana1
Yana2
0.02

E3_control_protein_14

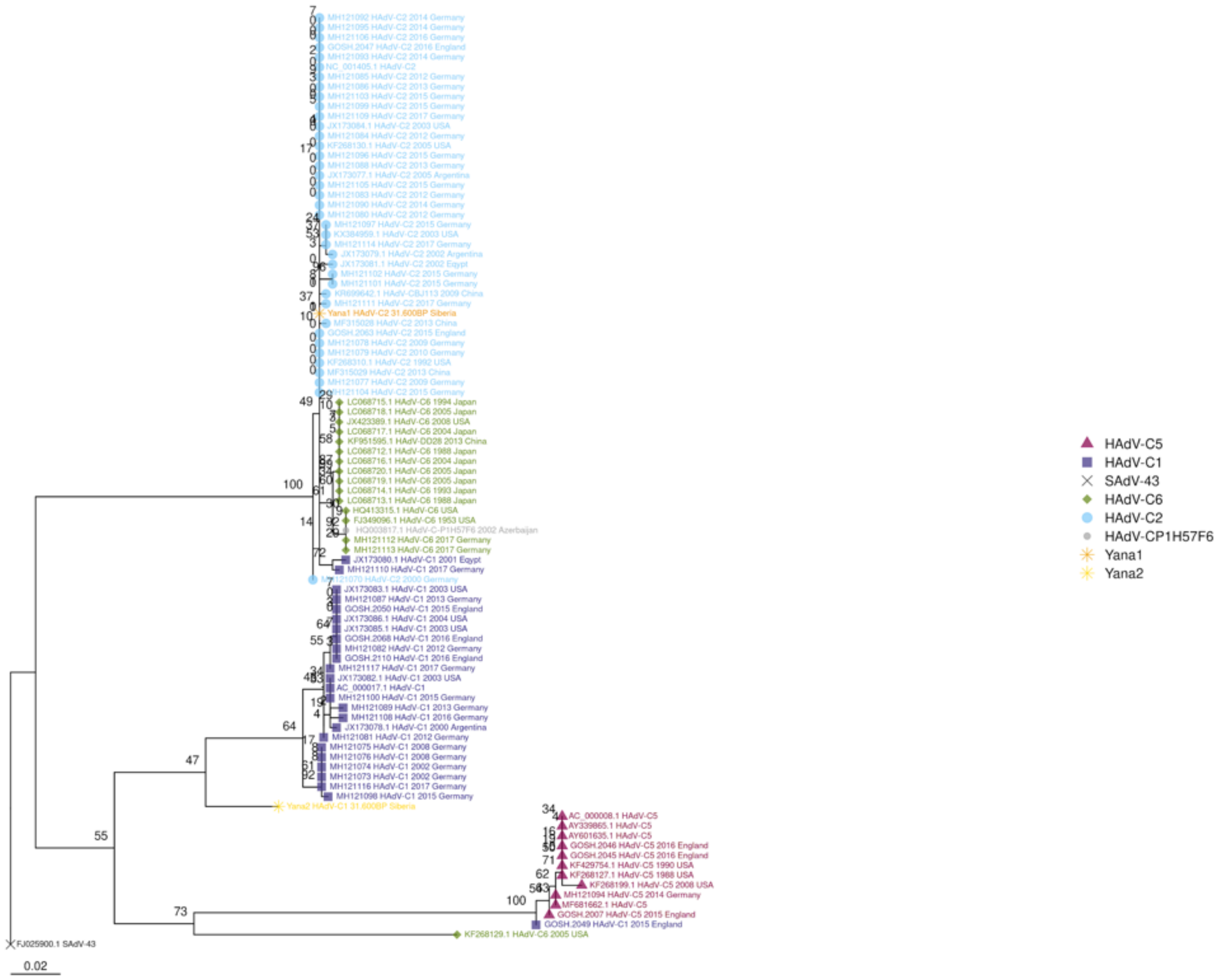

E3_CR1-alpha

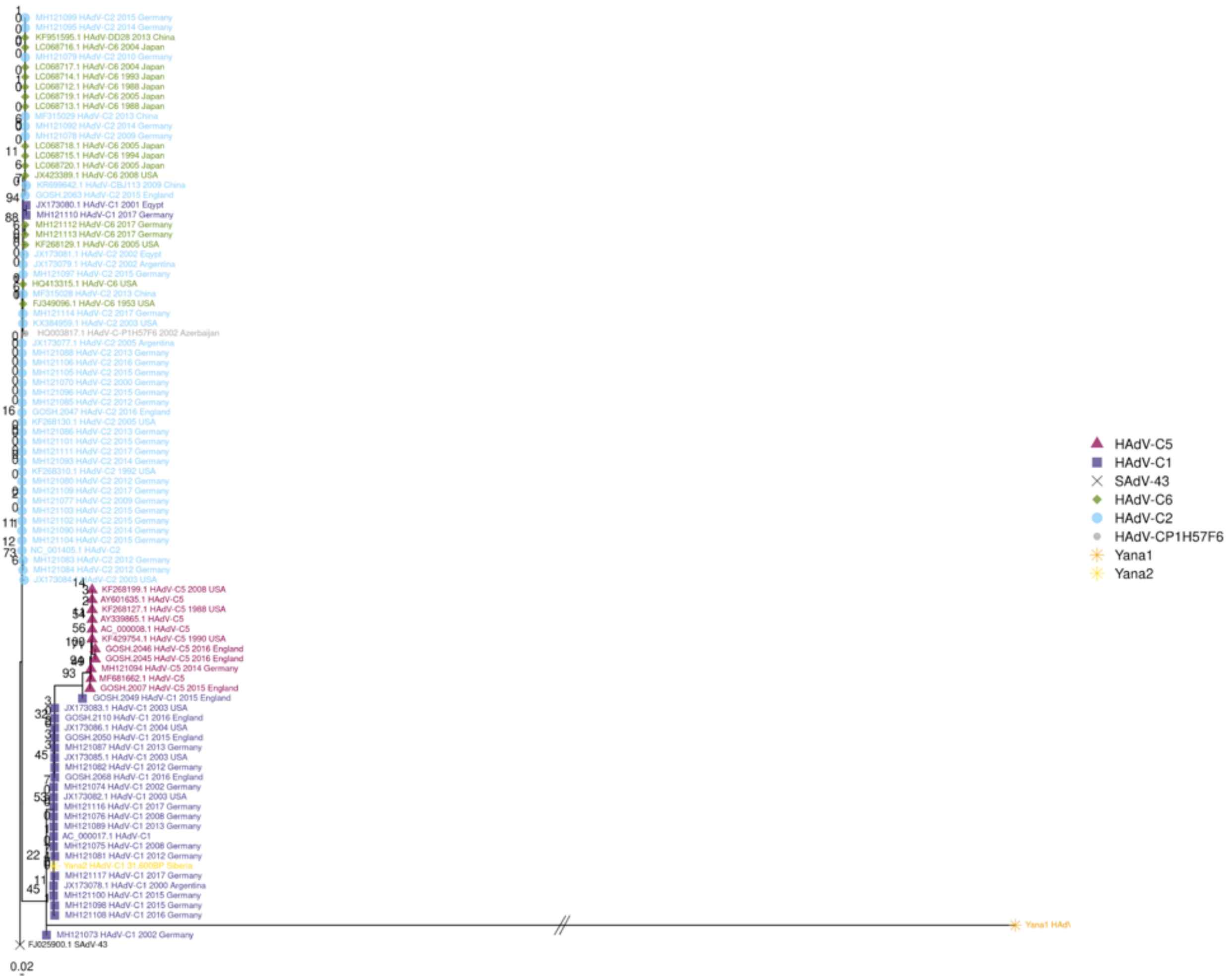
HAdV-C5
HAdV-C1
SAdV-43
HAdV-C6
HAdV-C2
HAdV-CP1H57F6
Yana1
Yana2
0.02

E3_CR1-beta

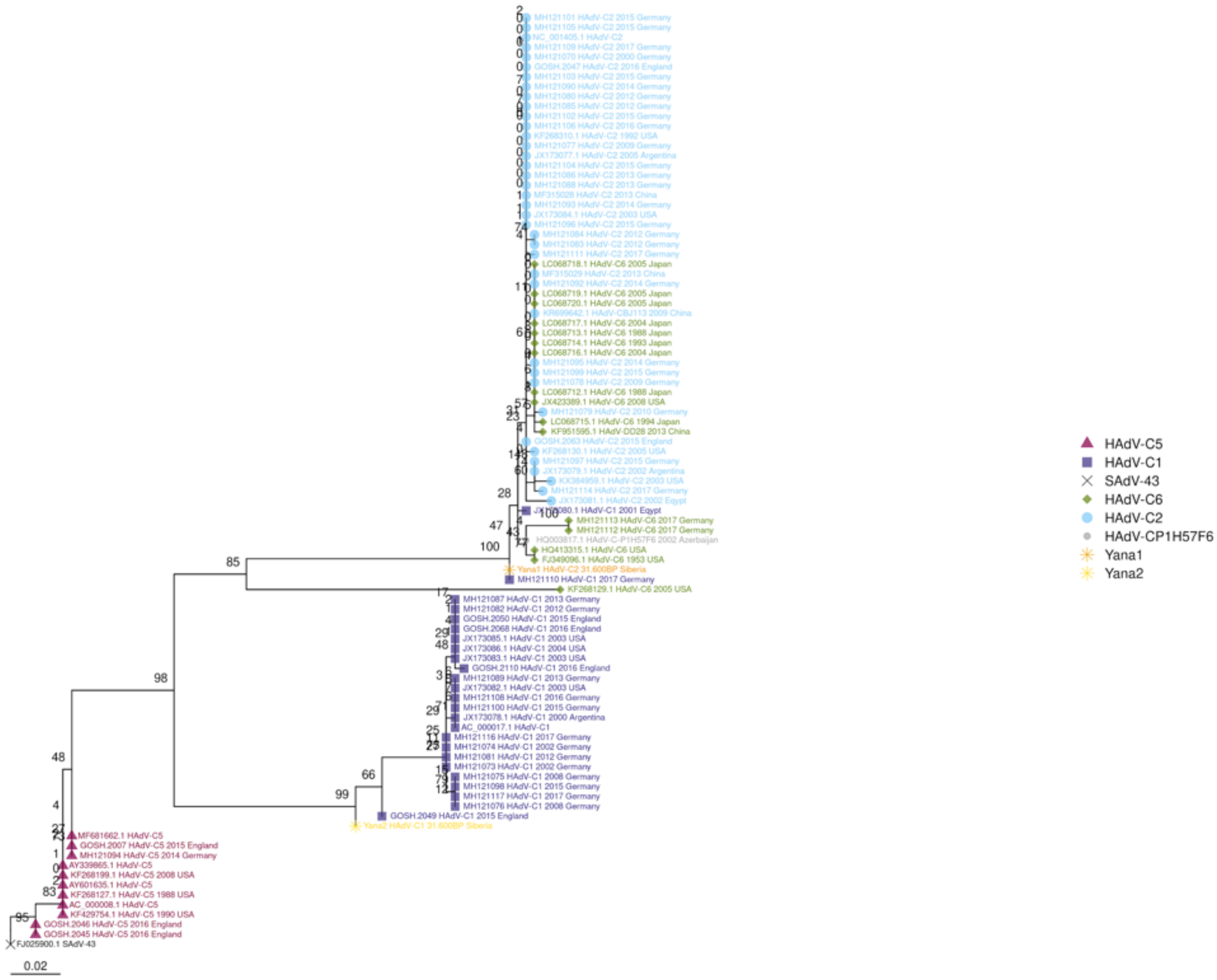
HAdV-C5
HAdV-C1
SAdV-43
HAdV-C6
HAdV-C2
HAdV-CP1H57F6
Yana1
Yana2
0.02

E3_immune_modulating_protein_18

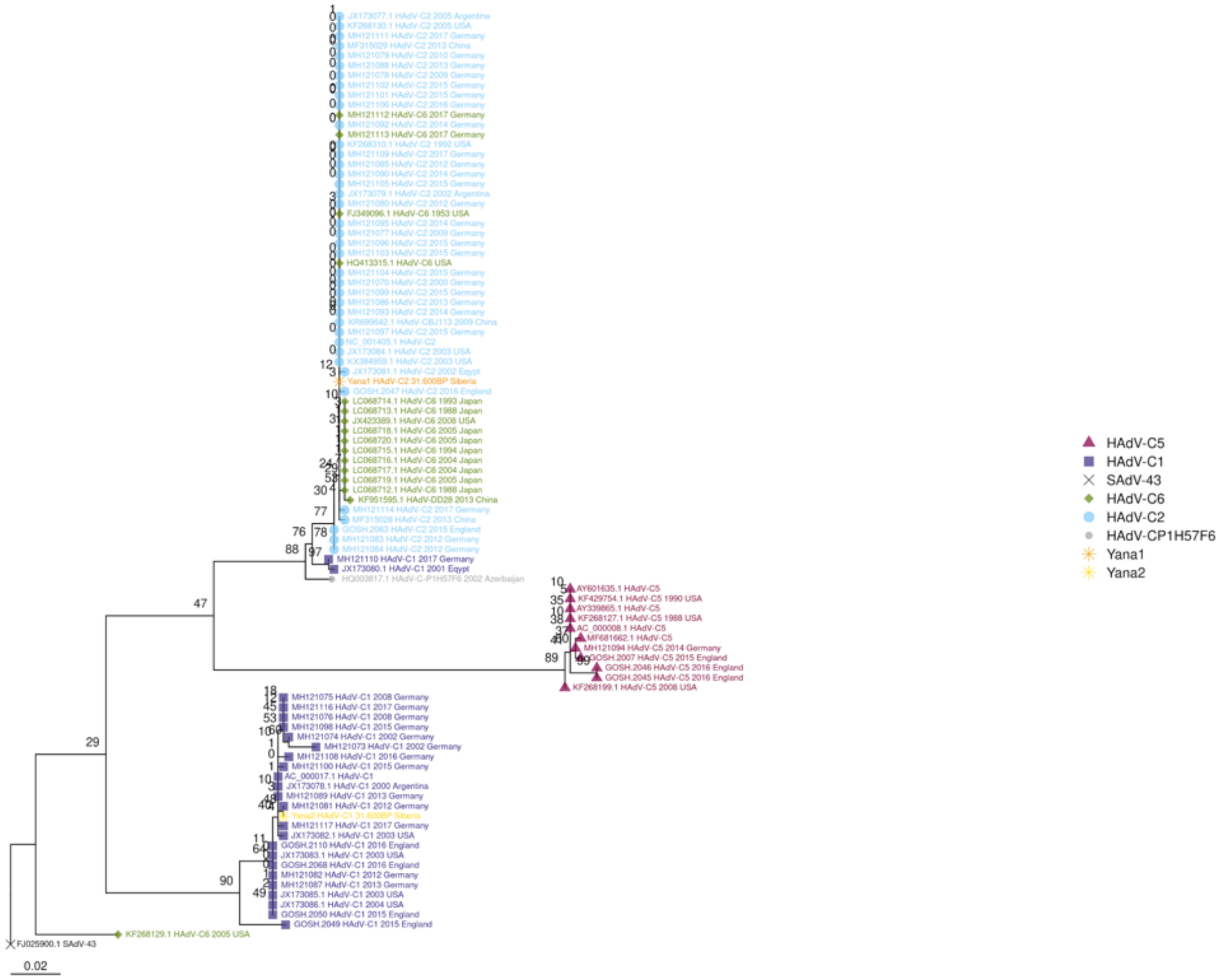

E3_RID-alpha

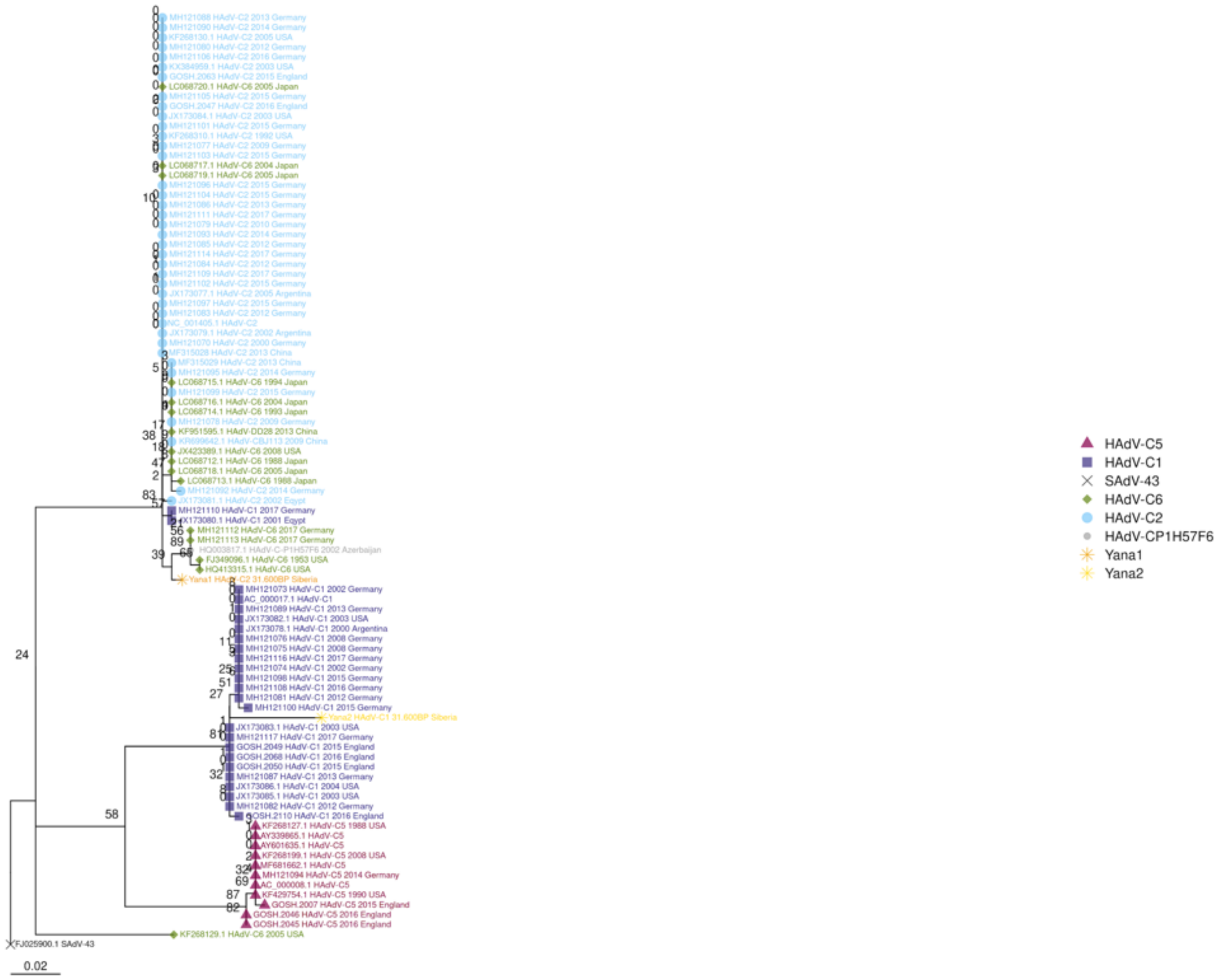

HAdV-C5
HAdV-C1
SAdV-43
HAdV-C6
HAdV-C2
HAdV-CP1H57F6
Yana1
Yana2
0.02

E3_RID-beta

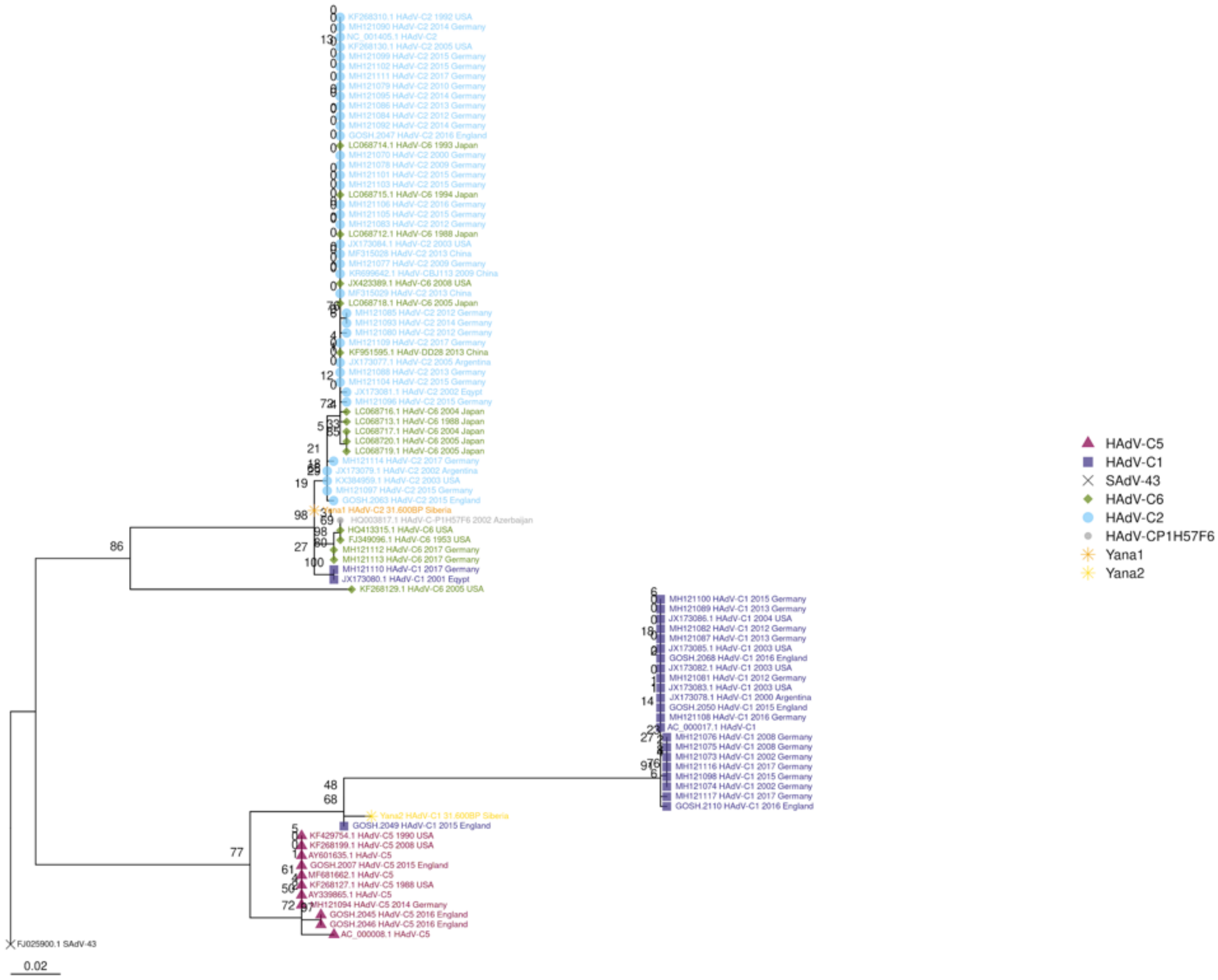
HAdV-C5
HAdV-C1
SAdV-43
HAdV-C6
HAdV-C2
HAdV-CP1H57F6
Yana1
Yana2
0.02

E3_CR1-alpha

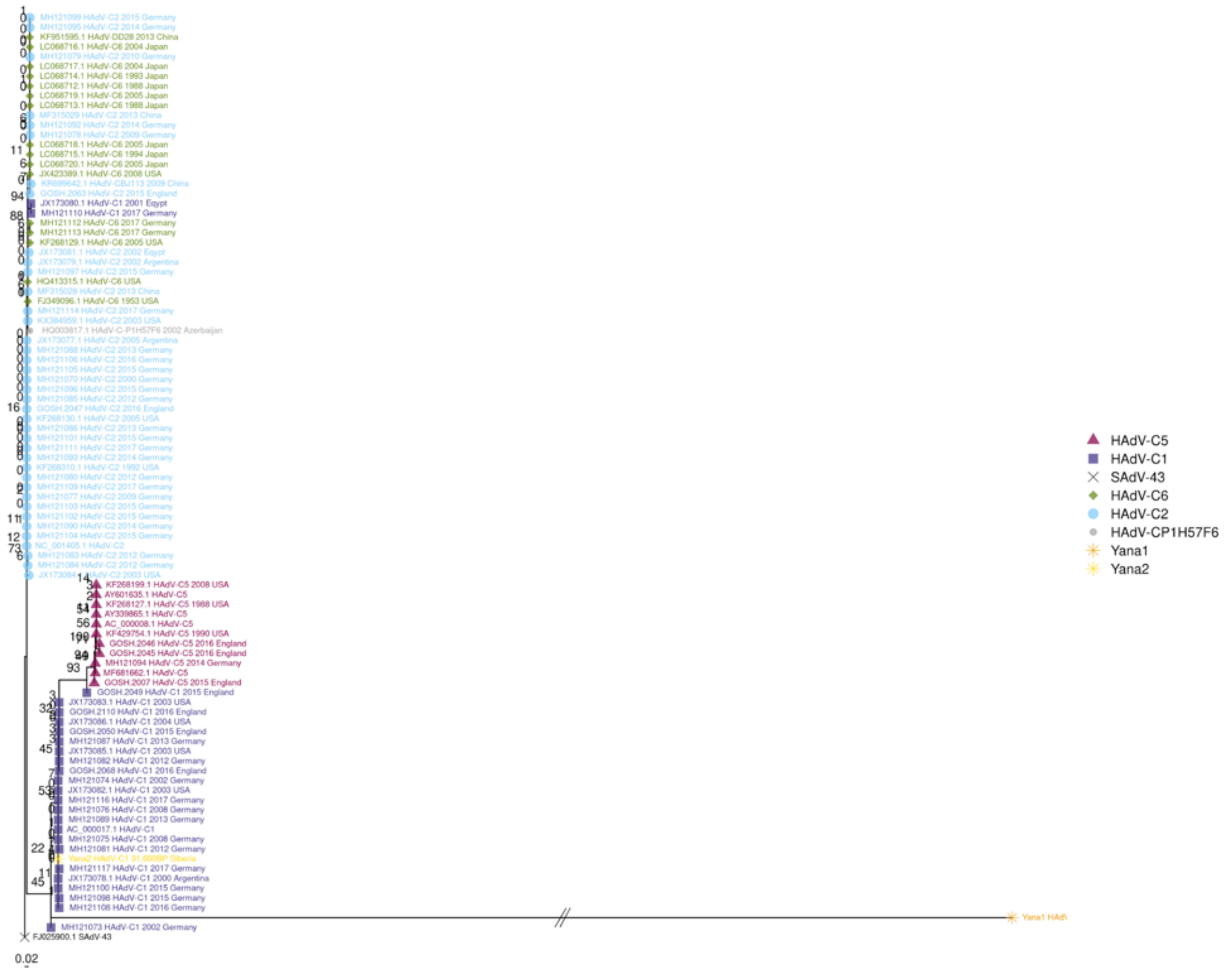
HAdV-C5
HAdV-C1
SAdV-43
HAdV-C6
HAdV-C2
HAdV-CP1H57F6
Yana1
Yana2
0.02

E4_control_protein_orf1

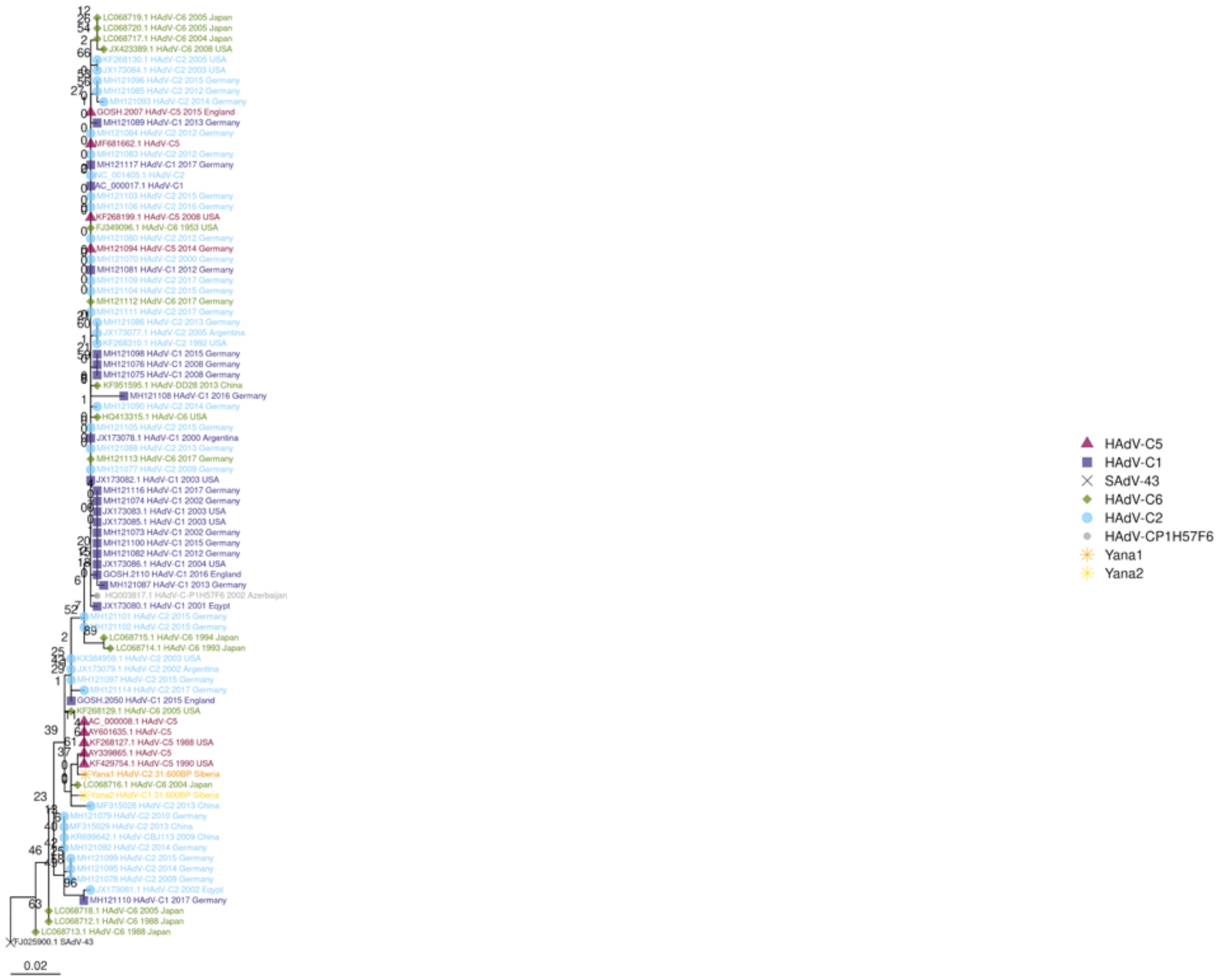

E4_control_protein_orf2

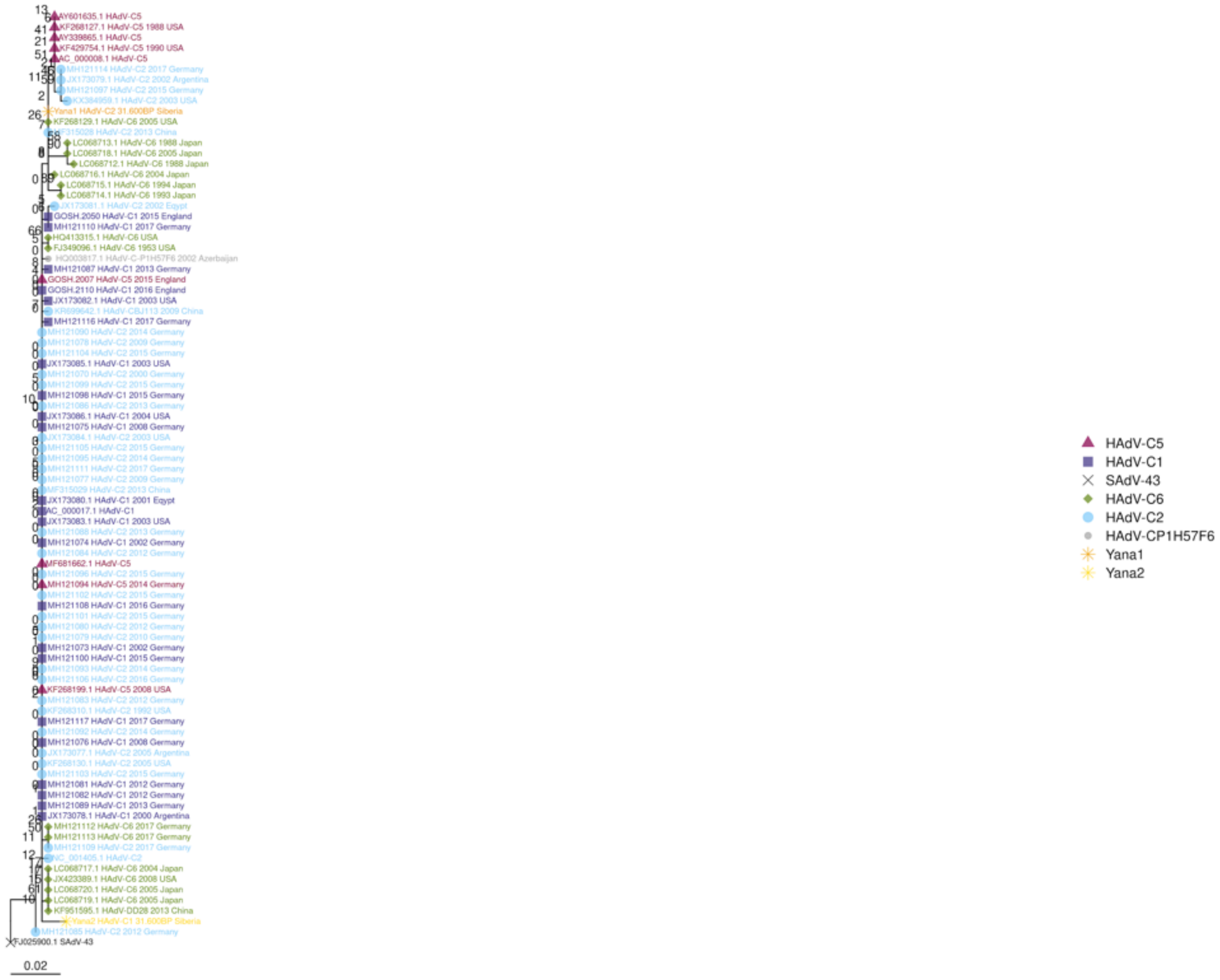

E4_control_protein_orf3_4

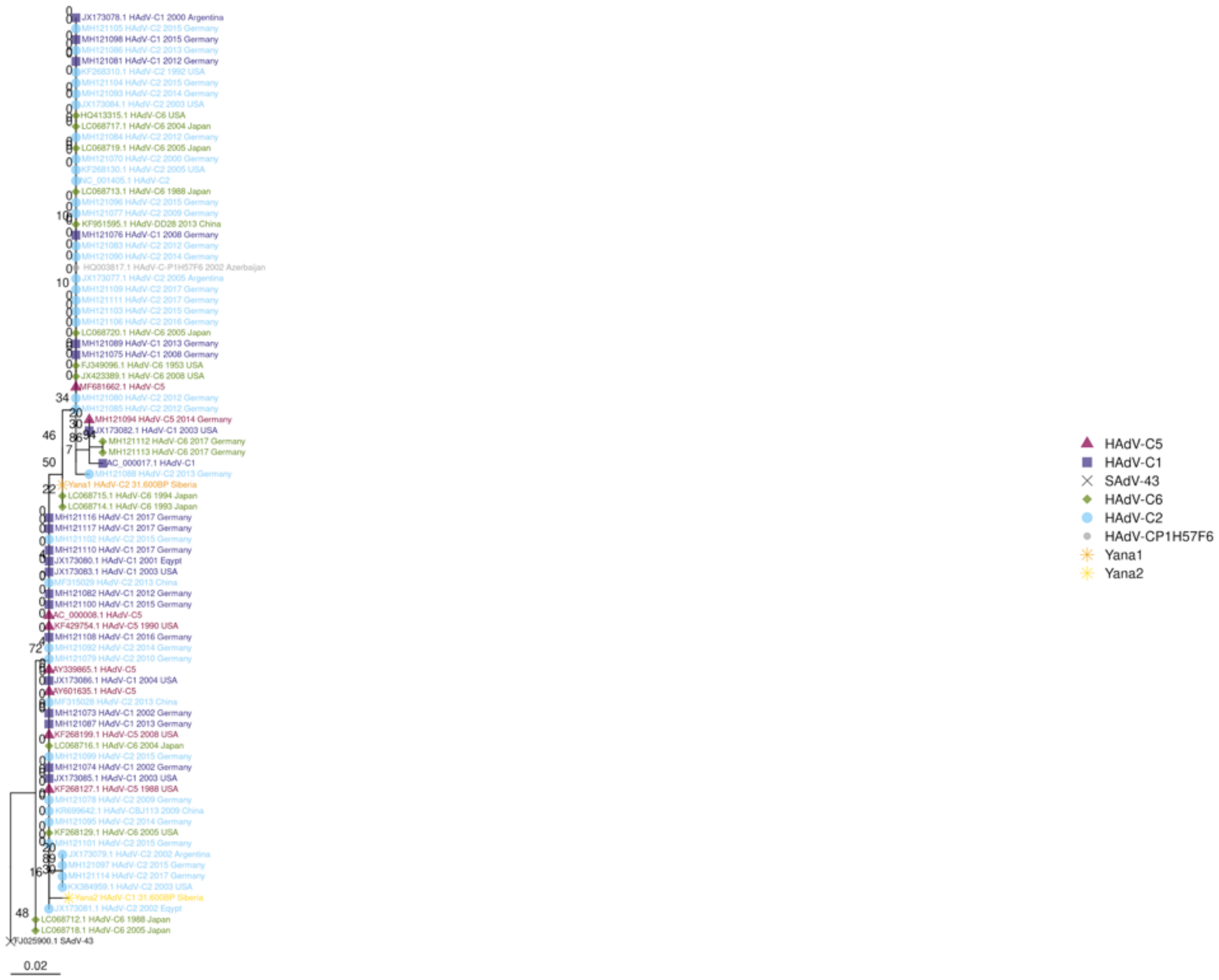

E4_control_protein_orf3

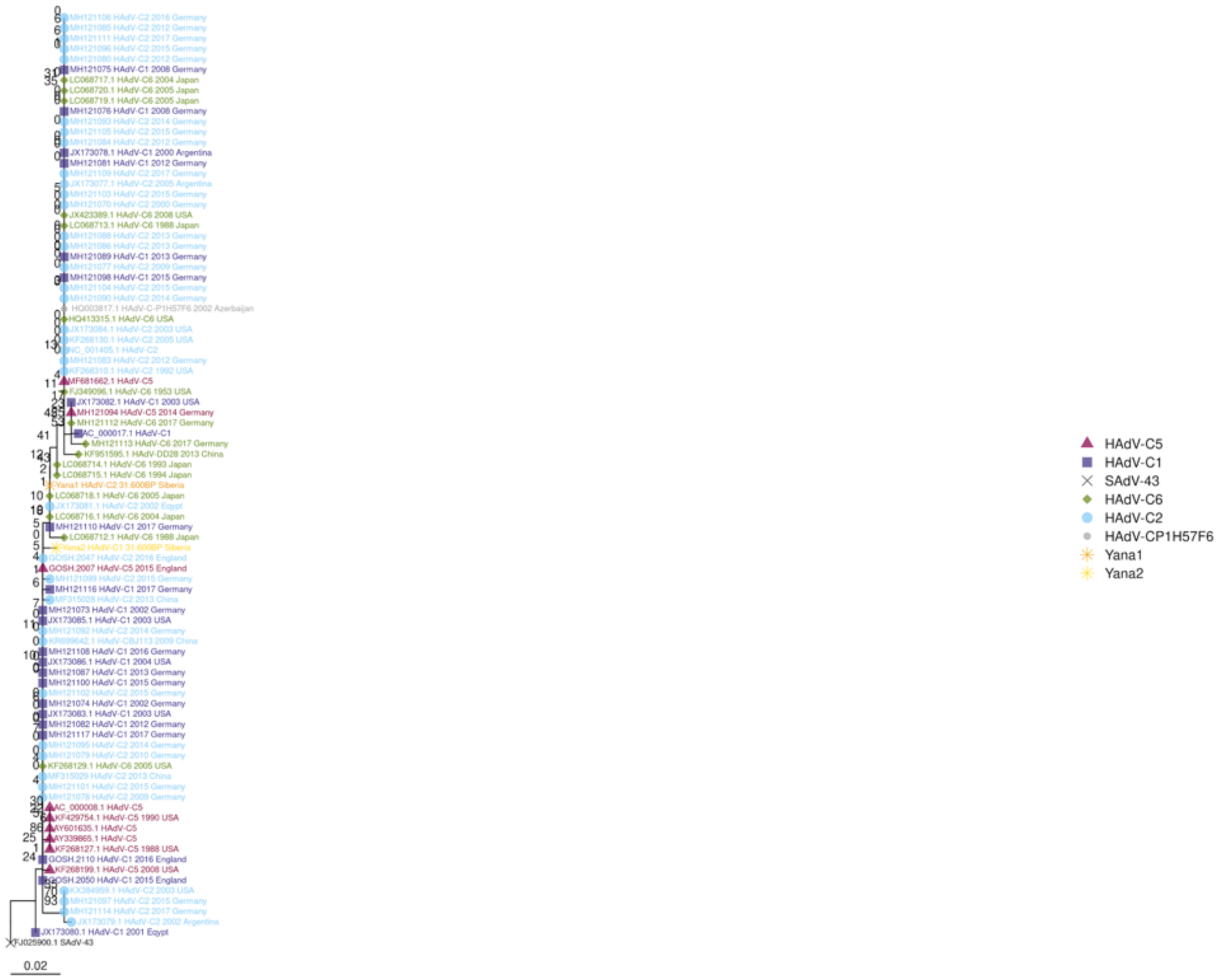

E4_control_protein_orf4

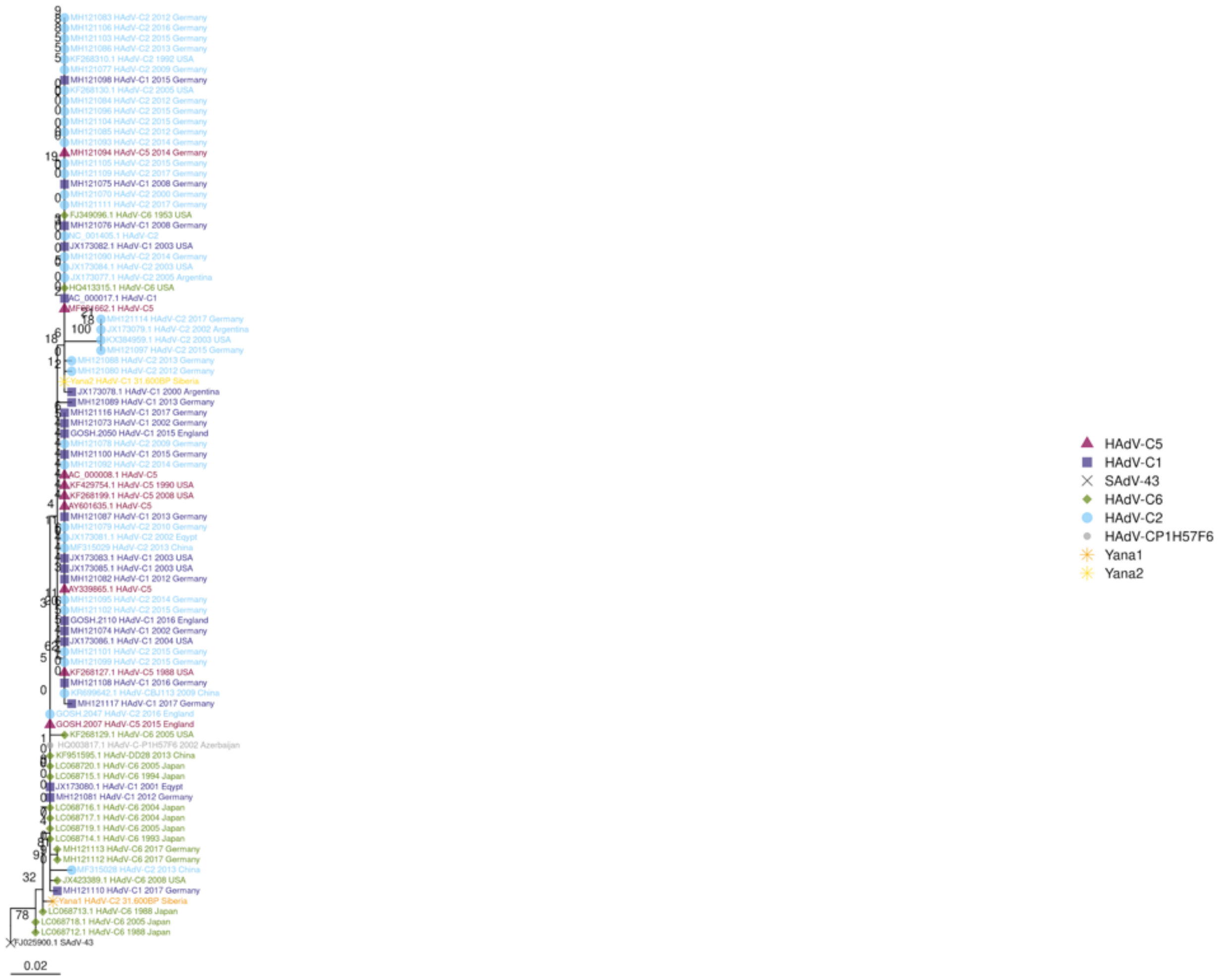

E4_control_protein_orf6_7

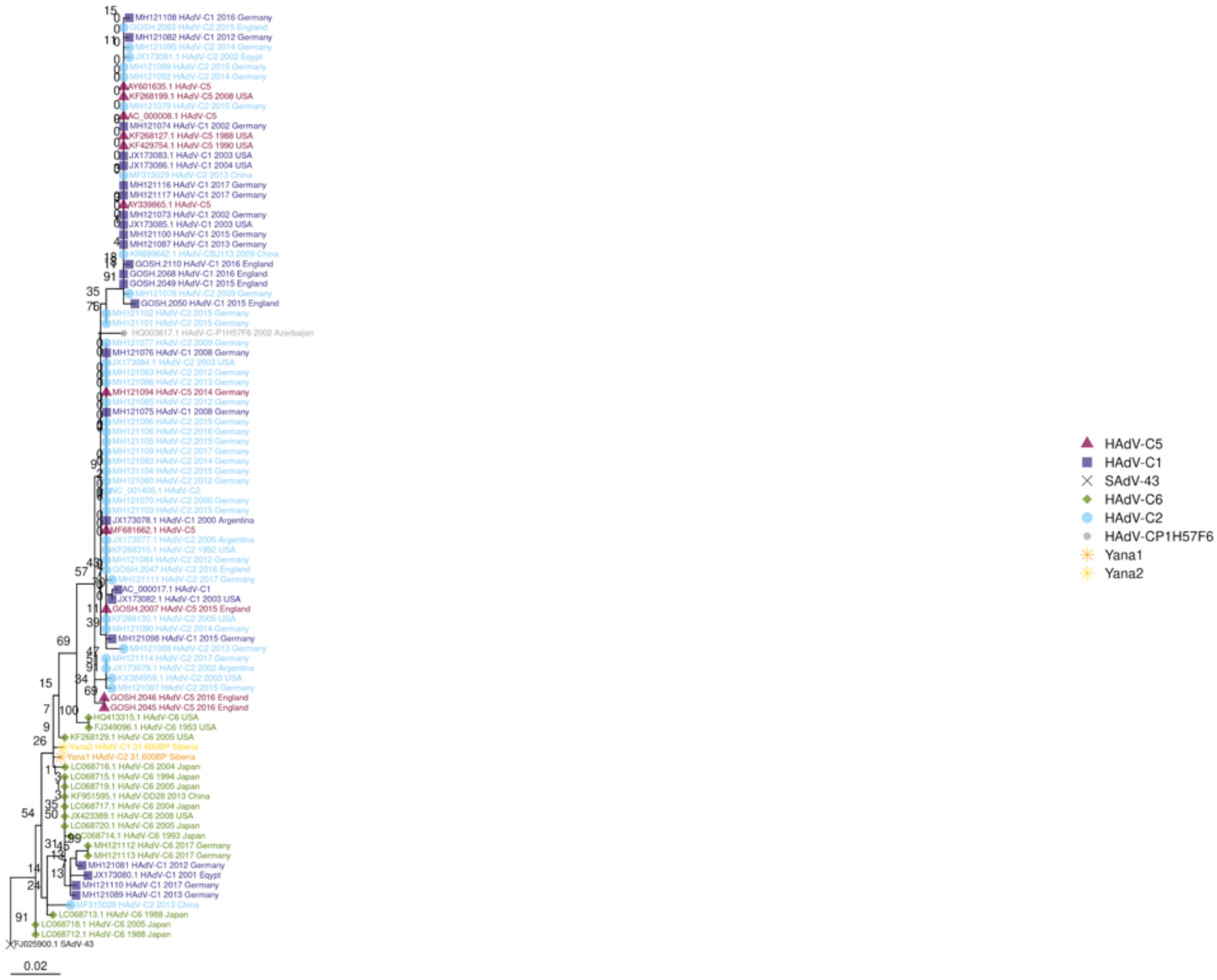

IVa2_maturation_protein

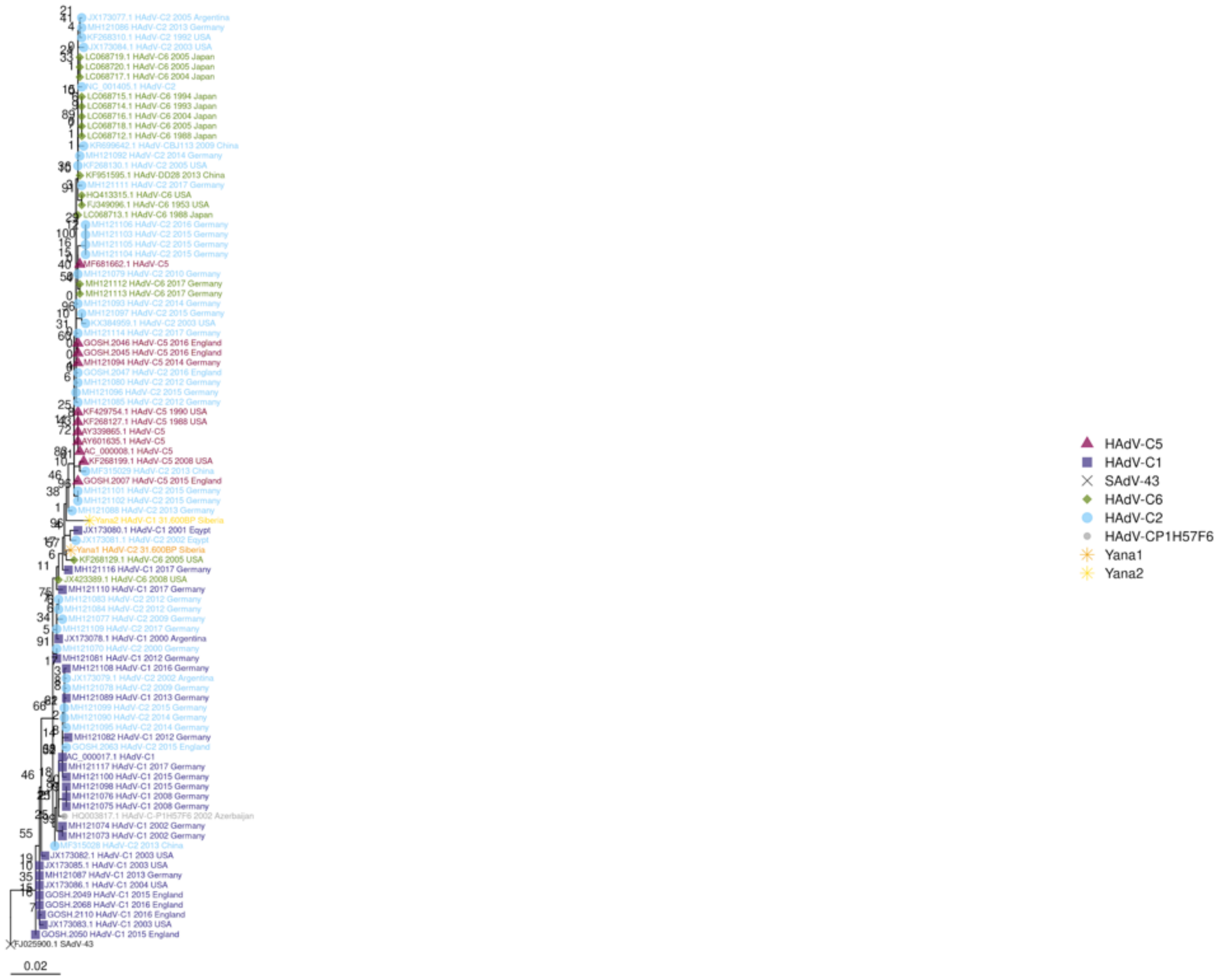

IX_hexon_associated_protein

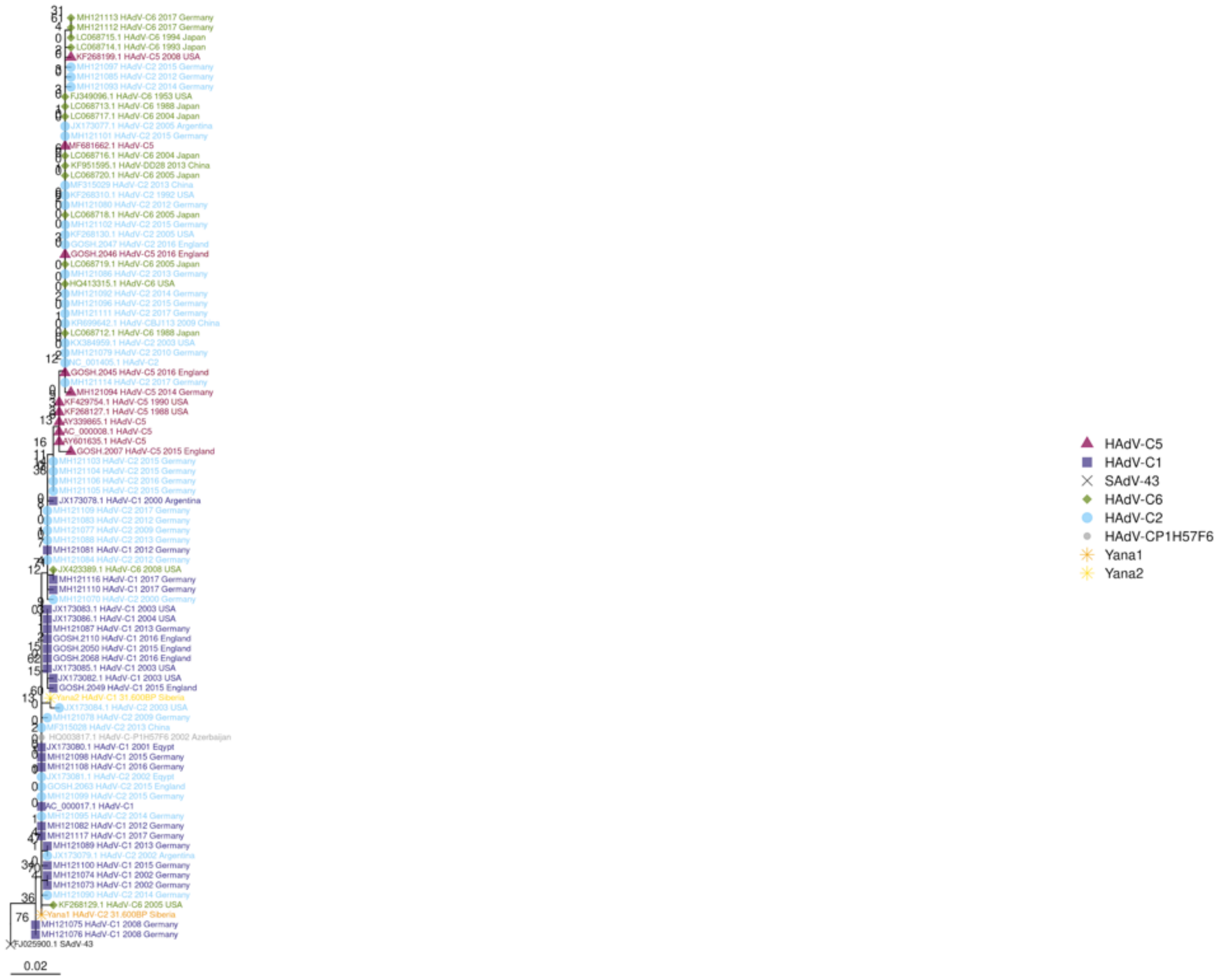

L1_13

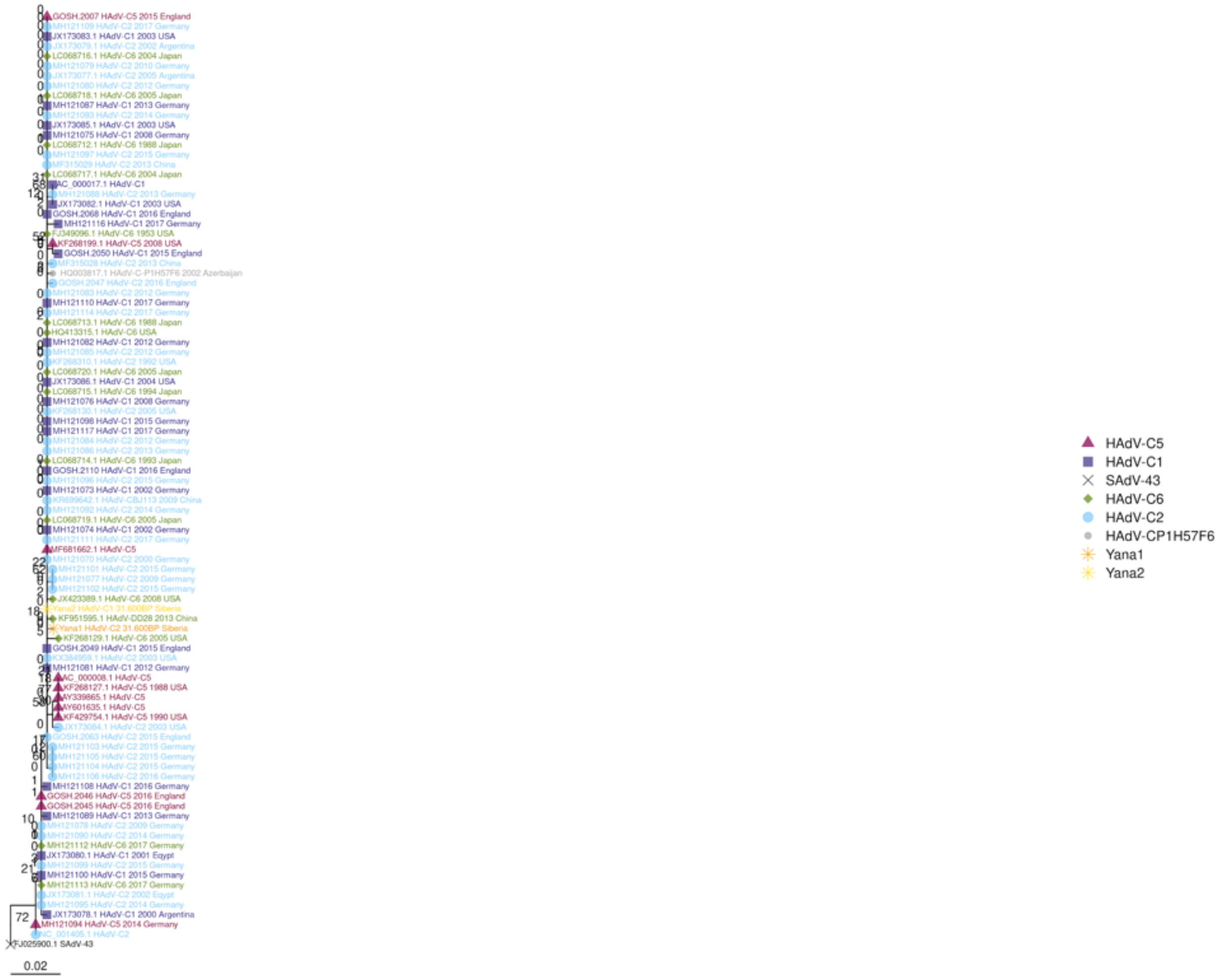

HAdV-C5
HAdV-C1
SAdV-43
HAdV-C6
HAdV-C2
HAdV-CP1H57F6
Yana1
Yana2
0.02

L1_52_kDa_protein

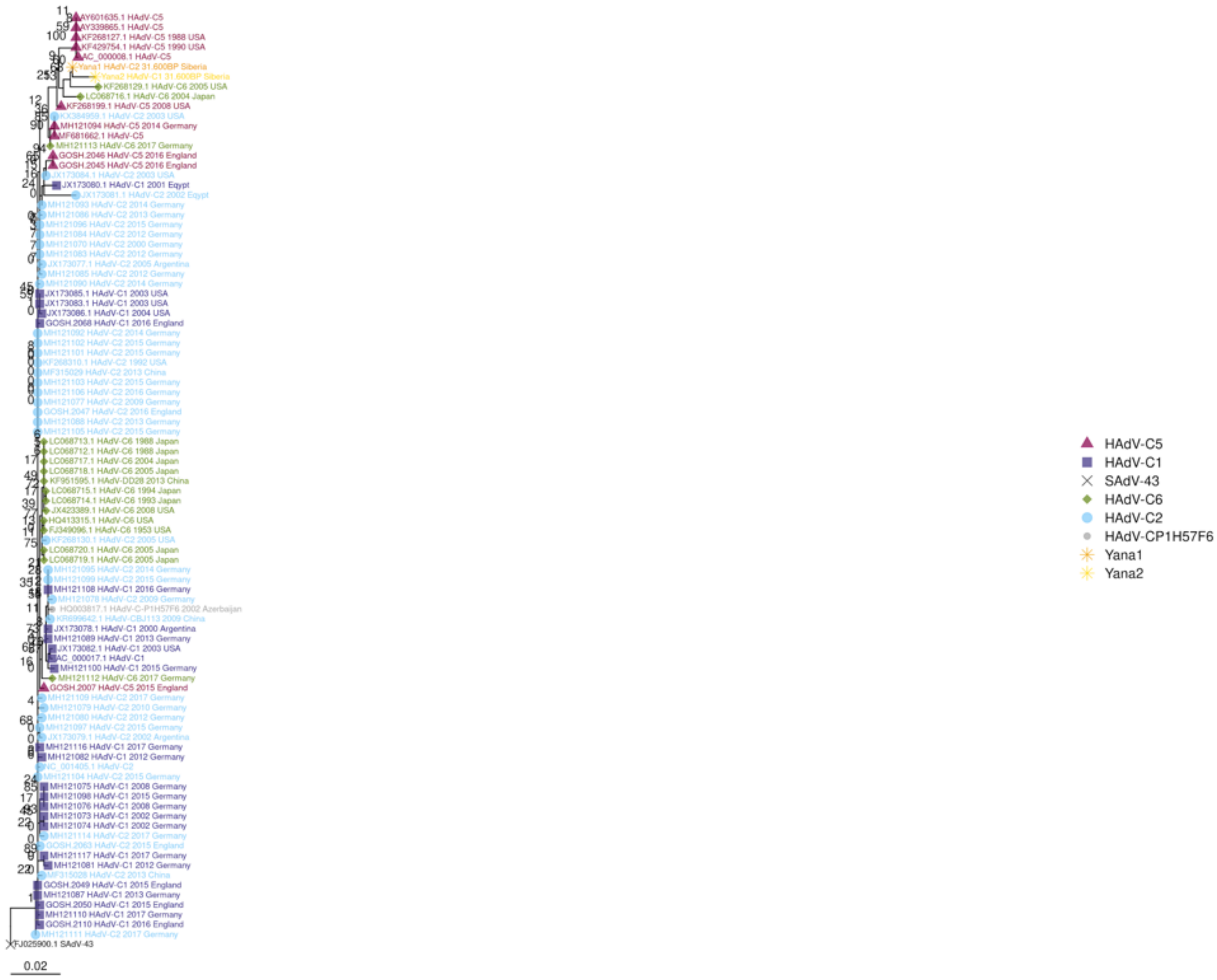

L1_IIIa_capsid_protein

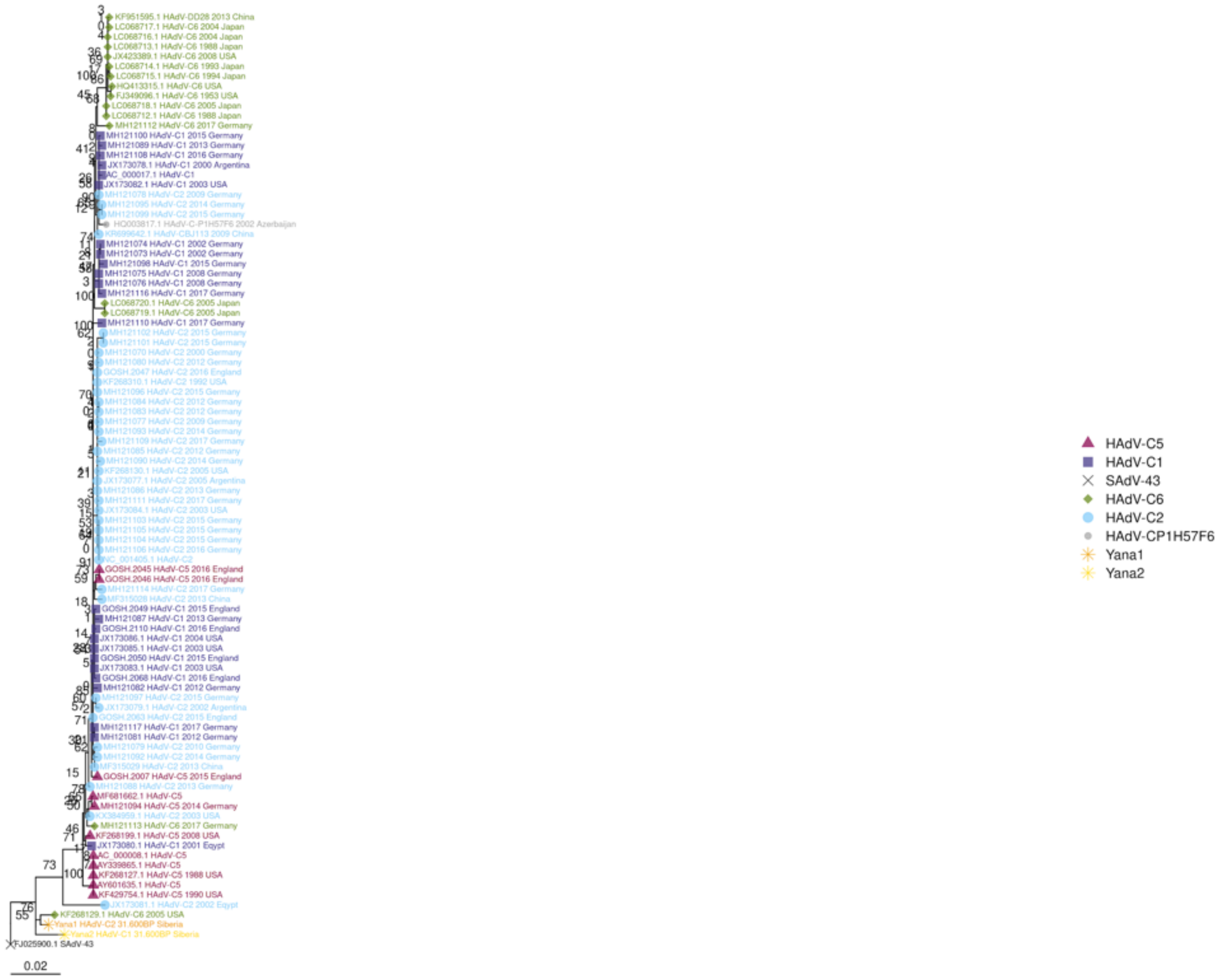

L2_penton_base

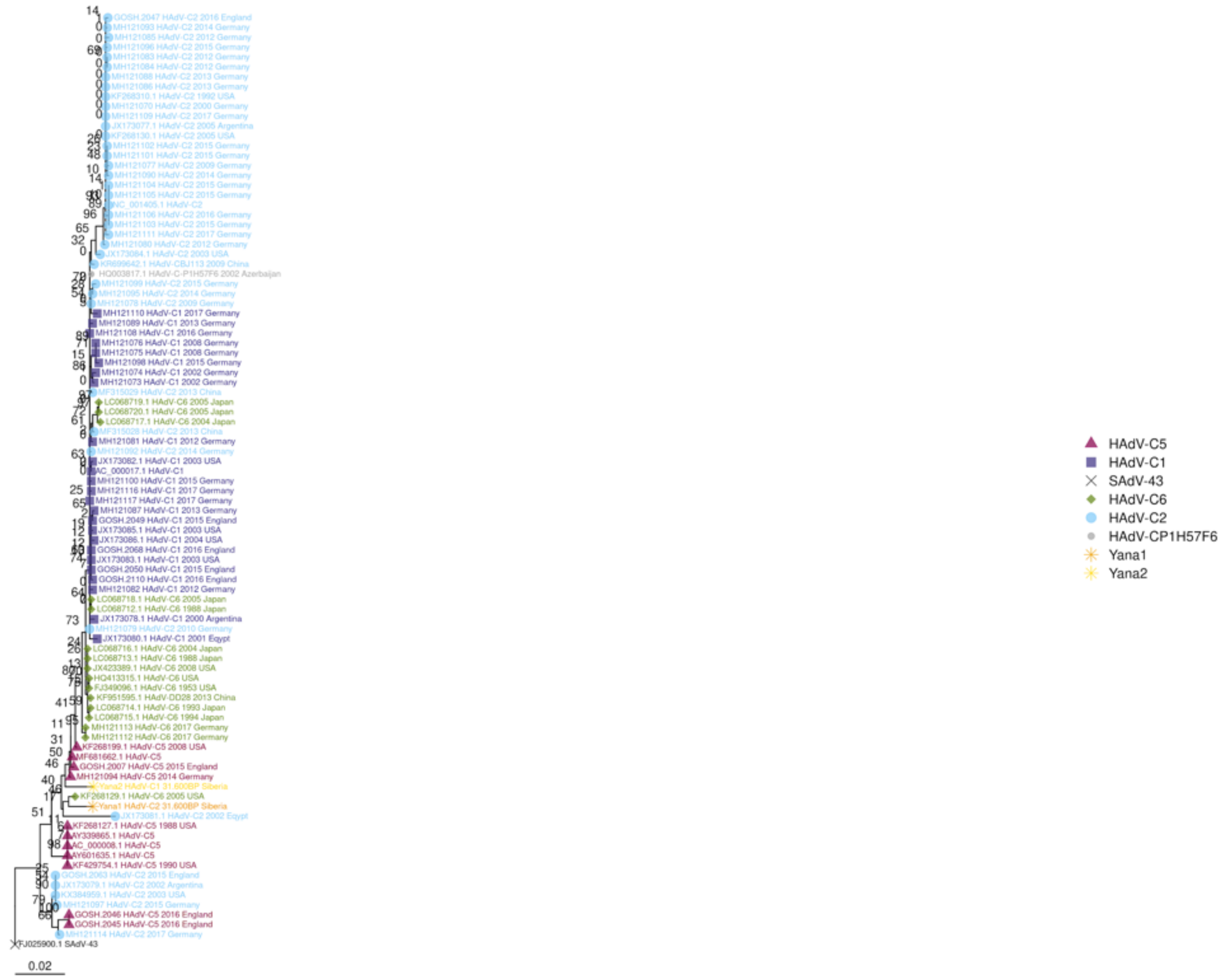

L2_V_core_protein

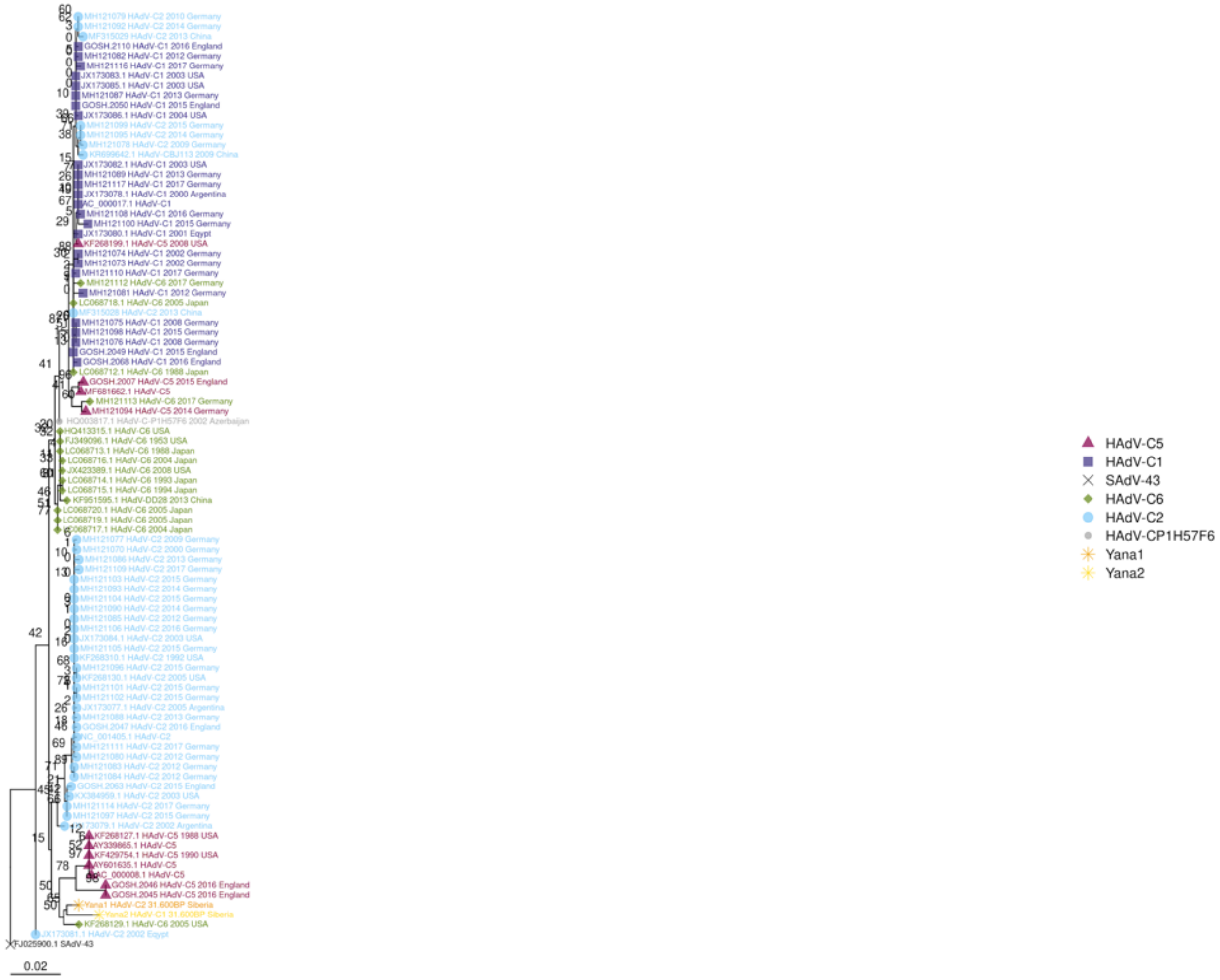

L2_VII_core_protein

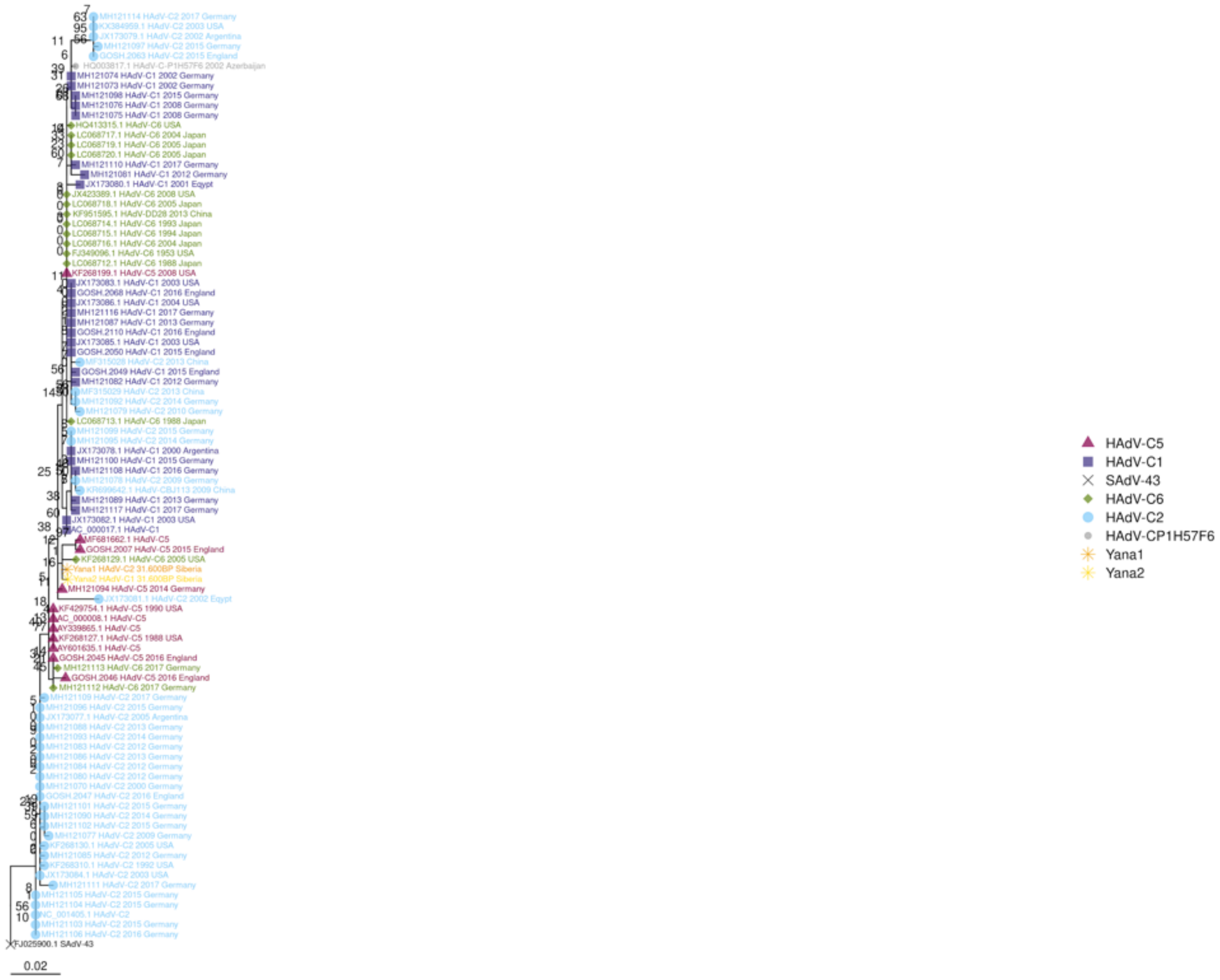

L2_X_core_protein

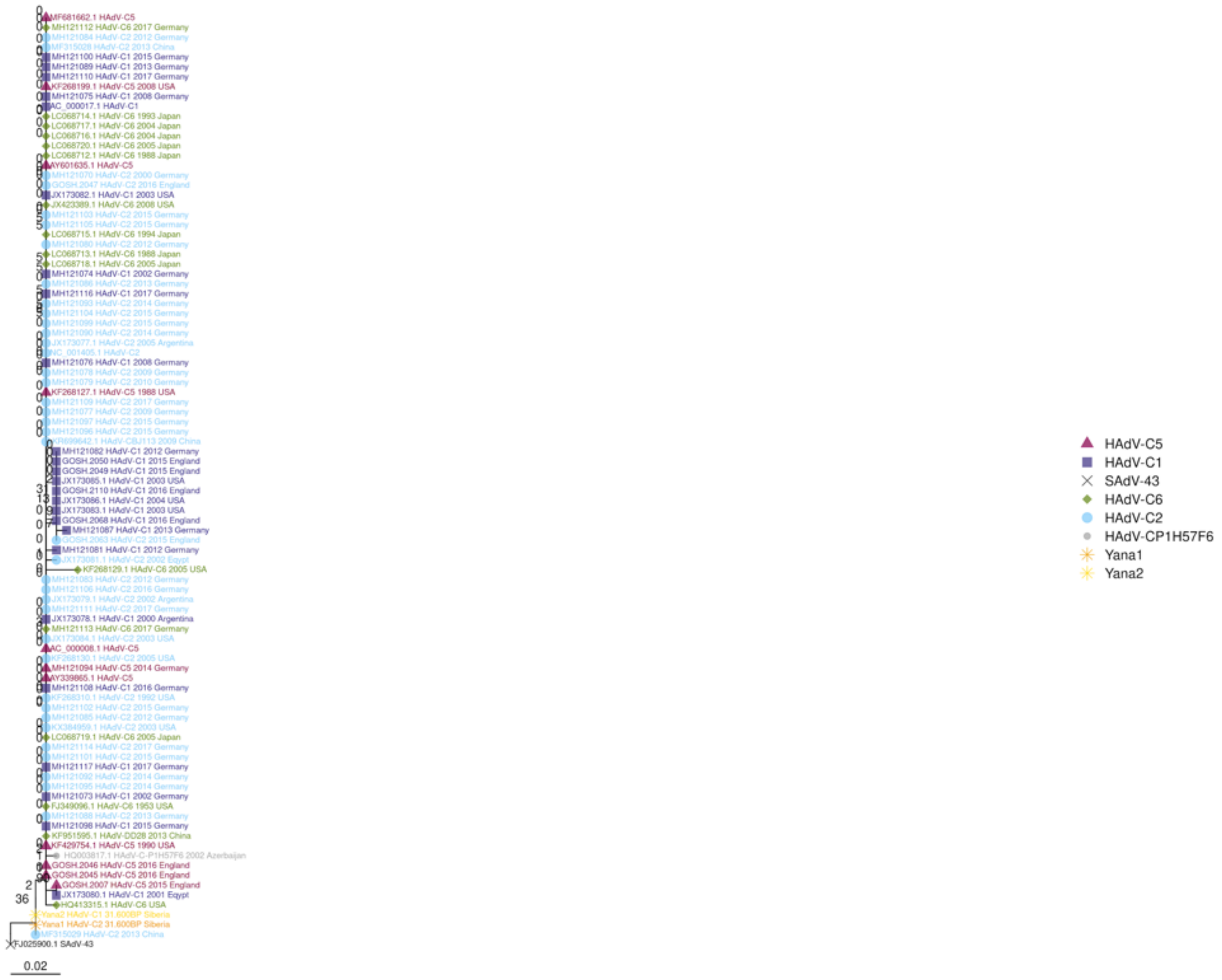

L3_hexon

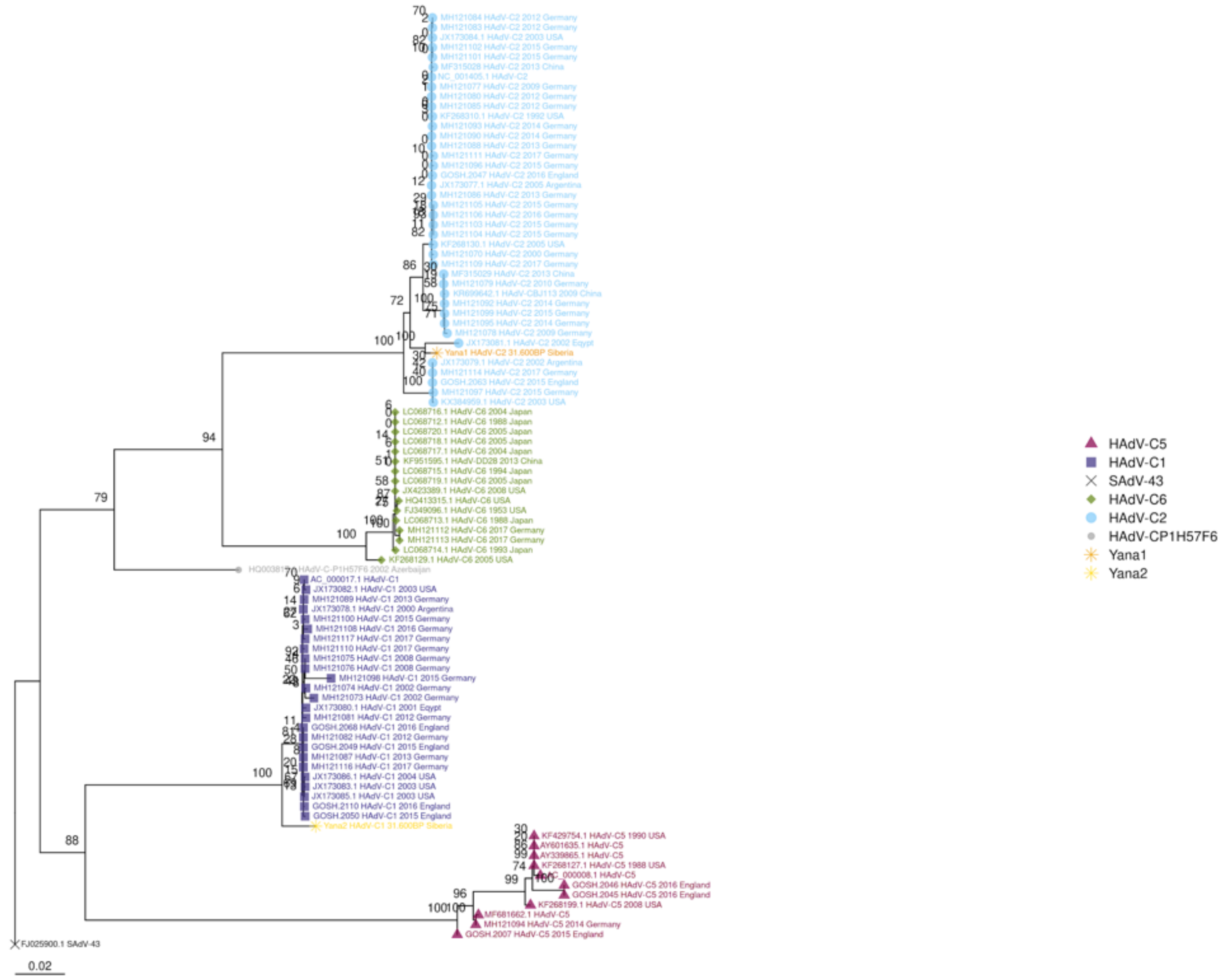
HAdV-C5
HAdV-C1
SAdV-43
HAdV-C6
HAdV-C2
HAdV-CP1H57F6
Yana1
Yana2
0.02

L3_protease

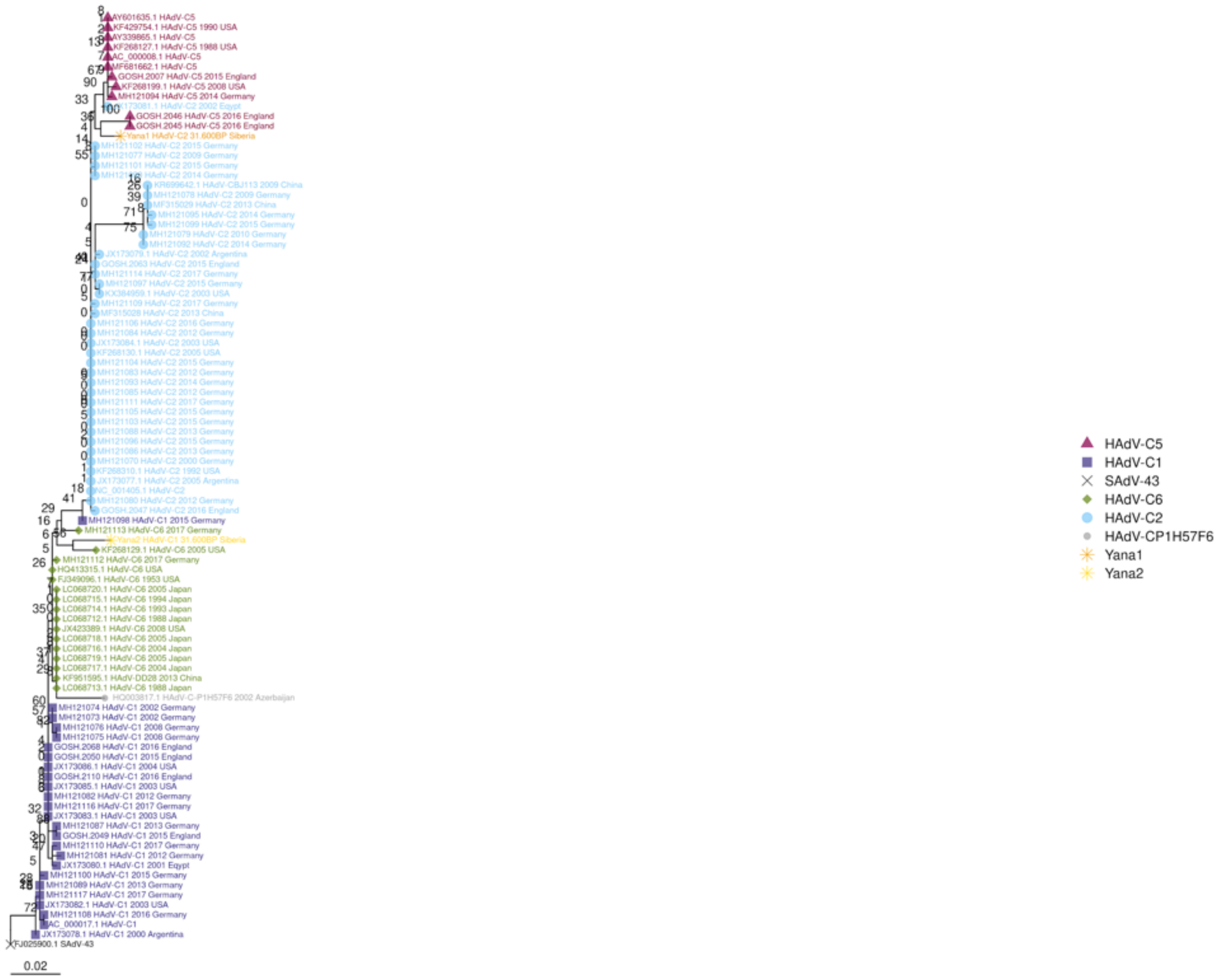
AY601635.1 HAdV-C5
KF429754.1 HAdV-C5 1990 USA
AY339865.1 HAdV-C5
KF268127.1 HAdV-C5 1988 USA
AC_000008.1 HAdV-C5
MF681662.1 HAdV-C5
GOSH.2007 HAdV-C5 2015 England
KF268199.1 HAdV-C5 2008 USA
MH121094 HAdV-C5 2014 Germany
X173081.1 HAdV-C2 2002 Egypt
GOSH.2046 HAdV-C5 2016 England
GOSH.2045 HAdV-C5 2016 England
Yana1 HAdV-C2 31.600BP Siberia
MH121102 HAdV-C2 2015 Germany
MH121077 HAdV-C2 2009 Germany
MH121101 HAdV-C2 2015 Germany
KR699642.1 HAdV-CBJ113 2009 China
MH121078 HAdV-C2 2009 Germany
MF315029 HAdV-C2 2013 China
MH121095 HAdV-C2 2014 Germany
MH121099 HAdV-C2 2015 Germany
MH121079 HAdV-C2 2010 Germany
MH121092 HAdV-C2 2014 Germany
JX173079.1 HAdV-C2 2002 Argentina
GOSH.2063 HAdV-C2 2015 England
MH121114 HAdV-C2 2017 Germany
MH121097 HAdV-C2 2015 Germany
KX384959.1 HAdV-C2 2003 USA
MH121109 HAdV-C2 2017 Germany
MF315028 HAdV-C2 2013 China
MH121106 HAdV-C2 2016 Germany
MH121084 HAdV-C2 2012 Germany
JX173084.1 HAdV-C2 2003 USA
KF268130.1 HAdV-C2 2005 USA
MH121104 HAdV-C2 2015 Germany
MH121083 HAdV-C2 2012 Germany
MH121093 HAdV-C2 2014 Germany
MH121085 HAdV-C2 2012 Germany
MH121111 HAdV-C2 2017 Germany
MH121105 HAdV-C2 2015 Germany
MH121103 HAdV-C2 2015 Germany
MH121088 HAdV-C2 2013 Germany
MH121096 HAdV-C2 2015 Germany
MH121086 HAdV-C2 2013 Germany
MH121070 HAdV-C2 2000 Germany
KF268310.1 HAdV-C2 1992 USA
JX173077.1 HAdV-C2 2005 Argentina
NC_001405.1 HAdV-C2
MH121080 HAdV-C2 2012 Germany
GOSH.2047 HAdV-C2 2016 England
MH121098 HAdV-C1 2015 Germany
MH121113 HAdV-C6 2017 Germany
Yana2 HAdV-C1 31.600BP Siberia
KF268129.1 HAdV-C6 2005 USA
MH121112 HAdV-C6 2017 Germany
HQ413315.1 HAdV-C6 USA
FJ349096.1 HAdV-C6 1953 USA
LC068720.1 HAdV-C6 2005 Japan
LC068715.1 HAdV-C6 1994 Japan
LC068714.1 HAdV-C6 1993 Japan
LC068712.1 HAdV-C6 1988 Japan
JX423389.1 HAdV-C6 2008 USA
LC068718.1 HAdV-C6 2005 Japan
LC068716.1 HAdV-C6 2004 Japan
LC068719.1 HAdV-C6 2005 Japan
LC068717.1 HAdV-C6 2004 Japan
KF951595.1 HAdV-DD28 2013 China
LC068713.1 HAdV-C6 1988 Japan
HQ003817.1 HAdV-C-P1H57F6 2002 Azerbaijan
MH121074 HAdV-C1 2002 Germany
MH121073 HAdV-C1 2002 Germany
MH121076 HAdV-C1 2008 Germany
MH121075 HAdV-C1 2008 Germany
GOSH.2068 HAdV-C1 2016 England
GOSH.2050 HAdV-C1 2015 England
JX173086.1 HAdV-C1 2004 USA
GOSH.2110 HAdV-C1 2016 England
JX173085.1 HAdV-C1 2003 USA
MH121082 HAdV-C1 2012 Germany
MH121116 HAdV-C1 2017 Germany
JX173083.1 HAdV-C1 2003 USA
MH121087 HAdV-C1 2013 Germany
GOSH.2049 HAdV-C1 2015 England
MH121110 HAdV-C1 2017 Germany
MH121081 HAdV-C1 2012 Germany
JX173080.1 HAdV-C1 2001 Egypt
MH121100 HAdV-C1 2015 Germany
MH121089 HAdV-C1 2013 Germany
MH121117 HAdV-C1 2017 Germany
JX173082.1 HAdV-C1 2003 USA
MH121108 HAdV-C1 2016 Germany
AC_000017.1 HAdV-C1
JX173078.1 HAdV-C1 2000 Argentina
FJ025900.1 SAdV-43
0.02
HAdV-C5
HAdV-C1
SAdV-43
HAdV-C6
HAdV-C2
HAdV-CP1H57F6
Yana1
Yana2

L3_VI_capsid_protein

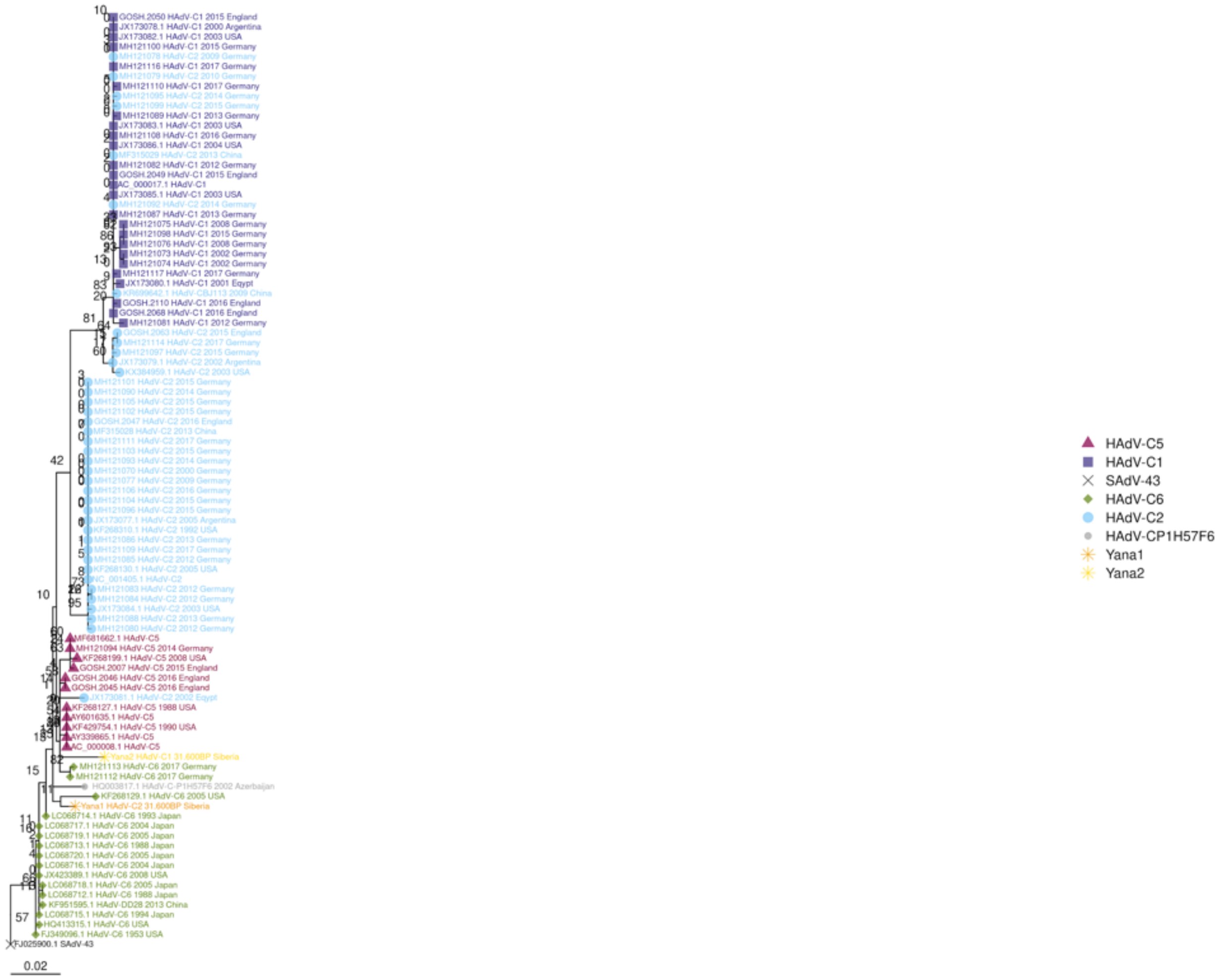

L4_encapsidation_protein

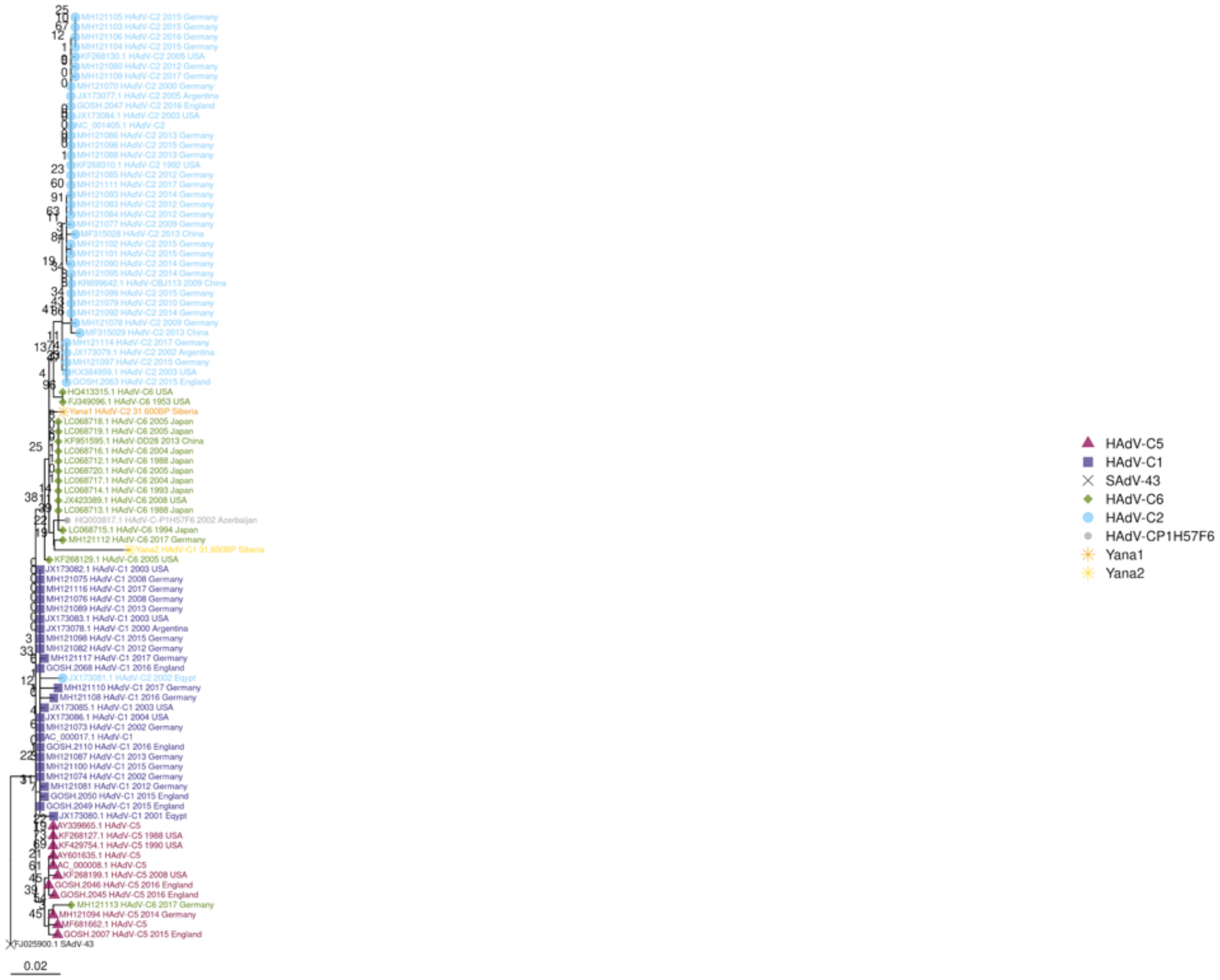

L4_hexon-assembly_protein

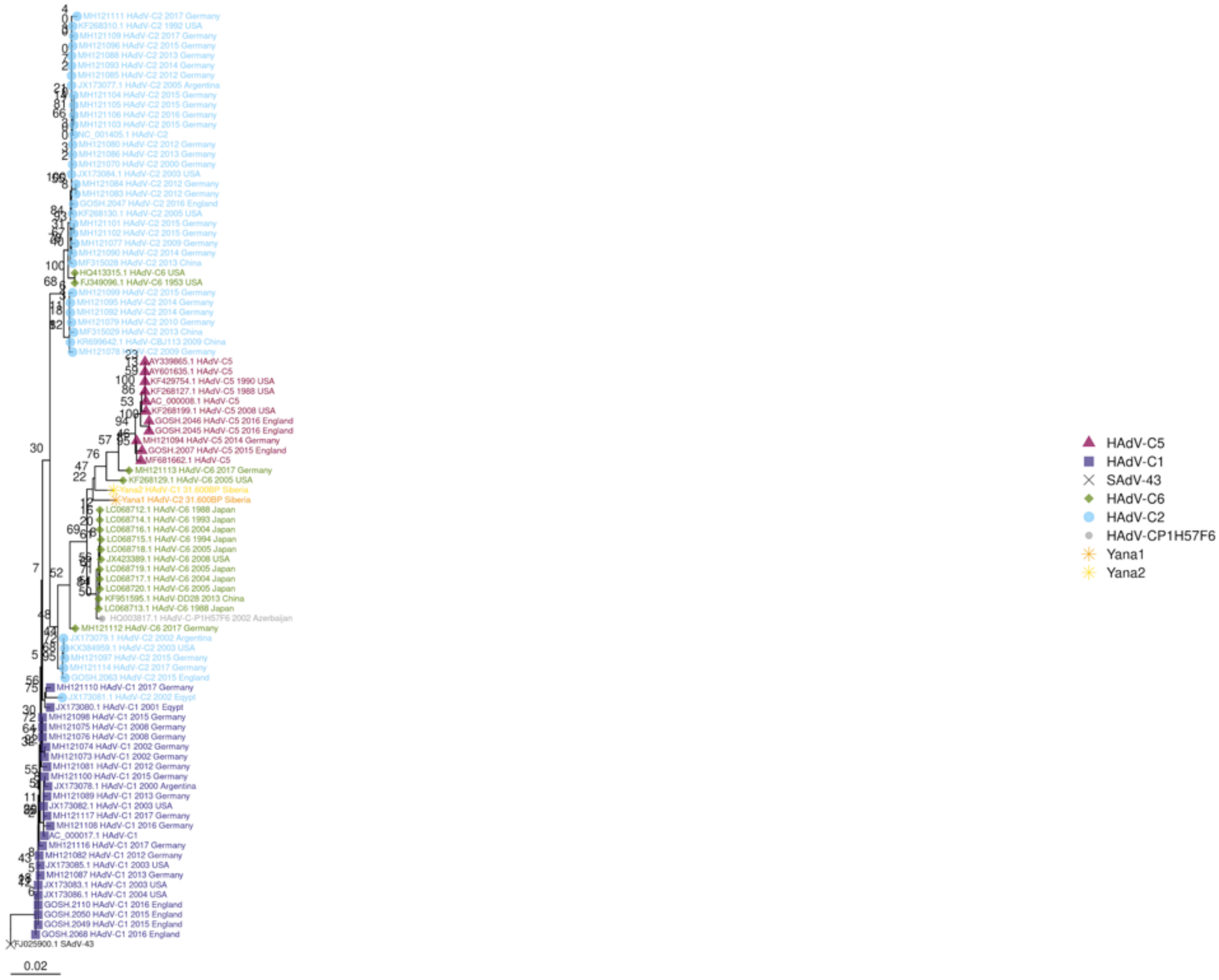

L4_splicing_factor

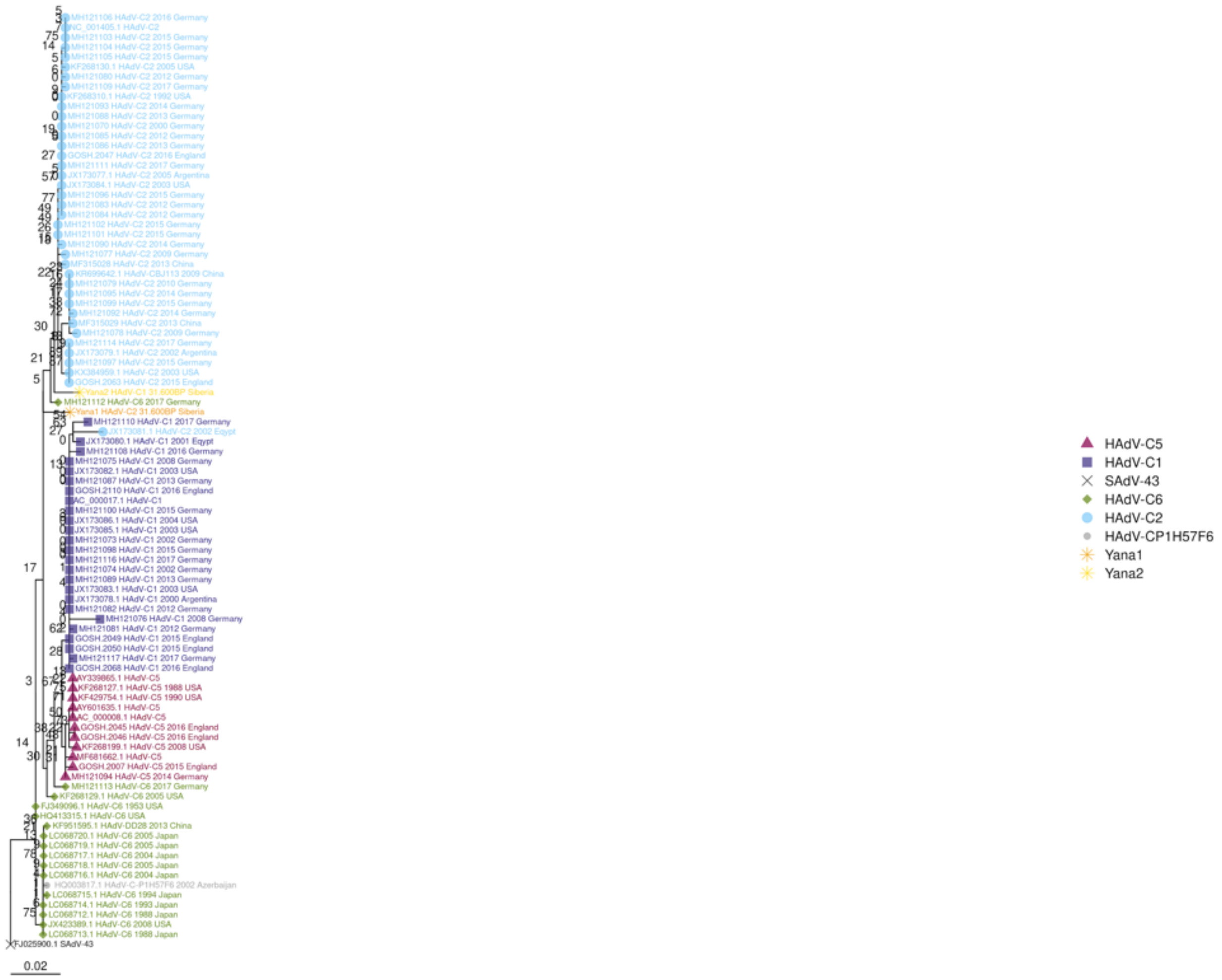

L4_VIII_capsid_protein

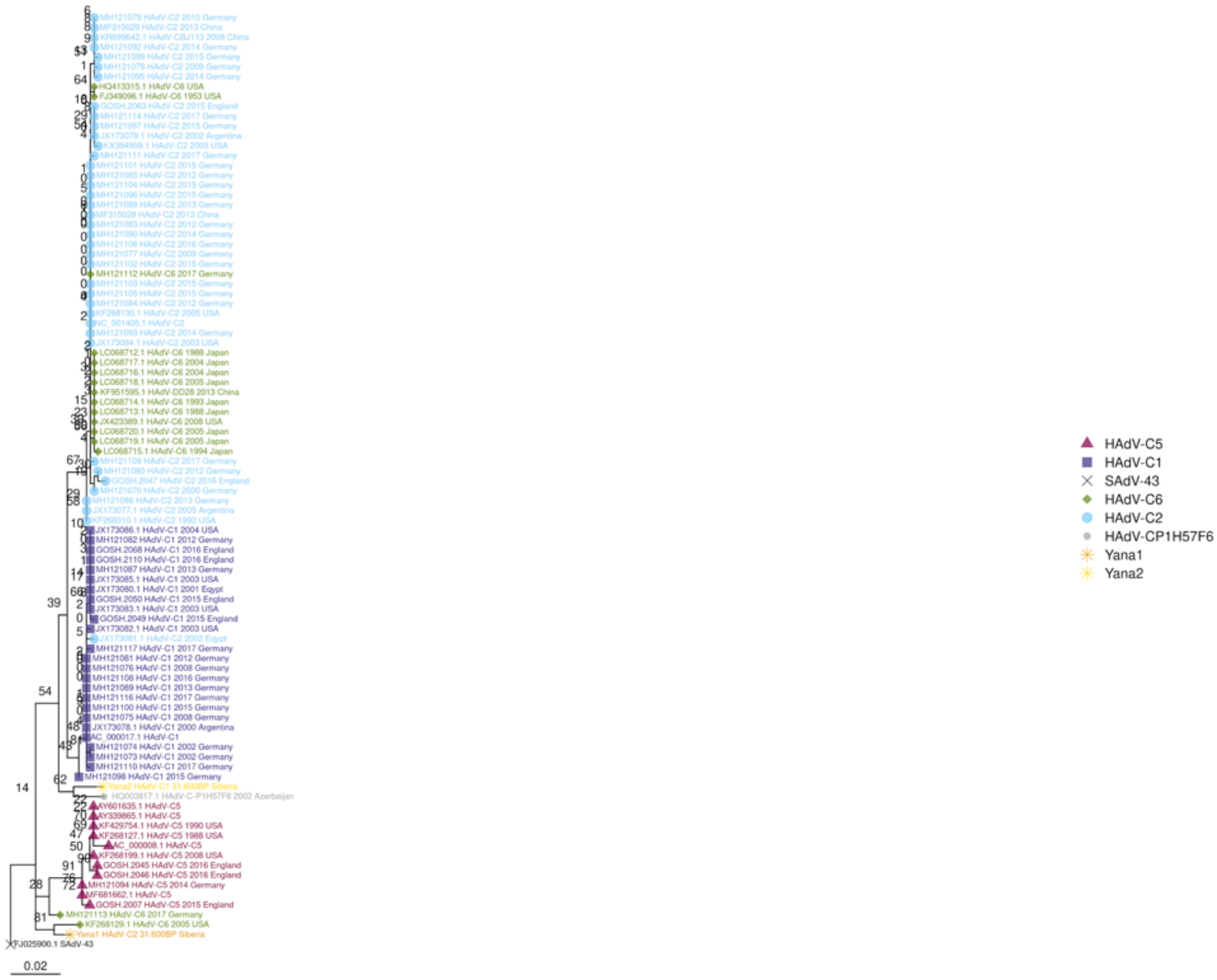

L5_fiber

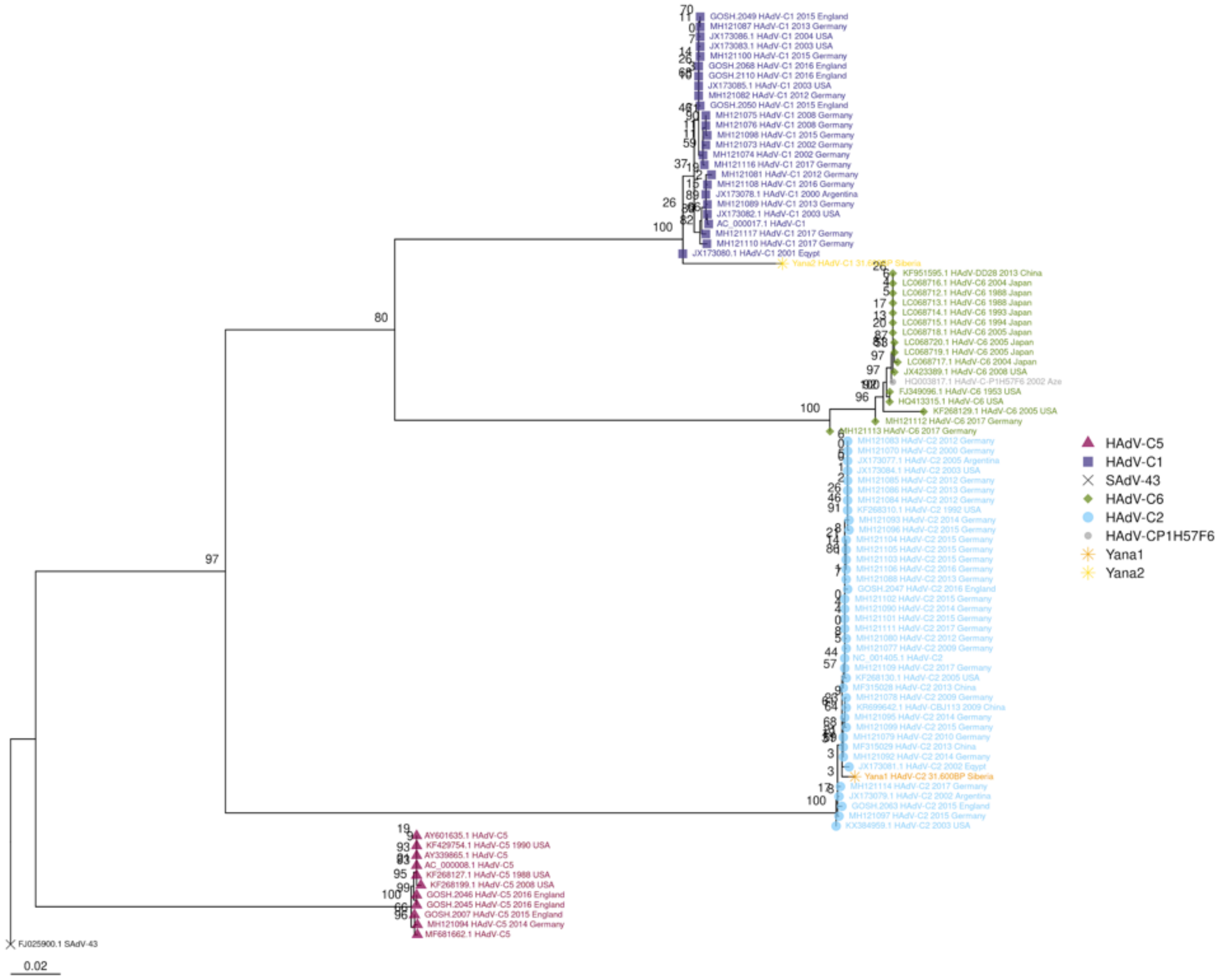
GOSH.2049 HAdV-C1 2015 England
MH121087 HAdV-C1 2013 Germany
JX173086.1 HAdV-C1 2004 USA
JX173083.1 HAdV-C1 2003 USA
MH121100 HAdV-C1 2015 Germany
GOSH.2068 HAdV-C1 2016 England
GOSH.2110 HAdV-C1 2016 England
JX173085.1 HAdV-C1 2003 USA
MH121082 HAdV-C1 2012 Germany
GOSH.2050 HAdV-C1 2015 England
MH121075 HAdV-C1 2008 Germany
MH121076 HAdV-C1 2008 Germany
MH121098 HAdV-C1 2015 Germany
MH121073 HAdV-C1 2002 Germany
MH121074 HAdV-C1 2002 Germany
MH121116 HAdV-C1 2017 Germany
MH121081 HAdV-C1 2012 Germany
MH121108 HAdV-C1 2016 Germany
JX173078.1 HAdV-C1 2000 Argentina
MH121089 HAdV-C1 2013 Germany
JX173082.1 HAdV-C1 2003 USA
AC_000017.1 HAdV-C1
MH121117 HAdV-C1 2017 Germany
MH121110 HAdV-C1 2017 Germany
JX173080.1 HAdV-C1 2001 Eqypt
Yana2 HAdV-C1 31.6 P Siberia
KF951595.1 HAdV-DD28 2013 China
LC068716.1 HAdV-C6 2004 Japan
LC068712.1 HAdV-C6 1988 Japan
LC068713.1 HAdV-C6 1988 Japan
LC068714.1 HAdV-C6 1993 Japan
LC068715.1 HAdV-C6 1994 Japan
LC068718.1 HAdV-C6 2005 Japan
LC068720.1 HAdV-C6 2005 Japan
LC068719.1 HAdV-C6 2005 Japan
LC068717.1 HAdV-C6 2004 Japan
JX423389.1 HAdV-C6 2008 USA
HQ003817.1 HAdV-C-P1H57F6 2002 Aze
FJ349096.1 HAdV-C6 1953 USA
HQ413315.1 HAdV-C6 USA
KF268129.1 HAdV-C6 2005 USA
MH121112 HAdV-C6 2017 Germany
MH121113 HAdV-C6 2017 Germany
MH121083 HAdV-C2 2012 Germany
MH121070 HAdV-C2 2000 Germany
JX173077.1 HAdV-C2 2005 Argentina
JX173084.1 HAdV-C2 2003 USA
MH121085 HAdV-C2 2012 Germany
MH121086 HAdV-C2 2013 Germany
MH121084 HAdV-C2 2012 Germany
KF268310.1 HAdV-C2 1992 USA
MH121093 HAdV-C2 2014 Germany
MH121096 HAdV-C2 2015 Germany
MH121104 HAdV-C2 2015 Germany
MH121105 HAdV-C2 2015 Germany
MH121103 HAdV-C2 2015 Germany
MH121106 HAdV-C2 2016 Germany
MH121088 HAdV-C2 2013 Germany
GOSH.2047 HAdV-C2 2016 England
MH121102 HAdV-C2 2015 Germany
MH121090 HAdV-C2 2014 Germany
MH121101 HAdV-C2 2015 Germany
MH121111 HAdV-C2 2017 Germany
MH121080 HAdV-C2 2012 Germany
MH121077 HAdV-C2 2009 Germany
NC_001405.1 HAdV-C2
MH121109 HAdV-C2 2017 Germany
KF268130.1 HAdV-C2 2005 USA
MF315028 HAdV-C2 2013 China
MH121078 HAdV-C2 2009 Germany
KR699642.1 HAdV-CBJ113 2009 China
MH121095 HAdV-C2 2014 Germany
MH121099 HAdV-C2 2015 Germany
MH121079 HAdV-C2 2010 Germany
MF315029 HAdV-C2 2013 China
MH121092 HAdV-C2 2014 Germany
JX173081.1 HAdV-C2 2002 Eqypt
Yana1 HAdV-C2 31,600BP Siberia
MH121114 HAdV-C2 2017 Germany
JX173079.1 HAdV-C2 2002 Argentina
GOSH.2063 HAdV-C2 2015 England
MH121097 HAdV-C2 2015 Germany
KX384959.1 HAdV-C2 2003 USA
AY601635.1 HAdV-C5
KF429754.1 HAdV-C5 1990 USA
AY339865.1 HAdV-C5
AC_000008.1 HAdV-C5
KF268127.1 HAdV-C5 1988 USA
KF268199.1 HAdV-C5 2008 USA
GOSH.2046 HAdV-C5 2016 England
GOSH.2045 HAdV-C5 2016 England
GOSH.2007 HAdV-C5 2015 England
MH121094 HAdV-C5 2014 Germany
MF681662.1 HAdV-C5
FJ025900.1 SAdV-43
0.02
80
97
100
100
100
HAdV-C5
HAdV-C1
SAdV-43
HAdV-C6
HAdV-C2
HAdV-CP1H57F6
Yana1
Yana2

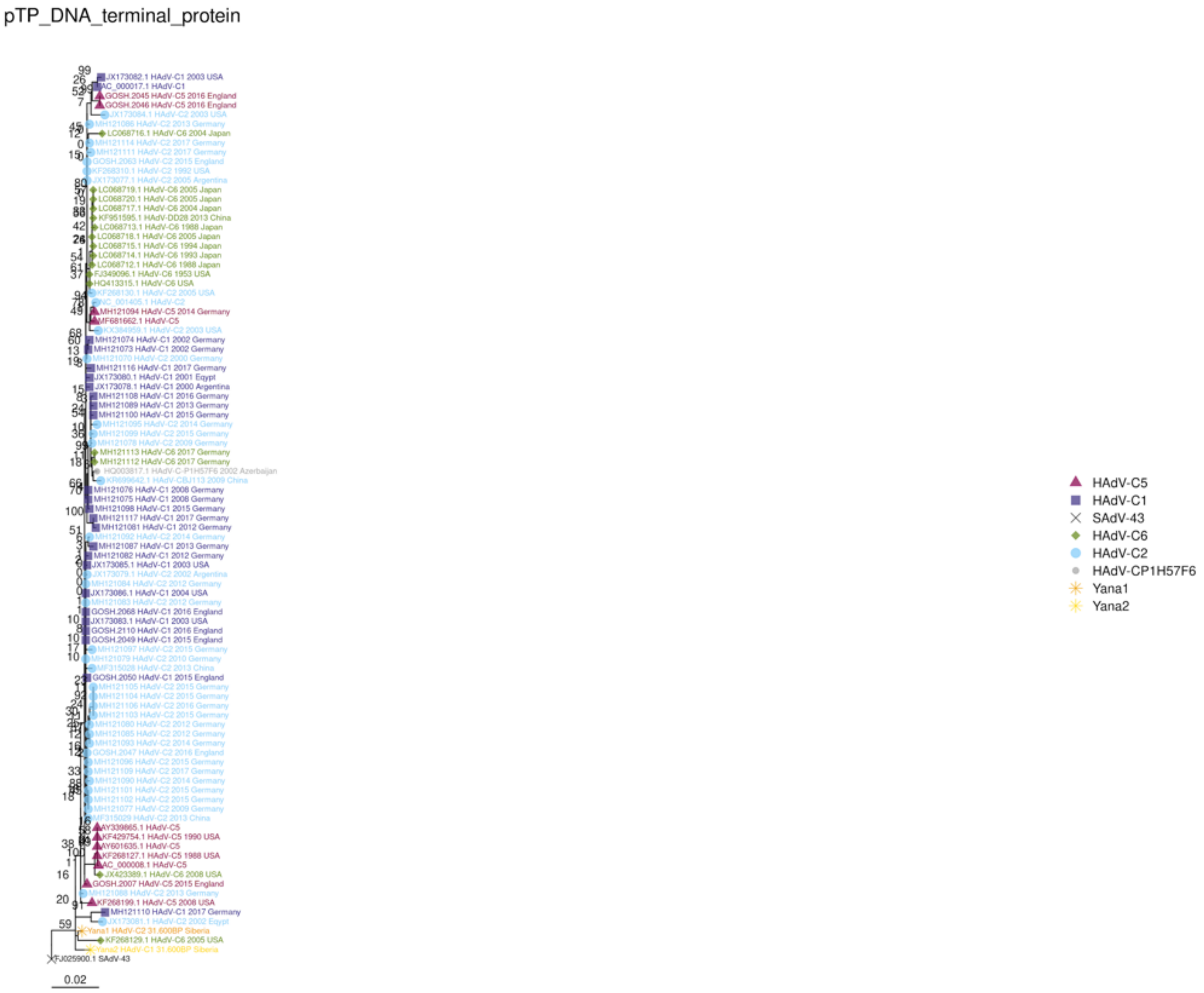

**Fig. S29. Maximum likelihood coding sequence (cds) trees**, model GTR+G-I, outgroup branch removed for visualization.

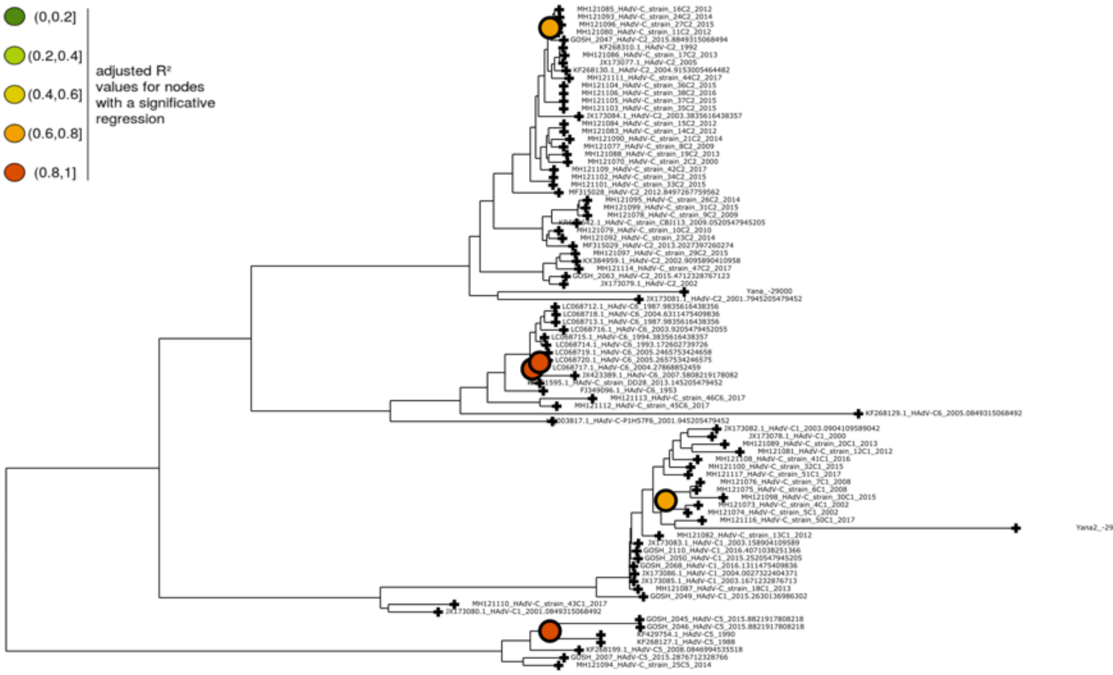

**Fig. S30. Linear regression full ML tree,** using PhyloSTemS, no significant regressions at the base of any clade.

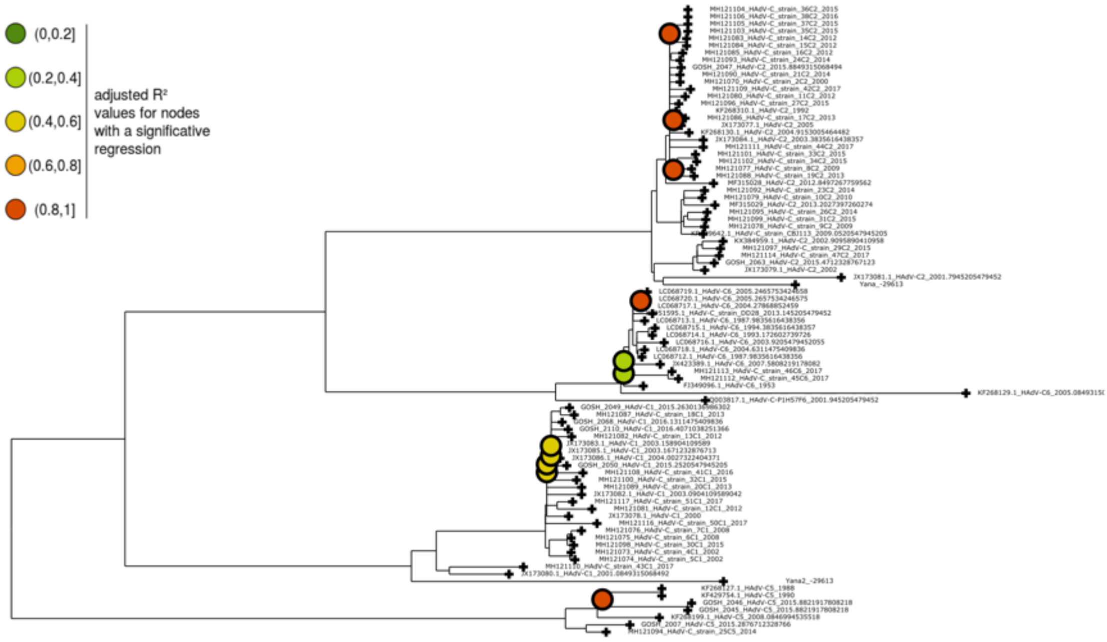

**Fig. S31. Linear regression, ML tree with homoplasies removed**, no significant regression at the base of any clade, using PhyloSTemS.

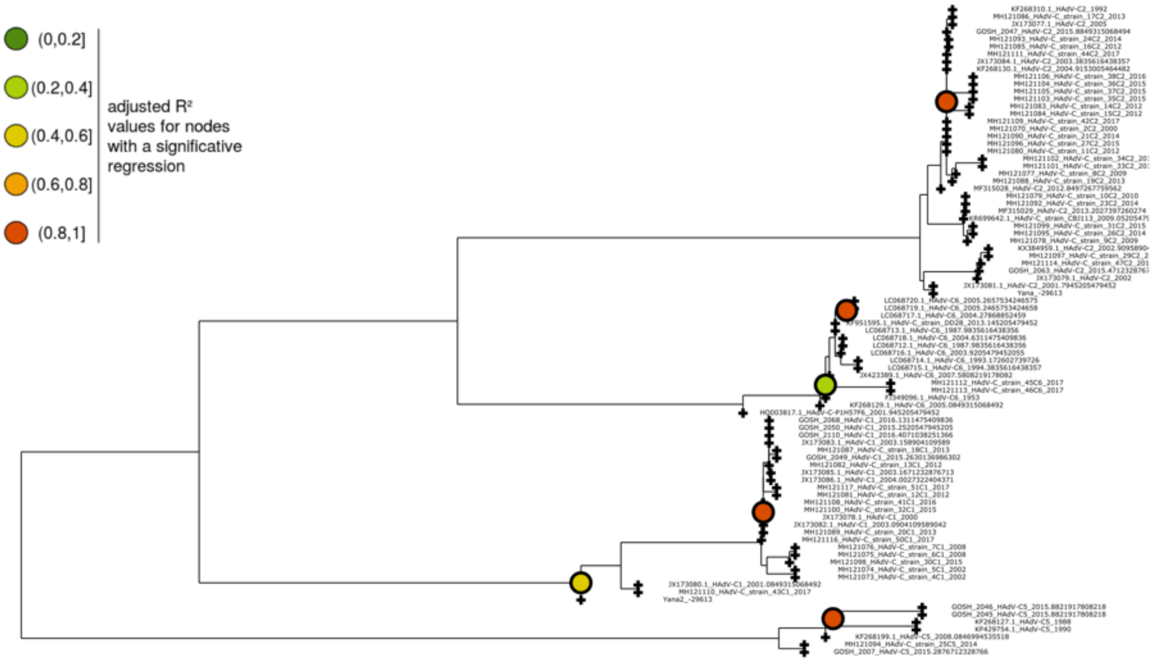

**Fig. S32. Linear regression, ML tree with both homoplasies and singletons removed**, significant regression at the base of the clade of HAdV-C1, using PhyloSTemS.

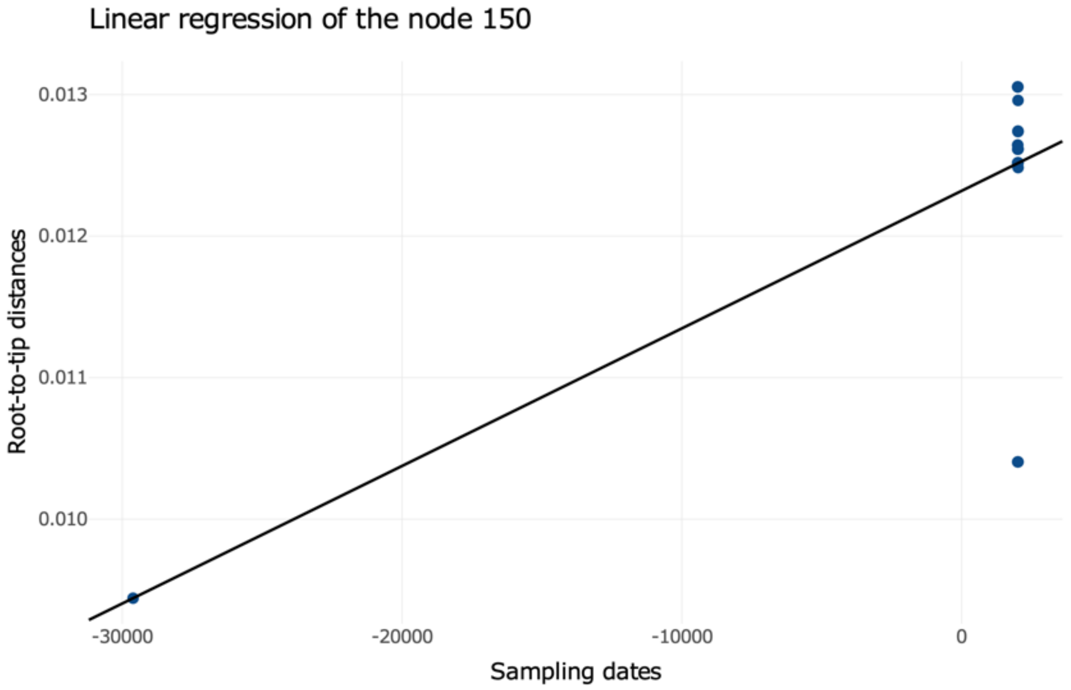

y = 0.012 + 9.7e-08 x
p-value = 0.000185
adjusted $R^2$ = 0.44
x-intercept = -123711.34

**Fig. S33. Linear regression at node of Yana2,** ML tree with both homoplasies and singletons removed

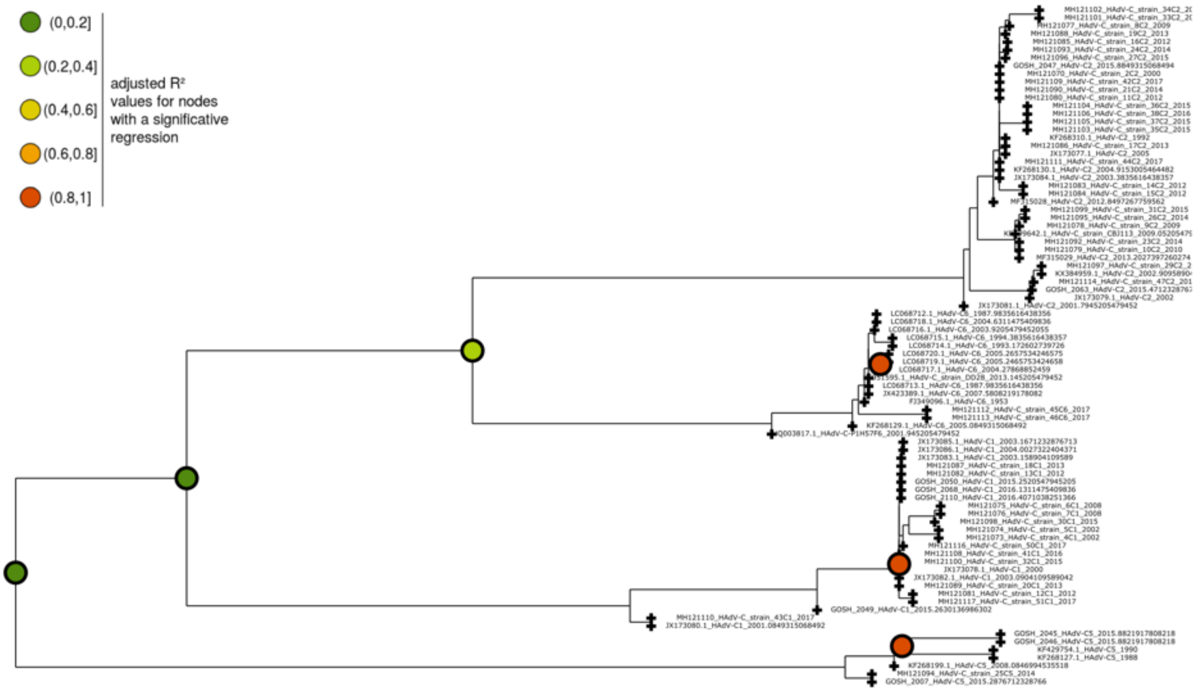

**Fig. S34. Linear regression, ML tree without ancient samples,** the significant regression at the clade including Yana2 before disappears, using PhyloSTemS.

Rate=2.62e−04,MRCA=−225432.84,R2=0.04,p=3.43e−02

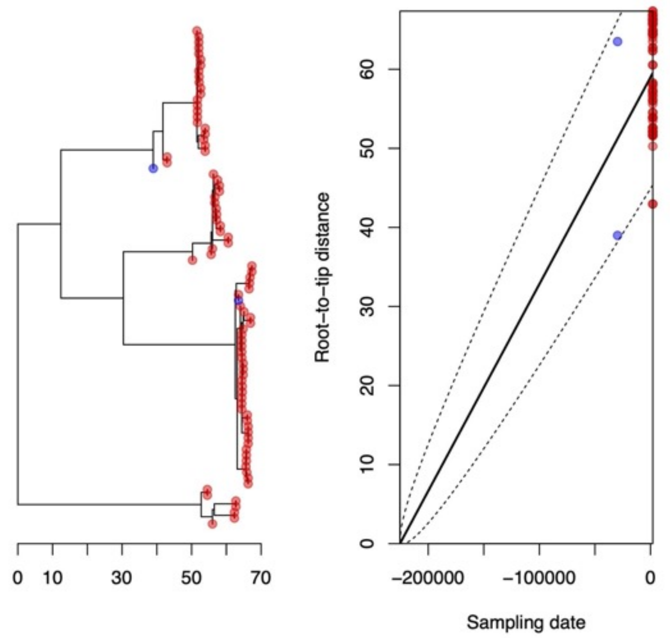

**Fig. S35. Root to tip regression,** using BactDating, including ancient samples.

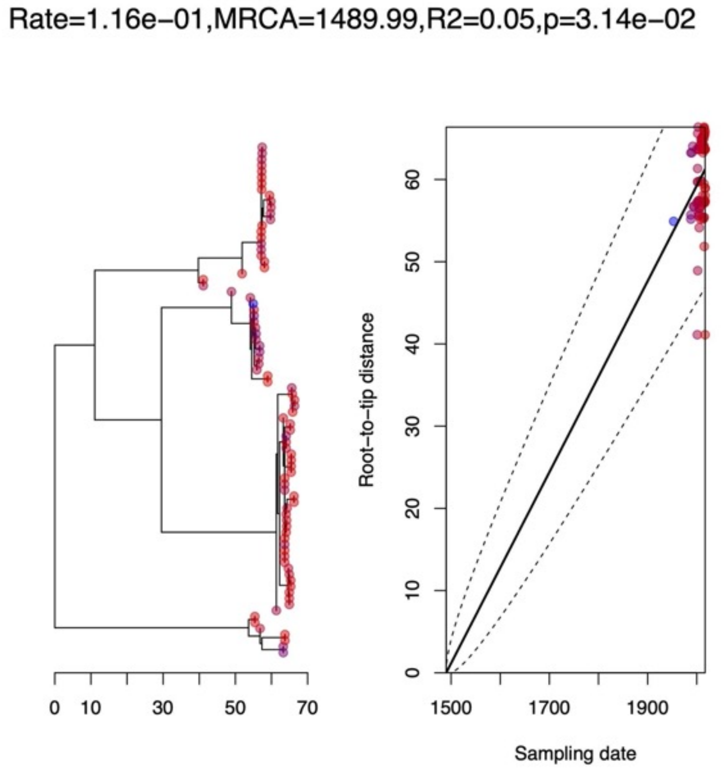

**Fig. S36. Root to tip regression,** using BactDating without ancient samples

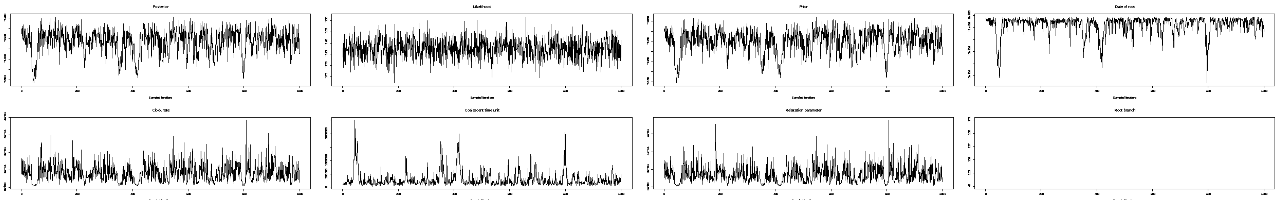

**Fig. S37: Convergence of BactDate**. Seven million iterations, including ancient samples.

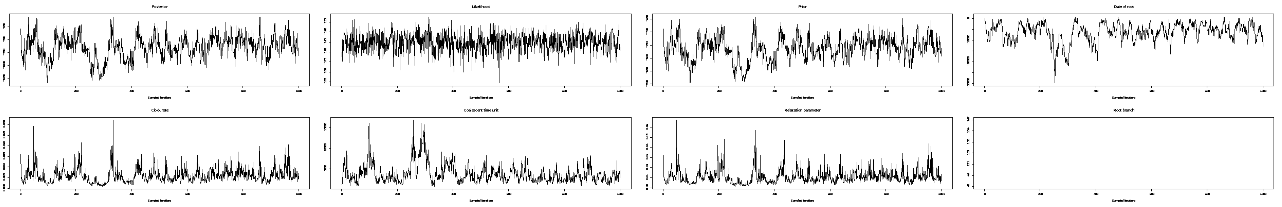

**Fig. S38. Convergence of BactDate**. Seven million iterations, without ancient samples.

**File: snps.log item: substmodel**

Models with blue circles are inside 95%HPD, red outside, and without circles have at most 1,14% support.

| posterior support | cumulative support | model |
|---|---|---|
| 42,42% | 42,42% | 121323 |
| 28,25% | 70,67% | 121343 |
| 12,25% | 82,93% | 123424 |
| 8,63% | 91,56% | 123454 |
| 3,90% | 95,46% | 121324 |
| 2,61% | 98,08% | 121345 |
| 1,14% | 99,22% | 123425 |
| 0,78% | 100,00% | 123456 |

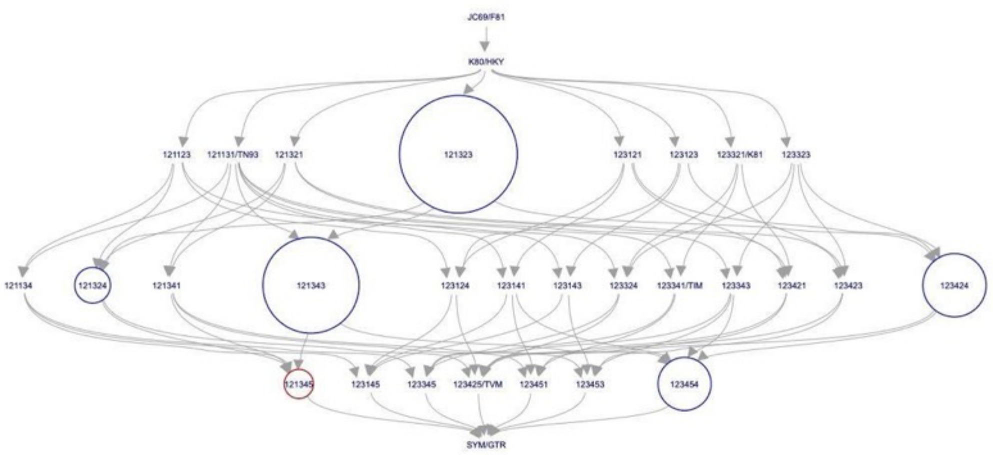

**Fig. S39. BModelTest,** substitution model illustration, no model has above 50% support.

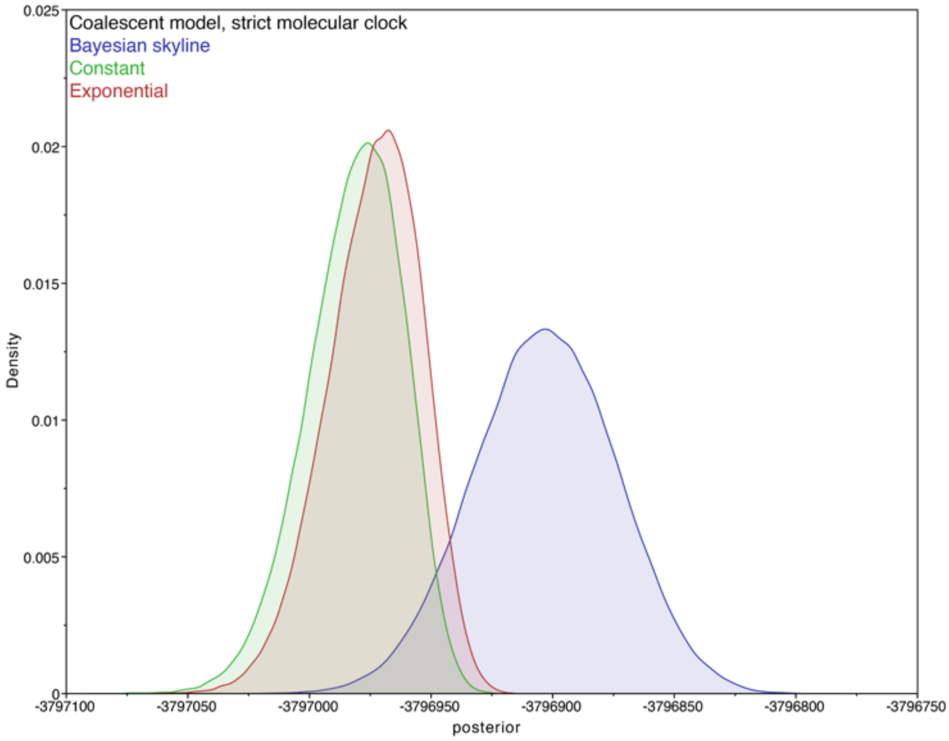

**Fig. S40. BEAST2 Posterior**, marginal density of three of the coalescent models, with strict molecular clock

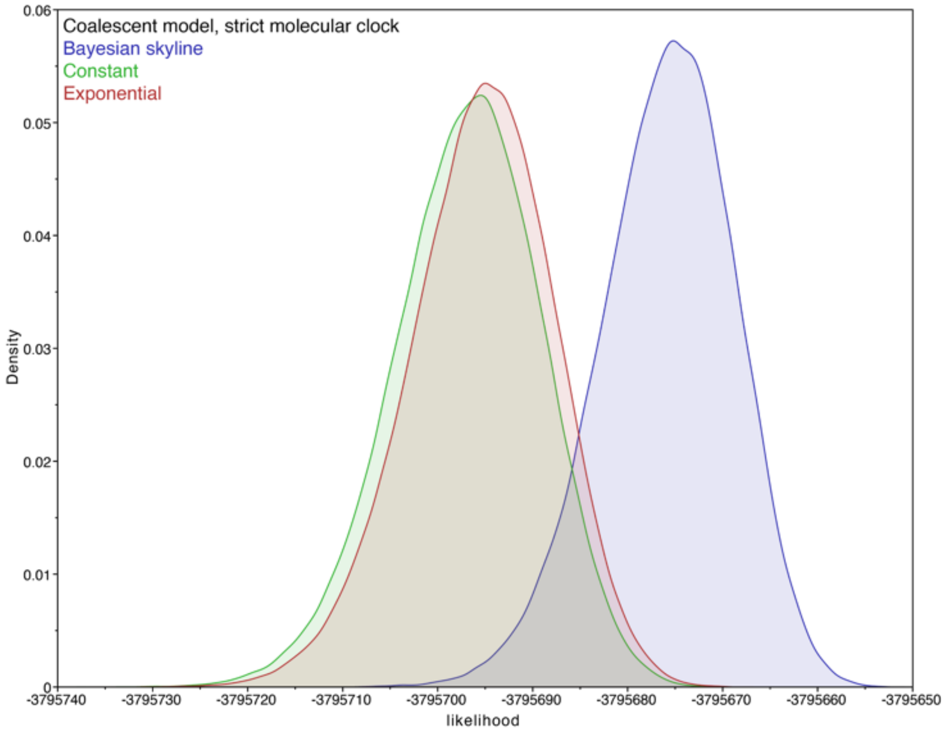

**Fig. S41. BEAST2 Likelihood**, marginal density of three of the coalescent models, with strict molecular clock, Bayesian skyline has the highest.

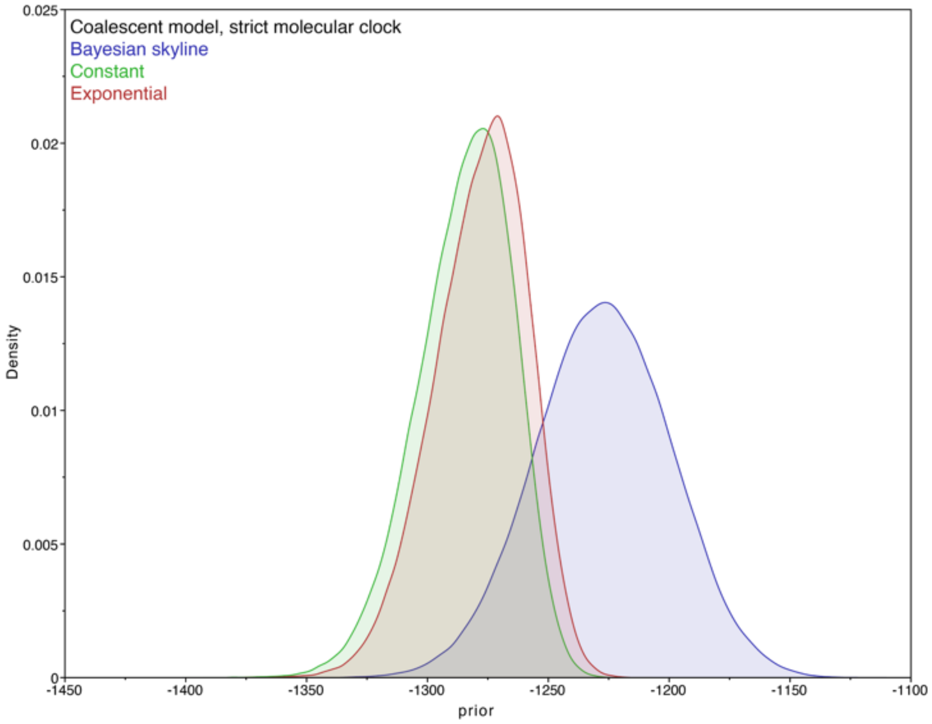

**Fig. S42. BEAST2 prior**, marginal density of three of the coalescent models, with strict molecular clock

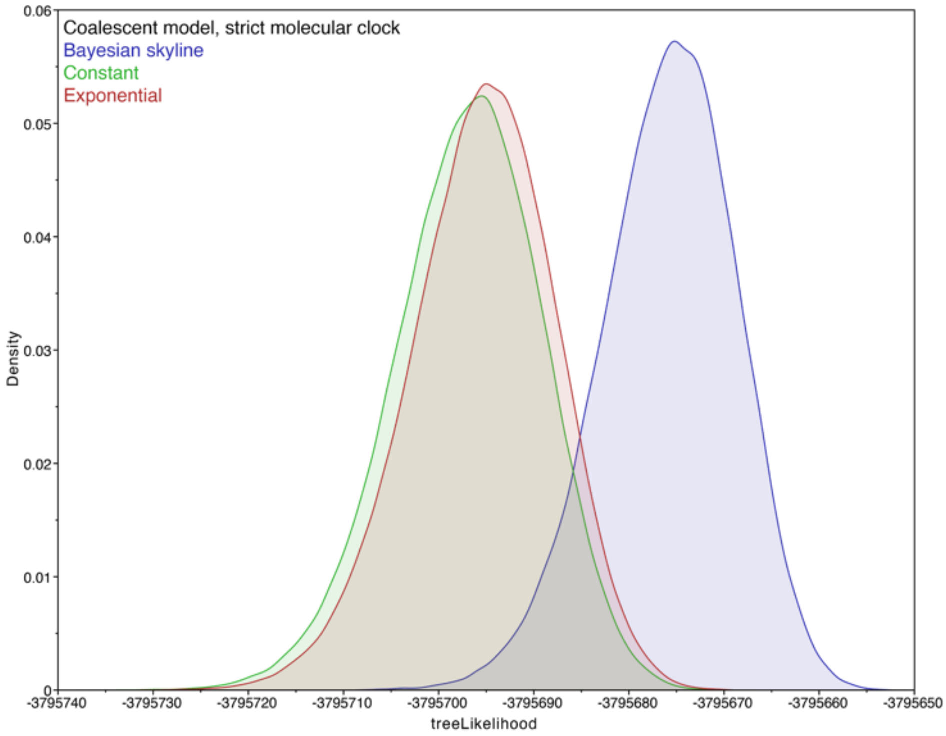

**Fig. S43. BEAST2 tree likelihood**, marginal density of three of the coalescent models, with strict molecular clock

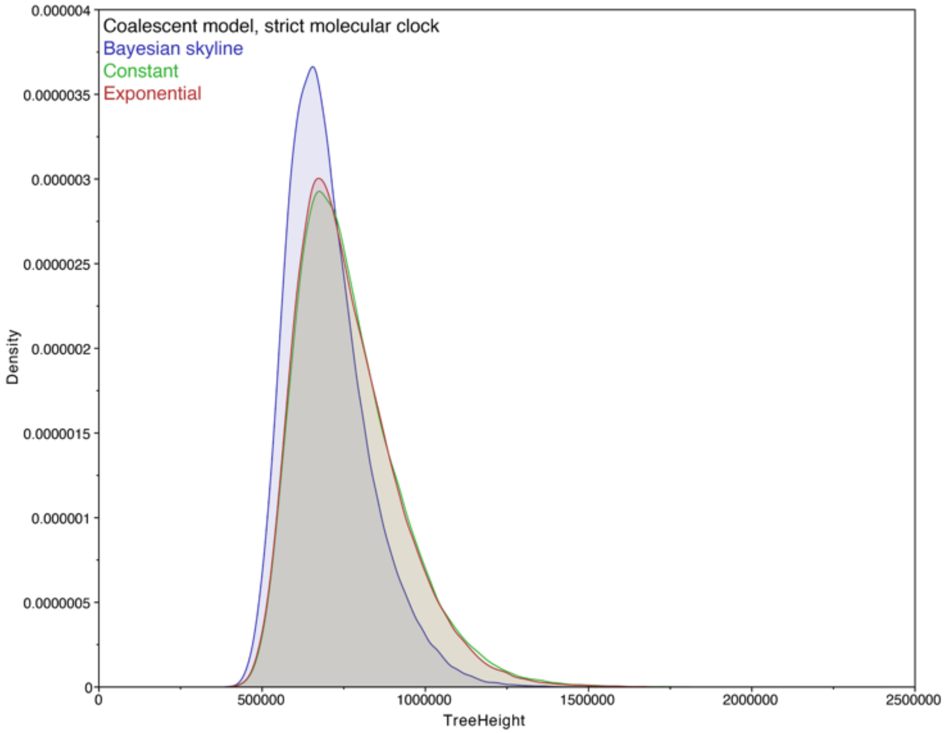

**Fig. S44. BEAST2 tree height** (divergence date), marginal density of three of the coalescent models, with strict molecular clock, very similar estimate with all three models.

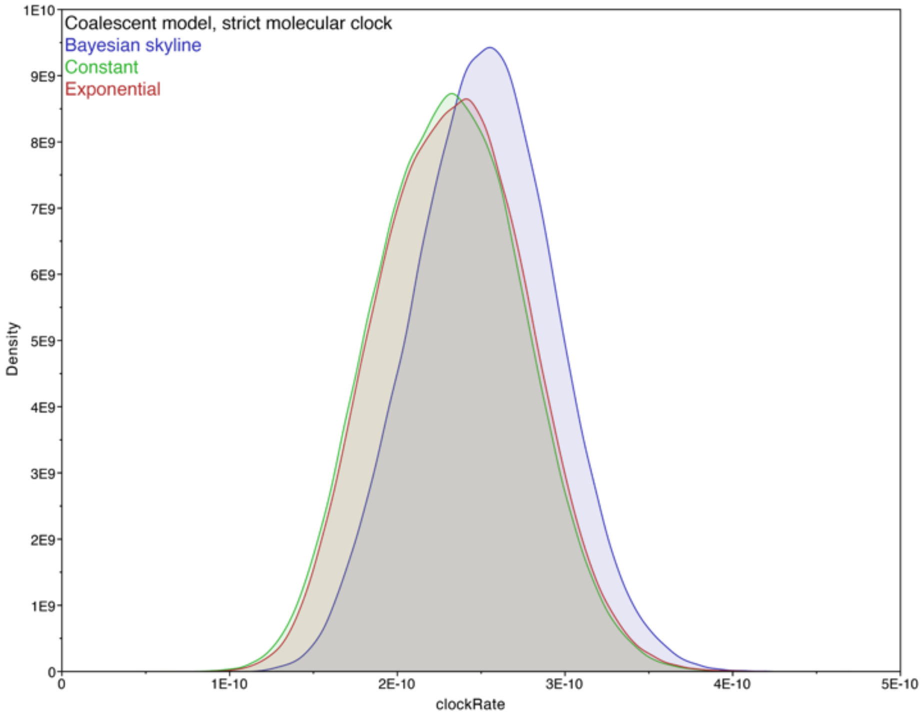

**Fig. S45. BEAST2 clock rate**, marginal density of three of the coalescent models, with strict molecular clock, similar estimate with all three models.

**Table S1.** Kraken, metagenomic classification results for adenovirus for each sample library.

| Sample | Percent of all reads classified | Number of reads classified on this taxonomic level and below | Number of reads classified on this taxonomic level | Taxonomic level | TaxId | Scientific name |
|---|---|---|---|---|---|---|
| Yana_L5 | 0 | 3151 | 20 | G | 10509 | Mastadenovirus |
| Yana_L5 | 0 | 3112 | 2587 | S | 129951 | Human mastadenovirus C |
| Yana_L5 | 0 | 264 | 264 | - | 28285 | Human adenovirus 5 |
| Yana_L5 | 0 | 261 | 261 | - | 10533 | Human adenovirus 1 |
| Yana_L5 | 0 | 10 | 10 | S | 130310 | Human mastadenovirus D |
| Yana_L5 | 0 | 5 | 5 | S | 130308 | Human mastadenovirus E |
| Yana_L5 | 0 | 2 | 2 | S | 108098 | Human mastadenovirus B |
| Yana_L5 | 0 | 2 | 0 | - | 413684 | unclassified Simian adenoviruses |
| Yana_L5 | 0 | 2 | 2 | S | 909210 | Simian adenovirus 18 |
| Yana_L1 | 0 | 1859 | 18 | G | 10509 | Mastadenovirus |
| Yana_L1 | 0 | 1824 | 1467 | S | 129951 | Human mastadenovirus C |
| Yana_L1 | 0 | 183 | 183 | - | 28285 | Human adenovirus 5 |
| Yana_L1 | 0 | 174 | 174 | - | 10533 | Human adenovirus 1 |
| Yana_L1 | 0 | 11 | 11 | S | 130310 | Human mastadenovirus D |

| | | | | | | |
|---|---|---|---|---|---|---|
| Yana_L1 | 0 | 6 | 0 | - | 413684 | unclassified Simian adenoviruses |
| Yana_L1 | 0 | 3 | 3 | S | 909210 | Simian adenovirus 18 |
| Yana_L1 | 0 | 3 | 0 | S | 1165908 | Simian adenovirus B |
| Yana_L1 | 0 | 3 | 3 | - | 995022 | Simian adenovirus 49 |
| Yana_L3 | 0 | 284 | 0 | G | 10509 | Mastadenovirus |
| Yana_L3 | 0 | 284 | 244 | S | 129951 | Human mastadenovirus C |
| Yana_L3 | 0 | 28 | 28 | - | 10533 | Human adenovirus 1 |
| Yana_L3 | 0 | 12 | 12 | - | 28285 | Human adenovirus 5 |
| Yana_L4 | 0 | 543 | 6 | G | 10509 | Mastadenovirus |
| Yana_L4 | 0 | 535 | 423 | S | 129951 | Human mastadenovirus C |
| Yana_L4 | 0 | 65 | 65 | - | 28285 | Human adenovirus 5 |
| Yana_L4 | 0 | 47 | 47 | - | 10533 | Human adenovirus 1 |
| Yana_L4 | 0 | 1 | 1 | S | 130310 | Human mastadenovirus D |
| Yana_L4 | 0 | 1 | 0 | - | 413684 | unclassified Simian adenoviruses |
| Yana_L4 | 0 | 1 | 1 | S | 909210 | Simian adenovirus 18 |
| Yana_L4 | 0 | 1 | 0 | G | 10552 | Aviadenovirus |
| Yana_L4 | 0 | 1 | 1 | S | 190063 | Fowl aviadenovirus C |
| Yana_L6 | 0 | 474 | 10 | G | 10509 | Mastadenovirus |

| | | | | | | |
|---|---|---|---|---|---|---|
| Yana_L6 | 0 | 456 | 382 | S | 129951 | Human mastadenovirus C |
| Yana_L6 | 0 | 51 | 51 | - | 10533 | Human adenovirus 1 |
| Yana_L6 | 0 | 23 | 23 | - | 28285 | Human adenovirus 5 |
| Yana_L6 | 0 | 6 | 0 | - | 413684 | unclassified Simian adenoviruses |
| Yana_L6 | 0 | 6 | 6 | S | 909210 | Simian adenovirus 18 |
| Yana_L6 | 0 | 2 | 2 | S | 130310 | Human mastadenovirus D |
| Yana2_L1 | 0 | 138 | 0 | G | 10509 | Mastadenovirus |
| Yana2_L1 | 0 | 137 | 91 | S | 129951 | Human mastadenovirus C |
| Yana2_L1 | 0 | 34 | 34 | - | 10533 | Human adenovirus 1 |
| Yana2_L1 | 0 | 12 | 12 | - | 28285 | Human adenovirus 5 |
| Yana2_L1 | 0 | 1 | 0 | S | 130310 | Human mastadenovirus D |
| Yana2_L1 | 0 | 1 | 1 | - | 651580 | Human adenovirus 54 |
| Yana2_L2 | 0 | 140 | 0 | G | 10509 | Mastadenovirus |
| Yana2_L2 | 0 | 139 | 85 | S | 129951 | Human mastadenovirus C |
| Yana2_L2 | 0 | 29 | 29 | - | 10533 | Human adenovirus 1 |
| Yana2_L2 | 0 | 25 | 25 | - | 28285 | Human adenovirus 5 |
| Yana2_L2 | 0 | 1 | 0 | S | 130369 | Porcine mastadenovirus A |

| | | | | | | |
|---|---|---|---|---|---|---|
| Yana2_L2 | 0 | 1 | 1 | - | 35265 | Porcine adenovirus 3 |
| Yana2_L3 | 0 | 136 | 0 | G | 10509 | Mastadenovirus |
| Yana2_L3 | 0 | 135 | 103 | S | 129951 | Human mastadenovirus C |
| Yana2_L3 | 0 | 22 | 22 | - | 10533 | Human adenovirus 1 |
| Yana2_L3 | 0 | 10 | 10 | - | 28285 | Human adenovirus 5 |
| Yana2_L3 | 0 | 1 | 1 | S | 130310 | Human mastadenovirus D |
| Yana2_L4 | 0 | 146 | 7 | G | 10509 | Mastadenovirus |
| Yana2_L4 | 0 | 130 | 90 | S | 129951 | Human mastadenovirus C |
| Yana2_L4 | 0 | 25 | 25 | - | 10533 | Human adenovirus 1 |
| Yana2_L4 | 0 | 15 | 15 | - | 28285 | Human adenovirus 5 |
| Yana2_L4 | 0 | 5 | 5 | S | 130310 | Human mastadenovirus D |
| Yana2_L4 | 0 | 4 | 0 | - | 413684 | unclassified Simian adenoviruses |
| Yana2_L4 | 0 | 4 | 4 | S | 909210 | Simian adenovirus 18 |
| Yana2_L5 | 0 | 137 | 0 | G | 10509 | Mastadenovirus |
| Yana2_L5 | 0 | 136 | 90 | S | 129951 | Human mastadenovirus C |
| Yana2_L5 | 0 | 32 | 32 | - | 10533 | Human adenovirus 1 |
| Yana2_L5 | 0 | 14 | 14 | - | 28285 | Human adenovirus 5 |
| Yana2_L5 | 0 | 1 | 1 | S | 130310 | Human mastadenovirus D |

**Table S2.** Kraken, metagenomic classification results for herpesvirus for each sample library.

| Sample | Percent of all reads classified | Number of reads classified on this taxonomic level and below | Number of reads classified on this taxonomic level | Taxonomic level | TaxId | Scientific name |
|---|---|---|---|---|---|---|
| Yana_L5 | 0 | 839 | 0 | O | 548681 | Herpesvirales |
| Yana_L5 | 0 | 836 | 0 | F | 10292 | Herpesviridae |
| Yana_L5 | 0 | 722 | 0 | - | 10357 | Betaherpesvirinae |
| Yana_L5 | 0 | 301 | 301 | S | 32604 | Human herpesvirus 6B |
| Yana_L5 | 0 | 50 | 50 | S | 10372 | Human herpesvirus 7 |
| Yana_L5 | 0 | 11 | 11 | S | 32603 | Human herpesvirus 6A |
| Yana_L5 | 0 | 295 | 295 | S | 10359 | Human herpesvirus 5 |
| Yana_L5 | 0 | 2 | 2 | S | 548914 | Elephant endotheliotropic herpesvirus 4 |
| Yana_L5 | 0 | 114 | 0 | - | 10293 | Alphaherpesvirinae |
| Yana_L5 | 0 | 107 | 107 | S | 10298 | Human herpesvirus 1 |
| Yana_L5 | 0 | 6 | 6 | S | 10325 | Macacine herpesvirus 1 |
| Yana_L5 | 0 | 3 | 0 | F | 548682 | Alloherpesviridae |
| Yana_L5 | 0 | 2 | 2 | S | 180230 | Cyprinid herpesvirus 3 |
| Yana_L5 | 0 | 1 | 1 | S | 150286 | Anguillid herpesvirus 1 |

| | | | | | | |
|---|---|---|---|---|---|---|
| Yana_L1 | 0 | 795 | 0 | O | 548681 | Herpesvirales |
| Yana_L1 | 0 | 775 | 0 | F | 10292 | Herpesviridae |
| Yana_L1 | 0 | 682 | 0 | - | 10357 | Betaherpesvirinae |
| Yana_L1 | 0 | 323 | 323 | S | 32604 | Human herpesvirus 6B |
| Yana_L1 | 0 | 55 | 55 | S | 10372 | Human herpesvirus 7 |
| Yana_L1 | 0 | 1 | 1 | S | 32603 | Human herpesvirus 6A |
| Yana_L1 | 0 | 217 | 217 | S | 10359 | Human herpesvirus 5 |
| Yana_L1 | 0 | 1 | 1 | S | 188763 | Panine herpesvirus 2 |
| Yana_L1 | 0 | 10 | 10 | S | 548914 | Elephant endotheliotropic herpesvirus 4 |
| Yana_L1 | 0 | 87 | 0 | - | 10293 | Alphaherpesvirinae |
| Yana_L1 | 0 | 47 | 47 | S | 10325 | Macacine herpesvirus 1 |
| Yana_L1 | 0 | 40 | 40 | S | 10298 | Human herpesvirus 1 |
| Yana_L1 | 0 | 6 | 0 | - | 10374 | Gammaherpesvirinae |
| Yana_L1 | 0 | 5 | 5 | S | 10376 | Human herpesvirus 4 |
| Yana_L1 | 0 | 1 | 1 | S | 35252 | Alcelaphine herpesvirus 1 |
| Yana_L1 | 0 | 20 | 0 | F | 548682 | Alloherpesviridae |
| Yana_L1 | 0 | 18 | 18 | S | 180230 | Cyprinid herpesvirus 3 |
| Yana_L1 | 0 | 1 | 1 | S | 317858 | Cyprinid herpesvirus 1 |

| | | | | | | |
|---|---|---|---|---|---|---|
| Yana_L1 | 0 | 1 | 1 | S | 317878 | Cyprinid herpesvirus 2 |
| Yana_L3 | 0 | 207 | 0 | O | 548681 | Herpesvirales |
| Yana_L3 | 0 | 180 | 0 | F | 10292 | Herpesviridae |
| Yana_L3 | 0 | 143 | 0 | - | 10357 | Betaherpesvirinae |
| Yana_L3 | 0 | 98 | 98 | S | 32604 | Human herpesvirus 6B |
| Yana_L3 | 0 | 24 | 24 | S | 10359 | Human herpesvirus 5 |
| Yana_L3 | 0 | 12 | 12 | S | 548914 | Elephant endotheliotropic herpesvirus 4 |
| Yana_L3 | 0 | 37 | 0 | - | 10293 | Alphaherpesvirinae |
| Yana_L3 | 0 | 18 | 18 | S | 10298 | Human herpesvirus 1 |
| Yana_L3 | 0 | 14 | 14 | S | 10325 | Macacine herpesvirus 1 |
| Yana_L3 | 0 | 5 | 5 | S | 35246 | Cercopithecine herpesvirus 9 |
| Yana_L3 | 0 | 27 | 0 | F | 548682 | Alloherpesviridae |
| Yana_L3 | 0 | 17 | 17 | S | 150286 | Anguillid herpesvirus 1 |
| Yana_L3 | 0 | 10 | 10 | S | 180230 | Cyprinid herpesvirus 3 |
| Yana_L4 | 0 | 281 | 0 | O | 548681 | Herpesvirales |
| Yana_L4 | 0 | 274 | 0 | F | 10292 | Herpesviridae |
| Yana_L4 | 0 | 203 | 0 | - | 10357 | Betaherpesvirinae |
| Yana_L4 | 0 | 98 | 98 | S | 32604 | Human herpesvirus 6B |

| | | | | | | |
|---|---|---|---|---|---|---|
| Yana_L4 | 0 | 16 | 16 | S | 10372 | Human herpesvirus 7 |
| Yana_L4 | 0 | 2 | 2 | S | 32603 | Human herpesvirus 6A |
| Yana_L4 | 0 | 65 | 65 | S | 10359 | Human herpesvirus 5 |
| Yana_L4 | 0 | 1 | 1 | S | 50290 | Aotine herpesvirus 1 |
| Yana_L4 | 0 | 1 | 0 | S | 50292 | Cercopithecine herpesvirus 5 |
| Yana_L4 | 0 | 1 | 1 | S | 188763 | Panine herpesvirus 2 |
| Yana_L4 | 0 | 5 | 5 | S | 548914 | Elephant endotheliotropic herpesvirus 4 |
| Yana_L4 | 0 | 67 | 0 | - | 10293 | Alphaherpesvirinae |
| Yana_L4 | 0 | 37 | 37 | S | 10325 | Macacine herpesvirus 1 |
| Yana_L4 | 0 | 25 | 25 | S | 10298 | Human herpesvirus 1 |
| Yana_L4 | 0 | 4 | 4 | S | 80341 | Equid herpesvirus 3 |
| Yana_L4 | 0 | 1 | 1 | S | 10320 | Bovine herpesvirus 1 |
| Yana_L4 | 0 | 4 | 0 | - | 10374 | Gammaherpesvirinae |
| Yana_L4 | 0 | 3 | 3 | S | 10371 | Equid herpesvirus 5 |
| Yana_L4 | 0 | 1 | 1 | S | 10398 | Ovine herpesvirus 2 |
| Yana_L4 | 0 | 7 | 0 | F | 548682 | Alloherpesviridae |
| Yana_L4 | 0 | 3 | 3 | S | 150286 | Anguillid herpesvirus 1 |
| Yana_L4 | 0 | 3 | 3 | S | 180230 | Cyprinid herpesvirus 3 |

| | | | | | | |
|---|---|---|---|---|---|---|
| Yana_L4 | 0 | 1 | 1 | S | 317878 | Cyprinid herpesvirus 2 |
| Yana_L6 | 0 | 250 | 0 | O | 548681 | Herpesvirales |
| Yana_L6 | 0 | 242 | 0 | F | 10292 | Herpesviridae |
| Yana_L6 | 0 | 215 | 0 | - | 10357 | Betaherpesvirinae |
| Yana_L6 | 0 | 116 | 116 | S | 32604 | Human herpesvirus 6B |
| Yana_L6 | 0 | 18 | 18 | S | 10372 | Human herpesvirus 7 |
| Yana_L6 | 0 | 4 | 4 | S | 32603 | Human herpesvirus 6A |
| Yana_L6 | 0 | 65 | 65 | S | 10359 | Human herpesvirus 5 |
| Yana_L6 | 0 | 1 | 1 | S | 548914 | Elephant endotheliotropic herpesvirus 4 |
| Yana_L6 | 0 | 27 | 0 | - | 10293 | Alphaherpesvirinae |
| Yana_L6 | 0 | 18 | 18 | S | 10298 | Human herpesvirus 1 |
| Yana_L6 | 0 | 8 | 8 | S | 10325 | Macacine herpesvirus 1 |
| Yana_L6 | 0 | 1 | 0 | - | 35247 | unclassified Alphaherpesvirinae |
| Yana_L6 | 0 | 1 | 1 | S | 332937 | Chimpanzee alpha-1 herpesvirus |
| Yana_L6 | 0 | 8 | 0 | F | 548682 | Alloherpesviridae |
| Yana_L6 | 0 | 8 | 8 | S | 180230 | Cyprinid herpesvirus 3 |

| | | | | | | |
|---|---|---|---|---|---|---|
| Yana2_L1 | 0 | 558 | 0 | O | 548681 | Herpesvirales |
| Yana2_L1 | 0 | 558 | 0 | F | 10292 | Herpesviridae |
| Yana2_L1 | 0 | 548 | 0 | - | 10357 | Betaherpesvirinae |
| Yana2_L1 | 0 | 483 | 483 | S | 10359 | Human herpesvirus 5 |
| Yana2_L1 | 0 | 30 | 30 | S | 32604 | Human herpesvirus 6B |
| Yana2_L1 | 0 | 27 | 27 | S | 10372 | Human herpesvirus 7 |
| Yana2_L1 | 0 | 1 | 1 | S | 32603 | Human herpesvirus 6A |
| Yana2_L1 | 0 | 2 | 2 | S | 548914 | Elephant endotheliotropic herpesvirus 4 |
| Yana2_L1 | 0 | 6 | 0 | - | 10374 | Gammaherpesvirinae |
| Yana2_L1 | 0 | 6 | 5 | S | 10376 | Human herpesvirus 4 |
| Yana2_L1 | 0 | 1 | 1 | - | 12509 | Human herpesvirus 4 type 2 |
| Yana2_L1 | 0 | 4 | 0 | - | 10293 | Alphaherpesvirinae |
| Yana2_L1 | 0 | 4 | 4 | S | 10298 | Human herpesvirus 1 |
| Yana2_L2 | 0 | 682 | 0 | O | 548681 | Herpesvirales |
| Yana2_L2 | 0 | 681 | 0 | F | 10292 | Herpesviridae |
| Yana2_L2 | 0 | 673 | 0 | - | 10357 | Betaherpesvirinae |
| Yana2_L2 | 0 | 583 | 583 | S | 10359 | Human herpesvirus 5 |
| Yana2_L2 | 0 | 55 | 55 | S | 32604 | Human herpesvirus 6B |

| | | | | | | |
|---|---|---|---|---|---|---|
| Yana2_L2 | 0 | 21 | 21 | S | 10372 | Human herpesvirus 7 |
| Yana2_L2 | 0 | 5 | 0 | - | 10374 | Gammaherpesvirinae |
| Yana2_L2 | 0 | 3 | 3 | S | 10376 | Human herpesvirus 4 |
| Yana2_L2 | 0 | 2 | 2 | S | 10371 | Equid herpesvirus 5 |
| Yana2_L2 | 0 | 3 | 0 | - | 10293 | Alphaherpesvirinae |
| Yana2_L2 | 0 | 2 | 2 | S | 80341 | Equid herpesvirus 3 |
| Yana2_L2 | 0 | 1 | 1 | S | 10298 | Human herpesvirus 1 |
| Yana2_L2 | 0 | 1 | 0 | F | 548682 | Alloherpesviridae |
| Yana2_L2 | 0 | 1 | 1 | S | 317858 | Cyprinid herpesvirus 1 |
| Yana2_L3 | 0 | 821 | 0 | O | 548681 | Herpesvirales |
| Yana2_L3 | 0 | 821 | 0 | F | 10292 | Herpesviridae |
| Yana2_L3 | 0 | 798 | 0 | - | 10357 | Betaherpesvirinae |
| Yana2_L3 | 0 | 710 | 710 | S | 10359 | Human herpesvirus 5 |
| Yana2_L3 | 0 | 1 | 1 | S | 188763 | Panine herpesvirus 2 |
| Yana2_L3 | 0 | 50 | 50 | S | 32604 | Human herpesvirus 6B |
| Yana2_L3 | 0 | 28 | 28 | S | 10372 | Human herpesvirus 7 |
| Yana2_L3 | 0 | 15 | 0 | - | 10293 | Alphaherpesvirinae |
| Yana2_L3 | 0 | 6 | 6 | S | 10298 | Human herpesvirus 1 |

| | | | | | | |
|---|---|---|---|---|---|---|
| Yana2_L3 | 0 | 7 | 7 | S | 80341 | Equid herpesvirus 3 |
| Yana2_L3 | 0 | 8 | 0 | - | 10374 | Gammaherpesvirinae |
| Yana2_L3 | 0 | 8 | 8 | S | 10376 | Human herpesvirus 4 |
| Yana2_L4 | 0 | 704 | 0 | O | 548681 | Herpesvirales |
| Yana2_L4 | 0 | 700 | 0 | F | 10292 | Herpesviridae |
| Yana2_L4 | 0 | 692 | 0 | - | 10357 | Betaherpesvirinae |
| Yana2_L4 | 0 | 591 | 591 | S | 10359 | Human herpesvirus 5 |
| Yana2_L4 | 0 | 6 | 6 | S | 188763 | Panine herpesvirus 2 |
| Yana2_L4 | 0 | 48 | 48 | S | 32604 | Human herpesvirus 6B |
| Yana2_L4 | 0 | 27 | 27 | S | 10372 | Human herpesvirus 7 |
| Yana2_L4 | 0 | 1 | 1 | S | 32603 | Human herpesvirus 6A |
| Yana2_L4 | 0 | 1 | 1 | S | 548914 | Elephant endotheliotropic herpesvirus 4 |
| Yana2_L4 | 0 | 6 | 0 | - | 10293 | Alphaherpesvirinae |
| Yana2_L4 | 0 | 4 | 4 | S | 10298 | Human herpesvirus 1 |
| Yana2_L4 | 0 | 2 | 2 | S | 80341 | Equid herpesvirus 3 |
| Yana2_L4 | 0 | 2 | 0 | - | 10374 | Gammaherpesvirinae |
| Yana2_L4 | 0 | 2 | 2 | S | 10376 | Human herpesvirus 4 |
| Yana2_L4 | 0 | 4 | 0 | F | 548682 | Alloherpesviridae |

| | | | | | | |
|---|---|---|---|---|---|---|
| Yana2_L4 | 0 | 4 | 4 | S | 180230 | Cyprinid herpesvirus 3 |
| Yana2_L5 | 0 | 628 | 0 | O | 548681 | Herpesvirales |
| Yana2_L5 | 0 | 626 | 0 | F | 10292 | Herpesviridae |
| Yana2_L5 | 0 | 613 | 0 | - | 10357 | Betaherpesvirinae |
| Yana2_L5 | 0 | 545 | 545 | S | 10359 | Human herpesvirus 5 |
| Yana2_L5 | 0 | 35 | 35 | S | 32604 | Human herpesvirus 6B |
| Yana2_L5 | 0 | 22 | 22 | S | 10372 | Human herpesvirus 7 |
| Yana2_L5 | 0 | 2 | 2 | S | 548914 | Elephant endotheliotropic herpesvirus 4 |
| Yana2_L5 | 0 | 7 | 0 | - | 10293 | Alphaherpesvirinae |
| Yana2_L5 | 0 | 4 | 4 | S | 10298 | Human herpesvirus 1 |
| Yana2_L5 | 0 | 3 | 3 | S | 80341 | Equid herpesvirus 3 |
| Yana2_L5 | 0 | 6 | 0 | - | 10374 | Gammaherpesvirinae |
| Yana2_L5 | 0 | 6 | 6 | S | 10376 | Human herpesvirus 4 |
| Yana2_L5 | 0 | 2 | 0 | F | 548682 | Alloherpesviridae |
| Yana2_L5 | 0 | 2 | 2 | S | 180230 | Cyprinid herpesvirus 3 |

**Table S3.** Sequencing library mapping statistics for herpesvirus reference genomes.

**Table S4.** Genomic coverage for herpesvirus reference genomes.

**Table S5.** bactdate results, run with seven million iterations

| | Including ancient sample | Without ancient samples |
|---|---|---|
| Probability of root branch | 0.37 | 0.33 |
| likelihood | -1.62e+02 [-1.71e+02;-1.54e+02] | -1.65e+02 [-1.74e+02;-1.57e+02] |
| prior | -1.15e+03 [-1.26e+03;-1.05e+03] | -7.56e+02 [-8.26e+02;-6.84e+02] |
| mu | 8.64e-05 [1.53e-05;2.07e-04] | 6.23e-03 [2.34e-03;1.42e-02] |
| sigma | 1.52e-04 [2.66e-05;3.96e-04] | 1.14e-02 [3.98e-03;2.80e-02] |
| alpha | 2.92e+05 [7.57e+04;8.80e+05] | 3.27e+03 [1.21e+03;6.93e+03] |
| Root date | -724811.92 [-2583670.66;-150208.10] | -5280.70 [-11801.88;-371.53] |

**Table S6.** Marginal likelihood of demographic models from BEAST2, after pathsampler with 35 steps, MCMC chain length of 7 million and 50% burnin.

| | Marginal likelihood | |
|---|---|---|
| Demographic model | 880 million iterations | 1.9 billion iterations |
| Exponential, relaxed log normal clock | -3797282 | -3796619 |
| Bayesian Skyline, relaxed exponential clock | -3796769 | -3797527 |
| Exponential, strict clock | -3796383 | -3796931 |
| Constant, strict clock | -3796361 | - |
| Bayesian Skyline, strict clock | -3796240 | -3796223 |

**Table S7.** Stats from BEAST2, Coalescent Bayesian Skyline strict clock
